# Elongasome core proteins and class A PBP1a display zonal, processive movement at the midcell of *Streptococcus pneumoniae*

**DOI:** 10.1101/2024.01.10.575112

**Authors:** Amilcar J. Perez, Melissa M. Lamanna, Kevin E. Bruce, Marc A. Touraev, Julia E. Page, Sidney L. Shaw, Ho-Ching Tiffany Tsui, Malcolm E. Winkler

**Affiliations:** Department of Biology, Indiana University Bloomington, Bloomington, IN 47405; Department of Microbiology, Blavatnik Institute, Harvard Medical School, Boston, MA 02115

**Author notes:** Corresponding author: Malcolm E. Winkler, Department of Biology, Indiana University Bloomington, 1001 East Third Street Bloomington, Indiana 47405 USA. A.J.P., M.M.L. and K.E.B. contributed equally to this work.

**Keywords:** processive movement of PG synthesis proteins, PBP2b:RodA:MreC elongasome dynamics, Class A PBP circumferential movement, MpgA muramidase confined subdiffusion, diffusion of non-active PG synthesis proteins

## Abstract

Ovoid-shaped bacteria, such as *Streptococcus pneumoniae* (pneumococcus), have two spatially separated peptidoglycan (PG) synthase nanomachines that locate zonally to the midcell of dividing cells. The septal PG synthase bPBP2x:FtsW closes the septum of dividing pneumococcal cells, whereas the elongasome located on the outer edge of the septal annulus synthesizes peripheral PG outward. We showed previously by sm-TIRFm that the septal PG synthase moves circumferentially at midcell, driven by PG synthesis and not by FtsZ treadmilling. The pneumococcal elongasome consists of the PG synthase bPBP2b:RodA, regulators MreC, MreD, and RodZ, but not MreB, and genetically associated proteins Class A aPBP1a and muramidase MpgA. Given its zonal location separate from FtsZ, it was of considerable interest to determine the dynamics of proteins in the pneumococcal elongasome. We found that bPBP2b, RodA, and MreC move circumferentially with the same velocities and durations at midcell, driven by PG synthesis. However, outside of the midcell zone, the majority of these elongasome proteins move diffusively over the entire surface of cells. Depletion of MreC resulted in loss of circumferential movement of bPBP2b, and bPBP2b and RodA require each other for localization and circumferential movement. Notably, a fraction of aPBP1a molecules also moved circumferentially at midcell with velocities similar to those of components of the core elongasome, but for shorter durations. Other aPBP1a molecules were static at midcell or diffusing over cell bodies. Last, MpgA displayed non-processive, subdiffusive motion that was largely confined to the midcell region and less frequently detected over the cell body.

**SIGNIFICANCE:** This paper reports three types of single-molecule motions of PG synthesis proteins in the ovoid-shaped, pathogenic bacterium *Streptococcus pneumoniae*, not reported previously in other bacteria. The core elongasome exhibits zonal, circumferential motion in the absence of MreB filaments, independent of FtsZ treadmilling or the processive movement of the septal PG synthase. Class A aPBP1a also moves processively at midcell, but is not a persistent component of the core elongasome. These types of motions have implications for the functions of these PG synthases and indicate that processive motion in pneumococcus follows spatially separate tracks, possibly reflective of PG structure. In contrast, the MpgA muramidase displays a different kind of subdiffusive motion that is largely confined to midcell by an unknown mechanism.

## INTRODUCTION

The peptidoglycan (PG) cell wall protects bacteria from osmotic stress, determines cell shape, size, and chaining critical for adaptation and host interactions, and serves as a scaffolding in Gram-positive bacteria for the attachment of wall-teichoic acids, capsules, and extracellular enzymes and virulence factors (1–5). PG synthesis has provided targets for a large number of clinically relevant antibiotics and remains one of the most fertile sources for the discovery of new antibiotic targets in drug-resistant pathogens (6–11). Although a general picture of PG synthesis has emerged (12–16), many fundamental questions remain unanswered about the composition, coordination, chronology, and regulation of the nanomachines that carry out PG synthesis in bacteria. Moreover, major differences have recently emerged in how different bacteria carry out PG synthesis and use homologous proteins in different ways (see (17–23)). *Streptococcus pneumoniae* (pneumococcus; *Spn*) has emerged as a model for PG synthesis in ovoid-shaped bacteria that is tractable to genetic, biochemical, and cell biological approaches (reviewed in (4, 24, 25)). *S. pneumoniae* is a human nasopharyngeal commensal bacterium that becomes a serious opportunistic pathogen, killing over one million people annually worldwide (26–28). Moreover, *S. pneumoniae* is an antibiotic-resistant “superbug,” for which new antibiotic targets are urgently needed (10, 29–31).

Unlike in rod-shaped bacteria, all PG synthesis is zonal and confined to a band at the middle of dividing *S. pneumoniae* cells (Fig. S1) (24, 32, 33). At the start of cell division, the septal and elongation (peripheral) PG synthesis nanomachines locate to an FtsZ ring at the equators of predivisional daughter cells (Fig. S1*A*) (34, 35). Unlike *Bacillus subtilis* and *Staphylococcus aureus* (36–39), septal ring closure and cell separation occur simultaneously during the *S. pneumoniae* cell cycle, as do septal and elongation PG synthesis, with elongation PG synthesis likely starting slightly before septal PG synthesis in the initial FtsZ ring (24, 32, 33, 35, 40). As septation progresses, the septal and elongation PG synthesis nanomachines physically separate to the inside and outside edges, respectively, of the midcell annular disk (Fig. S1*B* and S1*C*) (24, 33, 41). Septal PG synthesis in the inner ring, which contains FtsZ and the Class B bPBP2x:FtsW (SEDS) complex, separates the dividing cells (24, 32–35). Elongation PG synthesis, which is carried out by the pneumococcal core elongasome in the outer ring (24, 32, 33), emanates outward and has been postulated to push MapZ-nascent FtsZ/FtsA/EzrA rings toward the equators of new daughter cells (Fig. S1*A*) (34, 42, 43). Notably, *S. pneumoniae* lacks Min and Noc systems (44, 45) and forms FtsZ rings over the nucleoid (22, 46, 47).

Consequently, dividing pneumococcal cells characteristically contain three FtsZ rings throughout most of division, one at the closing septum and two in the developing equatorial rings (Fig. S1*A*) (34, 46, 48). Interestingly, proteins that mediate bundling FtsZ filaments, such as SepF, ZapA, and ZapJ (ZipA is absent), arrive to the equatorial rings very late in cell division (34, 41, 49).

Septal ring closure is zonal and confined to a midcell ring in many bacteria (50–52). Dynamics studies showed that the pneumococcal bPBP2x:FtsW PG septal synthase moves circumferentially at midcell driven by PG synthesis itself, and not by FtsZ treadmilling (34), as was initially concluded for *B. subtilis* (53). Recent papers demonstrate that zonal septal closure is indeed driven by circumferential septal PG synthesis in *Escherichia coli* (54), *S. aureus* (55), and *B. subtilis* (56), rather than by FtsZ treadmilling.

However, elongation PG synthesis in rod-shaped cells is not zonal but is distributed along the curved cylindrical body of growing cells (57–59). This synthesis is carried out by the Rod-complex elongasome consisting of MreB and the PG elongasome complex. Actin-like MreB polymerizes into multiple short, curved filaments perpendicular to the cell long axis along the cell membrane, which is the region of maximal negative Gaussian curvature (16, 60–62). MreB interacts with the cytoplasmic helix-turn-helix domain of the bitopic RodZ protein that also interacts with the elongasome PG synthase complex facing outside the cell membrane (62–65). The core elongasome consists of the polytopic shape, elongation, division and sporulation (SEDS) family protein RodA (glycosyltransferase; GT), an essential bitopic Class B bPBP (transpeptidase; TP), and bitopic and polytopic positive regulators MreC and MreD, respectively (16, 66–70). Similar to septal PG synthesis, movement of the assembled Rod elongasome is driven by side-wall PG synthesis itself, rather than by ATPase-dependent treadmilling of MreB (57–59). Thus, MreB filaments serve as curvature-sensing “rudders” for processive synthesis of separate parallel strands of side-wall PG (61). In addition, Class A PBPs, which are bitopic proteins with extracellular GT and TP activities, move largely in non-ordered paths and often stop, possibly filling in or reinforcing gaps left by the Rod system (16, 71–74).

In contrast to rod-shaped bacteria, pneumococcal elongasome PG synthesis is zonal and confined to the initial FtsZ ring and later to the outer edge of the septal annular disk (24, 32, 33). The core pneumococcal elongasome contains RodA, Class B bPBP2b, RodZ, MreC, and MreD, but does not contain an MreB homolog (19, 25, 35, 46, 75–79). RodZ(*Spn*) serves as an organizer of elongasome assembly and is required for MreC midcell localization followed by localization of RodA and bPBP2b (19, 80). Immunofluorescence microscopy (IFM) indicated that bPBP2b, MreC, and RodZ co-localize throughout the cell cycle at midcell and localize in middle-to-late divisional cells to a larger ring than FtsZ and bPBP2x, the TP in septal PG synthesis (19, 35, 46). This larger ring corresponds to the outer elongasome PG synthesis ring observed at midcell (Fig. S1). Direct evidence for this conclusion came from examination of FDAA-labeled vertically oriented cells by high-resolution 3D-SIM (32). bPBP2x and bPBP2b engaged in PG synthesis localized initially with FtsZ to the equatorial ring of pre-divisional cells. Later in division, bPBP2x and FtsZ located primarily to the inner FDAA-labeled ring at midcell, while bPBP2b located to the outer ring (32). Given that the synthetically active elongasome complex contains a complex of bPBP2b, RodA, MreC complex in *S. pneumoniae* (16, 19), it is presumed here that active RodA at midcell locates to the FtsZ ring early in division and to the outer midcell ring later in division, similar to bPBP2b and MreC. IFM also indicated that Class A aPBP1a and the MpgA muramidase co-localize with bPBP2b and MreC, but not with bPBP2x, throughout the cell cycle, including to the outer midcell ring later in division (Fig. S1) (35, 46, 80). In addition, interactions were detected between aPBP1a and MpgA and components of the elongasome, including RodZ and MreC (19). Thus, aPBP1a and MpgA localize with peripheral PG synthesis, although it has not been determined whether they are persistent members of the core elongasome.

Paradoxically, the absence of aPBP1a suppresses the requirement for MreCD and RodZ (80, 81), whereas inactivation of *pbp1b*, which encodes Class A aPBP1b, suppresses the requirement for RodZ, but not for MreCD (19). In both cases, the bPBP2b:RodA PG synthase is still required for viability, suggesting that the absence of aPBP1a or aPBP1b may activate bypass pathways that obviate the requirement for certain regulators of the core elongasome (19, 80). However, amino acid changes that reduce the catalytic activity of the MpgA muramidase (80, 82) abolish the requirement for the entire pneumococcal PG elongasome, including the bPBP2b:RodA PG synthase (24, 80). The CozE protein was also implicated in pneumococcal elongation PG synthesis (83). However, CozE did not behave equivalently to MreCD in transformation assays of function (84), and the exact function of pneumococcal CozE remains unknown. A pneumococcal TseB homolog renamed as CopD was recently reported to bind to bPBP2b and aPBP1a. Absence of CopD produces wider cells, consistent with a defect in elongation PG synthesis (85); however, the mechanism underlying this morphology change is not known. Last, FtsX is located with the outer ring elongasome proteins later in pneumococcal division (32). FtsX is a subunit of the FtsEX:PcsB PG hydrolase that is thought to play an essential role in PG remodeling between septal and peripheral PG (86–89). Notably, the FtsX(*Bsu*) homolog is also associated with side-wall PG synthesis (90, 91).

In this paper, we show that the pneumococcal bPBP2b, RodA, and MreC core elongasome proteins move circumferentially with the same velocity and processivity at midcell at different division stages. Unless indicated otherwise, midcell refers to the central rings at the septa of dividing cells and to the equators of predivisional daughter cells about to start division. The processive movement of the core elongasome components is driven by PG synthesis, and is not affected by FtsZ treadmilling, consistent with the absence of FtsZ in the outer ring of elongasome PG synthesis later in division. Strikingly, at WT protein expression levels in exponentially growing cells, a large majority of bPBP2b, RodA, and MreC molecules diffuse in the membrane over the body of pneumococcal cells without synthesizing PG. Unexpectedly, Class A aPBP1a also moves circumferentially at midcell with a similar velocity, but in shorter tracks, than the core elongasome components. Processive movement of aPBP1a at midcell depended on PG synthesis, and again, a majority of aPBP1a molecules were found diffusing over the cell body without synthesizing PG. Finally, the MpgA muramidase showed a distinctive pattern of non-processive, subdiffusive movement, largely confined to midcell regions. These results demonstrate that the pneumococcal PG elongasome moves zonally, which is different from elongasome movement in rod-shaped bacteria, that a Class A PBP moves circumferentially as well as diffusively, and that a PG hydrolase shows unusual fast non-processive movement confined largely to the midcell region. The implications of these modes of movement to the functions of the pneumococcal PG elongasome, Class A PBPs, and PG muramidases and to patterns of PG-glycan strand placement are discussed.

## RESULTS

### Construction of functional fluorescent-protein fusions

Fusions to pneumococcal elongasome proteins bPBP2b, RodA, MreC, aPBP1a, and MpgA, were constructed at native chromosomal loci to contain an N-terminal expression-enhancing “i-tag”, fused to an intracellular HaloTag (HT) domain, followed by a 15 amino-acid linker, fused to the protein of interest (Tables S1 and S2), as described previously (32, 34, 92–95). Growth of strains expressing iHT-fused proteins in C+Y, pH 6.9 medium was indistinguishable from that of WT (Fig. S2*A*). Localization of iHT-fused proteins labeled with saturating amounts of HT-TMR ligand matched previously published localization patterns of epitope-tagged or GFP-fused proteins imaged by IFM or two-dimensional epifluorescence microscopy (2D-FM), respectively (Fig. S2*B*) (19, 32, 34, 35, 42, 46, 80, 81, 85, 96). Western blots of these fusions with intracellular iHT domains probed with antibody to native MreC, bPBP2b, aPBP1a, or MpgA showed expression of full-length fusions with minimal cleavage of the iHT domain (Fig. S3*A*, S4*B*, S5*B*, and S6). In contrast, western blots of an extracellular C-terminal HT fusion to MreC (MreC-HT) probed with anti-MreC antibody showed extensive cleavage of the HT domain not detectable in blots probed with anti-HT antibody (Fig. S3*A*, middle and right). Cleavage of extracellular HT domains was also reported in *E. coli* (97), and our results emphasize the need to validate protein fusion stability in blots with antibody to native proteins.

We reported previously that some tagged proteins, such as iHT-bPBP2b and sfGFP-bPBP2b, were expressed from their chromosomal locus in significantly lower amounts than WT bPBP2b in cells grown in BHI broth, with only minor effects on cell growth and morphology under this growth condition (32). We observed similar results for cells grown in C+Y, pH 6.9 medium, which is utilized in live-cell fluorescence microscopy due to low levels of autofluorescence (34). The cellular amounts of iHT-bPBP2b and iHT-aPBP1a were 15 ± 5% and 10 ± 10% (mean ±SD) of the WT bPBP2b and aPBP1a amounts, respectively (Fig. S4*B* and S5*B*; Table S3). Underproduced iHT-bPBP2b caused minor increases in cell width and size (Fig. S4*C* and S4*D*). However, the reduced amount of iHT-aPBP1a caused decreased cell width, which phenocopied a Δ*pbp1a* mutant (Fig. S5*C* and S5*D*) (81). The cellular amounts of iHT-MreC and iHT-MpgA expressed from their chromosomal loci were 82 ±18% and 171 ± 11% of the WT MreC and MpgA amounts, respectively (Fig. S3*A* and S6; Table S3). Cells expressing iHT-MreC showed minimal cell morphology differences from WT (Fig. S2*B* and S3*B*).

To increase cellular amounts of iHT-bPBP2b, iHT-aPBP1a, and iHT-RodA (antibody to native RodA was not available), we constructed merodiploid strains in which each iHT-tagged protein was also expressed from an ectopic site under the control of a zinc-inducible promoter (98). Addition of Zn inducer ([ZnCl_2_] indicated + 1/10 [MnSO_4_] to reduce zinc toxicity (41, 80, 99)) to cultures of these merodiploid strains did not affect growth (Fig. S4*A* and S5*A*). Zn addition complemented the low cellular amounts of iHT-bPBP2b and iHT-aPBP1a back up to 82 ± 6% and 101 ± 41% of the WT bPBP2b and aPBP1a amounts, respectively (Fig. S4*B* and S5*B*; Table S3). Complementation also alleviated minor cell shape defects detected in strains with reduced protein amounts (Fig. S4*C*, S4*D*, S5*C*, S5*D*, and S7). iHT-fused proteins localized strongly to midcell of Zn-induced merodiploid strains, but were also more dispersed over bodies of cells compared to haploid or uninduced merodiploid strains (Fig. S4*C* and S5*C*). Finally, merodiploid strains expressing iHT-bPBP2b or iHT-RodA showed minimal amounts of cleavage products when 0.2 mM or 0.25 mM Zn inducer was added to the growth medium (Fig. S4*B* and S7*B*). Effects of protein amounts on dynamics are described below.

### bPBP2b, RodA, and MreC display similar processive circumferential motion at midcell

Strains expressing functional iHT-fused proteins were labeled with limiting concentrations of HT-JF549 ligand (Table S4), and single-molecule total internal reflection fluorescence microscopy (sm-TIRFm) was performed for 180 s, as described previously (34) (*Materials and Methods*). *S. pneumoniae* division is highly regular with septal and equatorial FtsZ-rings forming perpendicular to the long axis at the geometric centers of ovoid-shaped cells (Fig. S*1*). Midcell regions were considered to be within 125 nm of geometric centers. Three criteria were used to determine different motion types. Processive circumferential motion was defined as a molecule moving in one direction for more than six consecutive frames (imaging rate = 1 frame per second (FPS)), with a linear velocity of ≥ 5 nm/s determined by kymograph analysis (Fig. 1*A*; *Materials and Methods*). Static molecules were defined as being visible for more than six consecutive frames with a velocity < 5 nm/s. Diffusively moving molecules were defined as moving, but not in a consistent direction, for 6 or more non-consecutive frames within a period of 90 s.

**Fig. 1.**
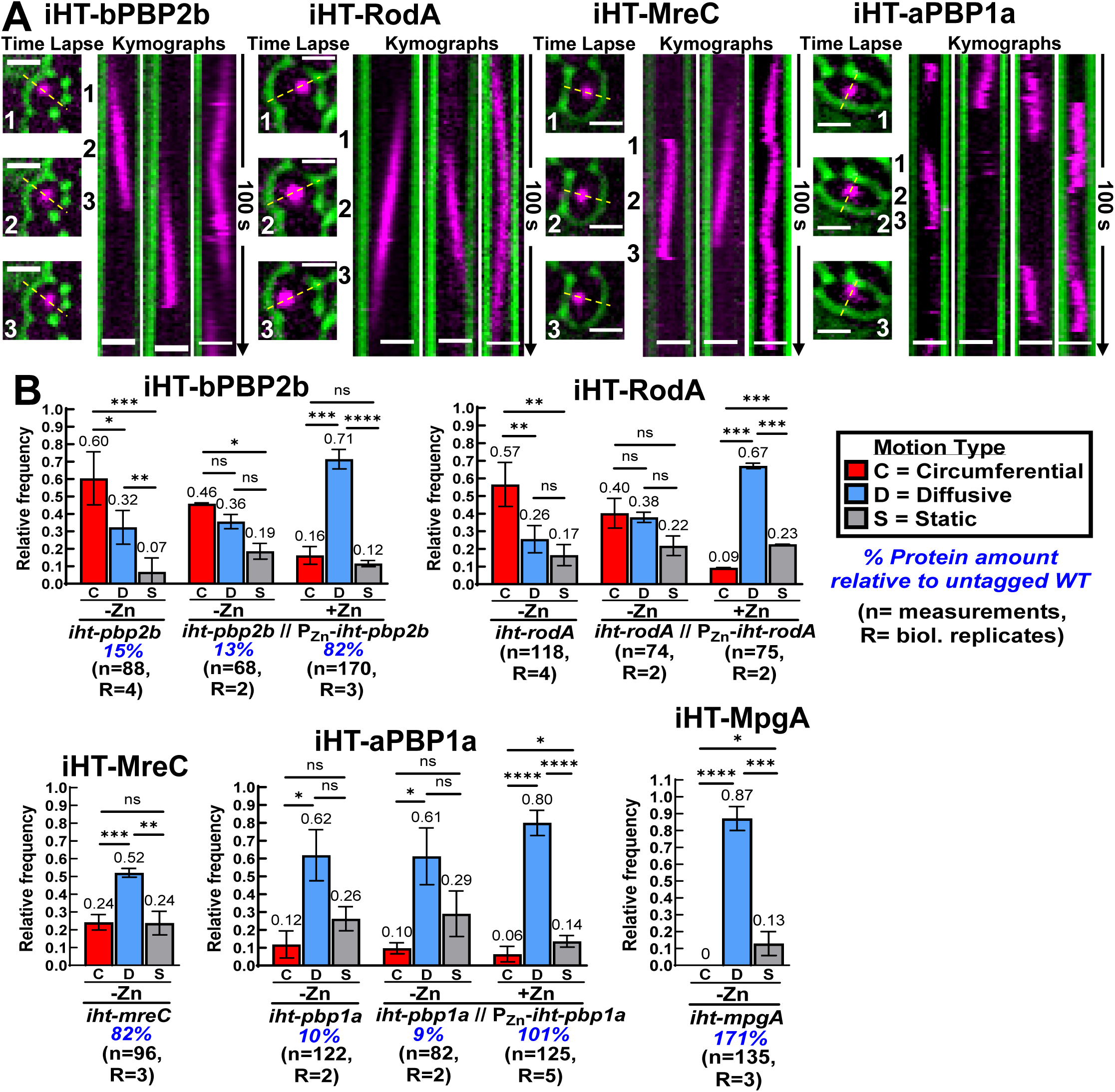
Elongation PG synthesis proteins displayed processive circumferential motion, and a limited number of molecules simultaneously engaged in PG synthesis. Sm-TIRFm was performed at 1 FPS as described in *Materials and Methods* on strains *iht-pbp2b* (IU15928), *iht-pbp2b* // P_Zn_*-iht-pbp2b* (IU16553), *iht-rodA* (IU15970), *iht-rodA* // P_Zn_*-iht-rodA* (IU16496), *iht-mreC* (IU16344), *iht-pbp1a* (IU16320), *iht-pbp1a* // P_Zn_*-iht-pbp1a* (IU16497), and *iht-mpgA* (IU15997). (*A*) Representative time-lapse images and kymographs of molecules displaying processive circumferential movement in strains IU15928, IU15970, IU16344, and IU16497. Cell outlines are DIC images pseudocolored green, and HT-labeled molecules are magenta. Location of lines used to make kymographs are shown in the time lapse images (yellow dashed lines). For each protein, the left kymograph was generated from the cell depicted in the time-lapse images. Numbers (1–3) denote when each image in the time lapse was taken. The other two kymographs are from cells that are not shown. Calculation of circumferential velocities from kymographs is described in *Materials and Methods*. Scale bars = 1 µm. (*B*) Movement patterns of HT-labeled sm molecules. Bars represent the mean relative frequency of each type of motion. For a given strain or condition, the relative frequencies of circumferential (red), diffusive (blue), and static (grey) molecules were determined manually. Frequencies were then averaged over all biological replicates to determine the mean (error bars represent ± SD). Mean values are indicated above each error bar, and the total number of molecules analyzed (n) from the number of biological replicates (R) for each strain is listed below the strain or condition. Circumferentially moving molecules were defined as molecules moving in one direction for 6 or more frames (at 1 FPS imaging rate) with a linear velocity ≥ 5 nm/s. Static molecules were defined as molecules not moving or moving for 6 or more frames with a velocity < 5 nm/s. Diffusive molecules were defined as molecules that moved, but not in a consistent direction, for 6 or more non-consecutive frames within a period of 90 s. Full criteria for determining movement patterns are described in *Materials and Methods*. The expression level of iHT-fusion proteins relative to the untagged WT level is shown as a percentage under the strain names. iHT-fusion proteins were expressed solely from native chromosomal loci or with an additional copy of the gene encoding the iHT-fusion protein at an ectopic site under control of a zinc-inducible promoter, with 0 or 0.25 mM Zn inducer added. Unpaired t-tests were performed to compare relative frequencies of motion types. *ns* (nonsignificant); \**P* < 0.05; \*\**P* < 0.01; \*\*\**P* < 0.001; \*\*\*\**P* < 0.0001.

Single molecules of iHT-bPBP2b, iHT-RodA, and iHT-MreC displayed similar processive circumferential motion exclusively at midcell (Fig. 1*A* and S8; Movies S1 to S3), that resembled the circumferential motion of iHT-bPBP2x and FtsW-HT reported previously (34). The frequency of circumferential, static, and diffusive molecules was compiled for each strain (Fig. 1*B*). Examination of the circumferential tracks showed that 73% (54/74) of iHT-bPBP2b and 74% (57/77) of iHT-MreC molecules moved unidirectionally at midcell. The remaining 27% of iHT-bPBP2b or 26% of iHT-MreC molecules moved in one direction for at least six consecutive frames and then reversed direction and moved in the opposite direction for at least six frames at approximately the same velocity (angle in kymograph; Fig. 1*A*, right). Slow or non-moving static molecules were mainly detected at midcell (Fig. S8), whereas diffusive molecules displayed rapid, erratic movement throughout the cell (Fig. 1*B*; S8; Movies S1 to S3). Fewer than 5% of molecules of these core elongasome proteins were observed to transition between movement states (e.g., static to diffusive or circumferential).

### A limited number of bPBP2b, RodA, and MreC molecules engage in active PG synthesis

The distribution of single-molecule movement states of iHT-bPBP2b and iHT-RodA was highly dependent on cellular protein levels. When expressed at 15% of the WT level, ≈60% of iHT-bPBP2b moved circumferentially and only 32% moved diffusively (Fig. 1*B*; Movie S1). The remaining 7% of iHT-bPBP2b molecules were static, mainly at midcell. iHT-RodA molecules expressed from the native locus showed a similar distribution, suggesting that iHT-RodA was underproduced (Fig. 1*B*; Movie S2). However, when iHT-PBP2b and iHT-RodA protein amounts were increased by ectopic expression (to ≈82% the WT level for iHT-bPBP2b), only 16% or 9% of single iHT-bPBP2b or iHT-RodA molecules, respectively, moved circumferentially, while ≈70% moved diffusively, with the remaining 12% iHT-bPBP2b or 23% iHT-RodA static, mostly at midcell (Fig. 1*B*; Movies S4 and S5). Uninduced merodiploid strains expressing iHT-bPBP2b or iHT-RodA displayed movement distributions similar to those of haploid strains (Fig. 1*B*). The amount iHT-MreC expressed from its native locus was nearly (82%) that of WT (Fig. S3*A*; Table S3). Approximately 24% of iHT-MreC molecules moved circumferentially, while 52% moved diffusively and 24% were static, similar to the movement distributions of iHT-bPBP2b and iHT-RodA when they were ectopically expressed (Fig. 1*B*; Movie S3).

High-resolution 3D structured-illumination microscopy (3D-SIM) on fixed cells corroborated conclusions based on movement distributions. At protein amounts corresponding to ≈15% of WT, iHT-bPBP2b localized primarily to regions of active PG synthesis at midcell and equators of dividing cells, as indicated by HADA fluorescent D-amino-acid labeling (Fig. 2*A*). When expressed near the WT level, iHT-bPBP2b remained at the HADA-labeled midcell and equators but was mostly dispersed over the body of the entire cell where HADA labeling was not detected (Fig. 2*B*). These results support the conclusion that iHT-PBP2b molecules that move diffusively in the cell membrane outside of midcell and equators are not actively synthesizing PG (Fig. 2*B*). Control experiments using a GFP fusion construct to bPBP2b and labeling with a different fluorescent D-amino acid (TADA) gave similar results (Fig. S9). 2D-FM images also showed an increase in dispersed iHT-bPBP2b localization when protein expression was increased to near the WT amount (Fig. S4*C*). We conclude that at any given time, a limited number of bPBP2b, RodA, and MreC molecules display processive circumferential motion at midcell where PG synthesis occurs. Conversely, bPBP2b, RodA, and MreC are expressed in excess in WT cells, and most molecules are diffusing over the cell surface and not synthesizing PG in cells under these and other growth conditions (Fig. 1*B,* 2*B,* and S4*C*; Table S3) (32).

**Fig. 2.**
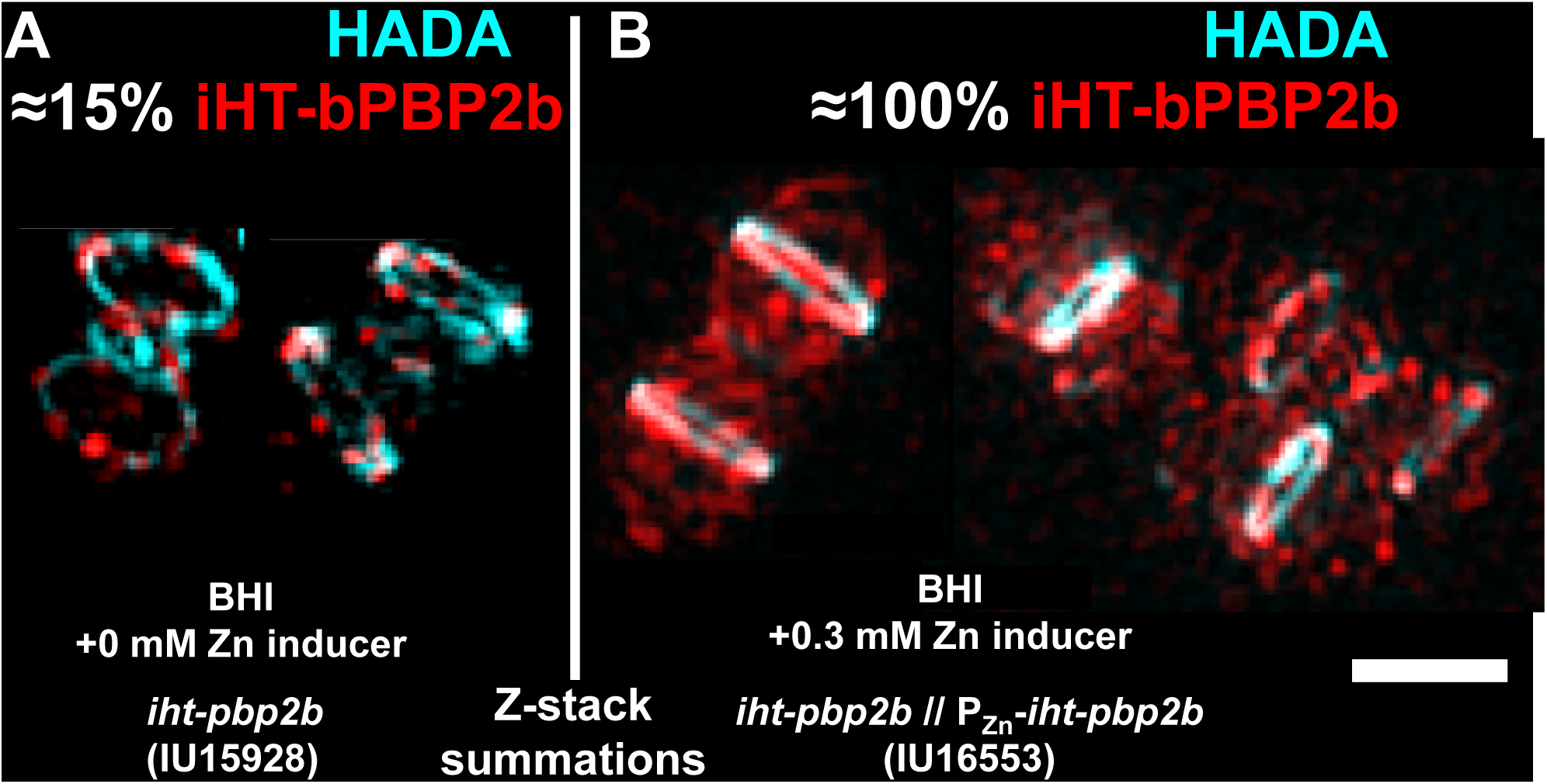
Diffusively moving components of the core PG elongasome did not actively synthesize PG (blue) in non-midcell regions of exponentially growing pneumococcal cells. 3D-SIM images, where iHT-bPBP2b is red, regions of PBP transpeptidase activity are blue, and midcell regions include septa of dividing cells and equators of predivisional daughter cells. Cells were grown in BHI ± 0.3 mM Zn inducer, labeled with a saturating amount of HT-ligand (500 nM) for 15 min, washed, labeled with 400 µM of the fluorescent D-amino acid HADA (blue) for 2.5 min, fixed, and imaged as described in *Materials and Methods*. Images are summed from 15 *Z*-plane sections (*Z*-stack) representing a total depth of 1.875 µm. Percentages indicate the amount of iHT-bPBP2b expressed relative to the untagged WT bPBP2b^+^ level (32). (*A*) *iht-pbp2b* (IU15928) and (*B*) *iht-pbp2b* // P_Zn_*-iht-pbp2b* (IU16553). Scale bar = 1 µm.

### bPBP2b, RodA, and MreC form a stable elongasome complex that undergoes circumferential motion

If bPBP2b, RodA, and MreC form an elongasome complex, we would expect that each molecule would move with the same velocity. To test this idea, velocities of circumferentially moving molecules were determined by kymograph analysis (Fig. 3*A*) (34). The length of time these molecules were observed moving circumferentially, referred to here as the circumferential duration, was also measured (Fig. 3*B*), and the particle distance each circumferential molecule moved was calculated by multiplying the velocity by the duration (Fig. S10*A*). When expressed near WT levels (+Zn in merodiploid and haploid *mreC* constructs) the mean circumferential velocities, durations, and particle distances of iHT-bPBP2b, iHT-RodA, and iHT-MreC were statistically the same, at ≈11 nm/s, ≈23 s, and ≈230 nm, respectively (Fig. 3*A*, 3*B*, and S10*A*). Additional strains containing different fusion constructs of bPBP2b or RodA moved at similar circumferential velocities (Fig. S11). These data support the idea that bPBP2b, RodA, and MreC form a stable, circumferentially moving elongasome complex at midcell.

**Fig. 3.**
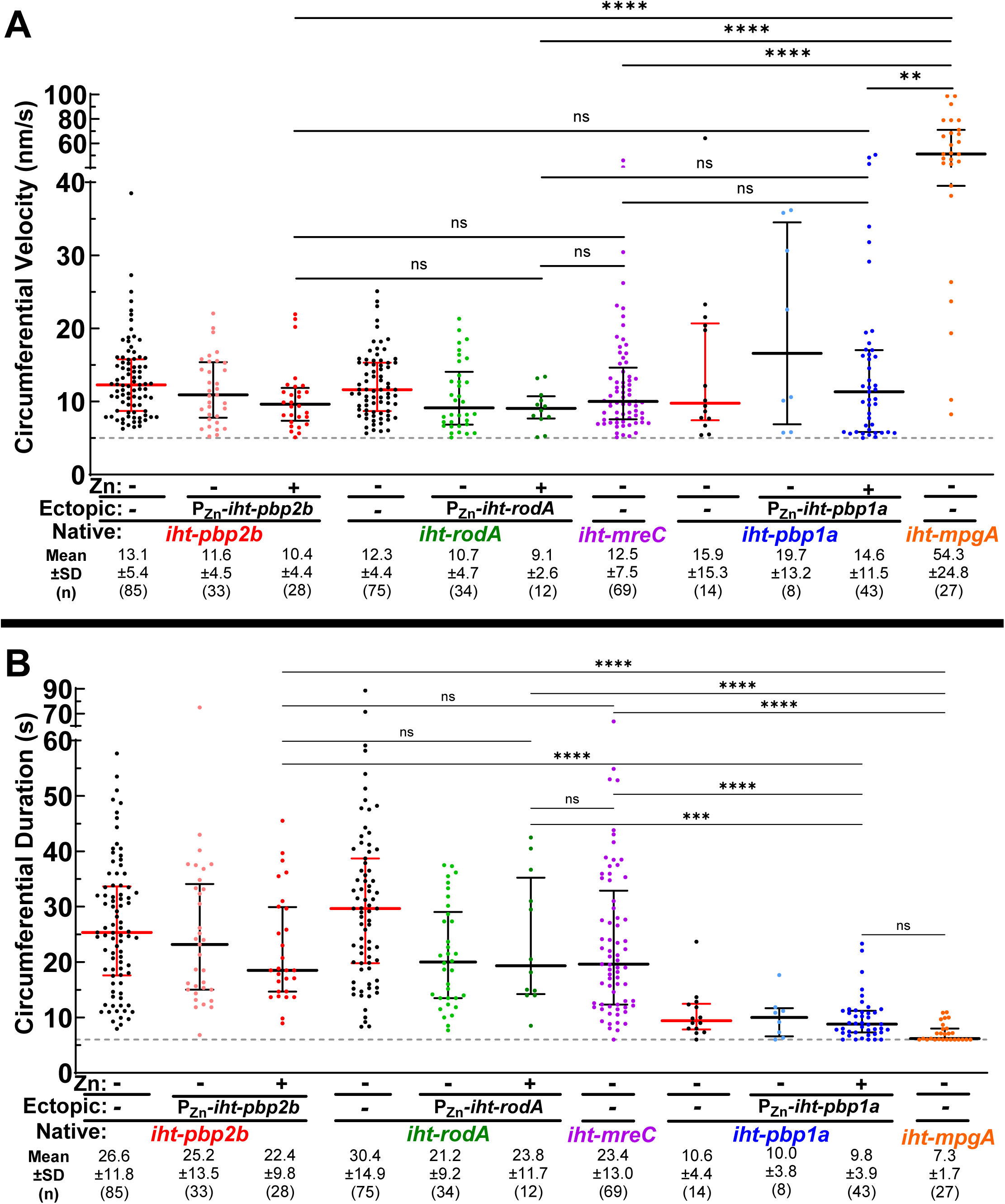
bPBP2b, RodA, and MreC formed a stable complex during active PG synthesis. Dot plots of (*A*) circumferential velocities and (*B*) circumferential durations of elongation PG synthesis proteins were determined by sm-TIRFm at 1 FPS. Strains are the same as in Figure. 1. +Zn indicates 0.25 mM Zn inducer was added. Black and red lines are median ± interquartile, and mean ± SD are indicated. n = total molecules analyzed from 2-5 biological replicates. Dotted grey lines indicate the minimum thresholds for (*A*) velocity (5 nm/s) and (*B*) duration (6 s). A Kruskal-Wallis with Dunn’s multiple comparisons test was used to compare circumferential velocities or circumferential durations in different strains. *ns* (nonsignificant); \*\**P* < 0.01; \*\*\**P* < 0.001; \*\*\*\**P* < 0.0001.

### Circumferential movement of bPBP2b and RodA is independent of FtsZ treadmilling and driven by PG synthesis

Circumferential movement of the pneumococcal septal PG synthase was shown to be independent of FtsZ treadmilling and driven by PG synthesis (34). A similar conclusion was later reported for the septal PG synthases of *S. aureus* (55) and *B. subtilis* (56) and for the population of active septal PG synthases in *E. coli* (54, 100). In early divisional cells, the pneumococcal PG elongasome is located in an FtsZ-ring that also contains the septal PG synthase (Fig. S1*A*), whereas at later stages of division, the elongasome separates from the FtsZ-ring and locates to the outer edge of the septal annular disk (Fig. S1*B* and S1*C*) (24, 32, 33). Notably, the same circumferential velocity of core elongasome protein iHT-MreC, which was expressed near the WT level (Fig. S3*A*), was detected at equators of early divisional cells, before ostensible constriction, and in later-divisional cells with clear constrictions, when septal (inner) and elongasome (outer) rings are separated at midcell (32) (Fig. S1*B* and S10*B*). Since the pneumococcal PG elongasome is initially in an FtsZ ring, we tested whether iHT-bPBP2b velocity is decreased when FtsZ treadmilling is greatly reduced by ectopic expression of GTPase mutant FtsZ(D214A), which reduces FtsZ treadmilling velocity from ≈32 nm/s to ≈4 nm/s (34). To the contrary, overproduction of FtsZ(D214A) caused a statistically minimal increase in iHT-bPBP2b velocity (Fig. 4*A*; Movie S6), with a marginal decrease in growth rate (Fig. S12). Consistent with a lack of dynamic coupling, the treadmilling velocity of FtsZ-sfGFP filaments was ≈2.4-fold greater than the circumferential velocity of iHT-bPBP2b single molecules expressed in the same cell (Fig. 4*B*; Movie S7). We conclude that circumferential movement of the pneumococcal PG elongasome is independent of FtsZ treadmilling.

**Fig. 4.**
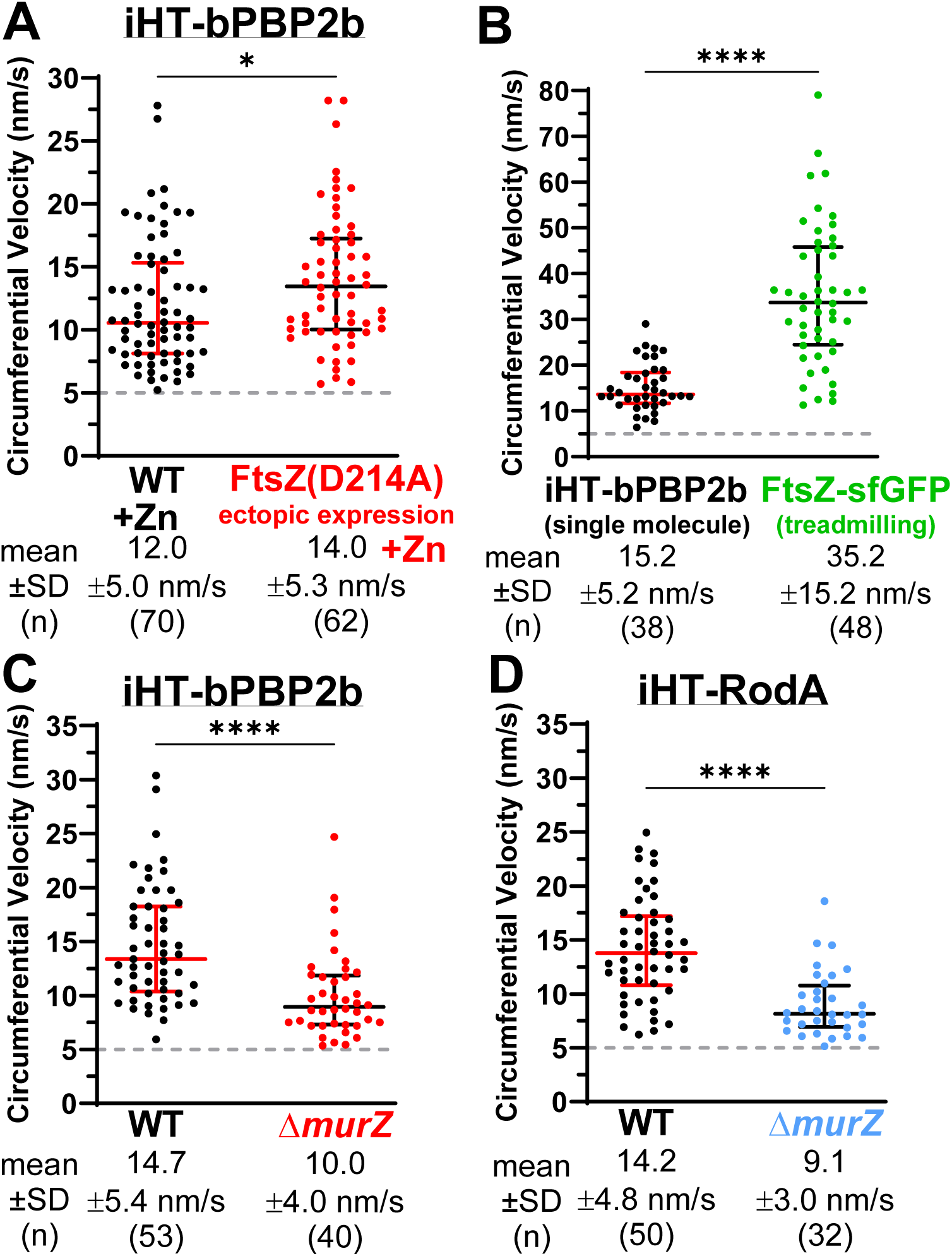
Circumferential movement of bPBP2b and RodA was independent of FtsZ treadmilling and reflective of PG synthesis. Sm-TIRFm was performed at 1 FPS on strains expressing iHT-bPBP2b or iHT-RodA as described in *Materials and Methods*. Dot plots of circumferential velocities are shown. Black and red lines are median ± interquartile, and mean ± SD are indicated. (*A*) Both *iht-pbp2b* (black, IU15928) and *iht-pbp2b* P_Zn_-*ftsZ*(D214A) (red, IU16091) strains were supplemented with 0.25 mM Zn inducer. (*B*) IU16056 expresses both iHT-bPBP2b and FtsZ-sfGFP. The treadmilling velocity of FtsZ-sfGFP filaments (green) and the circumferential velocity of iHT-bPBP2b single molecules (red) were determined. (*C*) *iht-pbp2b* (black, IU15928) and *iht-pbp2b* Δ*murZ* (red, IU16110) strains. (*D*) *iht-rodA* (black, IU15970) and *iht-rodA* Δ*murZ* (blue, IU16112) strains. n = total molecules analyzed from two biological replicates. Circumferential velocities in different strains were compared using a Mann-Whitney test. \**P* < 0.05; \*\*\*\**P* < 0.0001.

Two lines of evidence indicate that pneumococcal PG elongasome circumferential movement is driven by PG synthesis. First, we determined iHT-bPBP2b or iHT-RodA circumferential velocity in a Δ*murZ* (previously called Δ*murA1*) mutant, which lacks the predominant MurZ homolog that catalyzes the first committed step of PG precursor synthesis (34, 101). The circumferential velocity of iHT-bPBP2b or iHT-RodA was substantially reduced by ≈1.6-fold in the Δ*murZ* mutant compared to WT (Fig. 4*C* and 4*D;* Movies S8 and S9). Previously, we showed that the velocity of FtsZ treadmilling is unchanged in the Δ*murZ* mutant (34). Together, these results demonstrate that the rate of midcell elongasome movement is dependent on PG synthesis precursor amounts.

Second, catalytic mutants of bPBP2b TP or RodA GT activity did not exhibit processive, circumferential movement. No β-lactam antibiotic is known that preferentially inhibits bPBP2b alone (102). Both bPBP2b and RodA are essential in *S. pneumoniae* (35, 75, 103). Consequently, we constructed merodiploid strains with *iht-pbp2b*(S391A) (TP catalytic mutation) or *iht-rodA*(D283A) (GT catalytic mutation) (69) at chromosomal native sites and *pbp2b*^+^ or *rodA*^+^ under the control of the zinc-inducible promoter (P_Zn_) in an ectopic site (Fig. S13*A*). The merodiploid strains were constructed and grown in the presence of Zn inducer, which was removed to deplete the WT proteins (Fig. S13*B*). Interestingly, we were able to construct a *rodA*(D283A)//P_Zn_-*rodA*^+^ strain, but we were unable to construct a *pbp2b*(S391A)//P_Zn_-*pbp2b*^+^ strain in the presence of Zn inducer. This result suggested that *pbp2b*(S391A) was dominant-negative to WT *pbp2b*^+^. The reason we were able to construct the *iht-pbp2b*(S391A)//P_Zn_-*pbp2b*^+^ merodiploid was the low (≈15%) intrinsic expression of iHT-bPBP2b compared to WT bPBP2b (Fig. S4*B*). Consistent with a dominant-negative effect, depletion of WT bPBP2b in the *iht-pbp2b*(S391A)//P_Zn_-*pbp2b*^+^ merodiploid resulted in a more severe drop in OD_620_ than in Δ*pbp2b*//P_Zn_-*pbp2b*^+^ (Fig. S13*B*), possibly due to the toxicity of uncrosslinked PG produced from GT activity without TP activity (104). Western blot analysis confirmed that both iHT-bPBP2b(S391A) and iHT-RodA(D283A) were expressed and stable relative to iHT-bPBP2b^+^ and iHT-RodA^+^, respectively (Fig. S14*A*), and that bPBP2b^+^ expressed from the ectopic locus was fully depleted after 3h (Fig. S14*B*).

No midcell circumferential movement of iHT-bPBP2b(S391A) was detected in the presence or absence of WT bPBP2b (Fig. S13*C*; Movies S10 and S11). Instead, iHT-bPBP2b(S391A) was static at midcell (Fig. S13*D*) or was detected diffusing over the body of cells (Fig. S13*C*). Likewise, no midcell circumferential movement of iHT-RodA(D283A) was detected in the absence of RodA^+^ (Fig. S13*C*; Movie S12). Like iHT-bPBP2b(S391A), iHT-RodA(D283A) was static at midcell (Fig. S13*D*) or diffusing over the body of cells (Fig. S13*C*). Notably, both iHT-bPBP2b(S391A) and iHT-RodA(D283A) located to midcell (Fig. S13*E*), indicating that catalytic activity was not required for normal localization. From these combined results, we conclude that processive, circumferential movement of the pneumococcal elongasome PG synthase is driven by PG synthesis, but not by FtsZ treadmilling, even in predivisional cells.

### bPBP2b localization becomes diffuse and circumferential movement is lost when MreC is depleted

We reported previously that the pneumococcal PG elongasome is assembled sequentially (19). RodZ directs localization of MreC, which then directs localization of the bPBP2b:RodA synthase. In agreement with this conclusion, iHT-bPBP2b localization became diffuse when MreC was depleted (Fig. S15*A*). In addition, there was a large (≈30-fold) drop in the number of iHT-bPBP2b molecules moving circumferentially when MreC was depleted (Fig. S15*B*; Movie S13). Growth curve analysis confirmed reduced growth yield of cells depleted for MreC (Fig. S16*A*), and western blots confirmed the presence of iHT-bPBP2b during MreC depletion (Fig. S16*B* and S16*C*). We conclude that MreC is required for proper midcell localization and for elongasome PG synthesis by the bPBP2b:RodA synthase. This result is consistent with the previous conclusion that MreC allosterically activates the GT/TP activity of RodA:PBP2 in the PG elongasome of *E. coli* (16, 67, 105).

### bPBP2b and RodA require each other for localization

We further tested whether RodA was required for bPBP2b localization, and *vice versa*. Indeed, upon RodA depletion, iHT-bPBP2b went from locating primarily to midcell to being distributed at midcell and over the surface of rounded cells (Fig. S17*A* and S18*C*), indicative of loss of peripheral PG synthesis (35, 75, 81). Additionally, a growth yield phenotype was observed at 4 to 5 h into depletion (Fig. S18*A* and S18*B*). Upon bPBP2b depletion to ≈6% of WT after 3 h (Fig. S14*B*), iHT-RodA also relocated primarily from midcell to being distributed at midcell and over the surface of rounded cells, with a similar reduction in growth yield (Fig. S17*B* and S18). Therefore, there is an interdependence for normal localization of bPBP2b and RodA in the pneumococcal PG elongasome.

We next determined the dynamics of iHT-bPBP2b upon depletion of RodA or iHT-RodA upon depletion of bPBP2b. In both cases, the motion distribution shifted from primarily circumferential (≈59%) to diffusive (≈65%) movement (Fig. S17*C*; Movies S14 and S15), where iHT-bPBP2b and likely iHT-RodA were underproduced in these stains (Fig. 1*B*, S7, and S14*B*). Remaining circumferential movement at the midcell (≈24% of molecules) likely reflected bPBP2b:RodA complexes that assembled before depletion occurred. Consistent with this interpretation, the velocity of circumferentially moving proteins was unchanged by depletion of its cognate partner (Fig. S17*D* and S17*E*). These data support the idea that when bPBP2b or RodA is depleted, its unpaired cognate partner stops elongation PG synthesis at midcell and is released to diffuse over the cell body. Notably, released, unpaired iHT-bPBP2b or iHT-RodA did not move circumferentially at an increased velocity, suggesting that FtsZ filament end-tracking does not occur for these proteins (100).

### aPBP1a displays circumferential dynamics at midcell driven by PG synthesis but is not persistently part of the elongasome

Molecules of iHT-aPBP1a unexpectedly displayed circumferential, as well as diffusive and static dynamics (Fig. 1*A* and 1*B*; Movies S16 to S18). Like iHT-bPBP2b and likely iHT-RodA, iHT-aPBP1a was underproduced in the haploid strain and required ectopic expression to reach or surpass WT levels (Fig. 1*B* and S5*B*). However, in contrast to core elongasome components, the relative frequency of circumferentially moving iHT-aPBP1a molecules (≈10%) was largely independent of protein expression level (Fig. 1*B*), and a high percentage (>60%) of iHT-aPBP1a molecules moved diffusively, even at low expression levels (Fig. 1*B* and S5*B*; Table S3).

The mean circumferential velocity of iHT-aPBP1a expressed near WT levels was ≈15 nm/s, which was similar to, if not somewhat faster than, that of iHT-bPBP2b, iHT-RodA, and iHT-MreC (≈11 nm/s) (Fig. 1*A* and 3*A*). Strikingly, the mean circumferential duration of iHT-aPBP1a molecules was ≈10 s, more than two-fold shorter than that of iHT-bPBP2b, iHT-RodA, or iHT-MreC (≈23 s) (Fig. 1*A* and 3*B*). Altogether, these results indicate that circumferential motion of aPBP1a is distinct from that of bPBP2b, RodA, and MreC, suggesting that aPBP1a is not persistently part of the core elongasome complex, although shorter interactions are still possible.

To confirm that circumferential movement of iHT-aPBP1a molecules depends on PG synthesis, we constructed an *iht*-*pbp1a*(S370A)//P_Zn_-*iht*-*pbp1a*(S370A) merodiploid strain expressing catalytically inactive (TP null) iHT-aPBP1a(S370A) from both the native site and ectopic locus (Table S1). The growth rate of the catalytically inactive iHT-aPBP1a(S370A) merodiploid strain was reduced slightly (≈1.3-fold) when Zn inducer was added to the growth medium (Fig. S19*A*), but protein expression and localization were comparable to that of iHT-aPBP1a merodiploid strains (Fig. S19*B* and S19*C*).

Circumferential, diffusive, and static molecules were sorted by cellular location into midcell or locations other than midcell (non-midcell) (Fig. 5). Circumferential movement of iHT-aPBP1a decreased substantially from a relative frequency of ≈21% for iHT-aPBP1a to a marginal ≈3% for iHT-aPBP1a(S370A) at midcell, with the remainder of midcell iHT-aPBP1a or iHT-aPBP1a(S370A) molecules static (Fig. 5; Movie S19). Essentially no circumferentially moving molecules of iHT-aPBP1a or iHT-aPBP1a(S370A) were detected outside of midcell (Fig. 5). A similar conclusion was reached when data from Figure 1*B* was sorted into midcell or non-midcell locations (Fig. S20*A*). Since no PG synthesis was detectable by FDAA labeling away from the midcell (Fig. 2 and S9), and nearly all aPBP1a molecules away from midcell were moving diffusively (Fig. 5 and S20*A*), this further suggests that diffusively moving aPBP1a were not actively synthesizing PG in exponentially growing cells. Taken together, these data indicate that circumferentially moving iHT-aPBP1a molecules are actively synthesizing PG, while inactive molecules are either static at midcell or moving diffusively throughout the cell. A similar conclusion was reached for elongasome proteins iHT-bPBP2b, iHT-RodA, and iHT-MreC (Fig. 1*B* and 2).

**Fig. 5.**
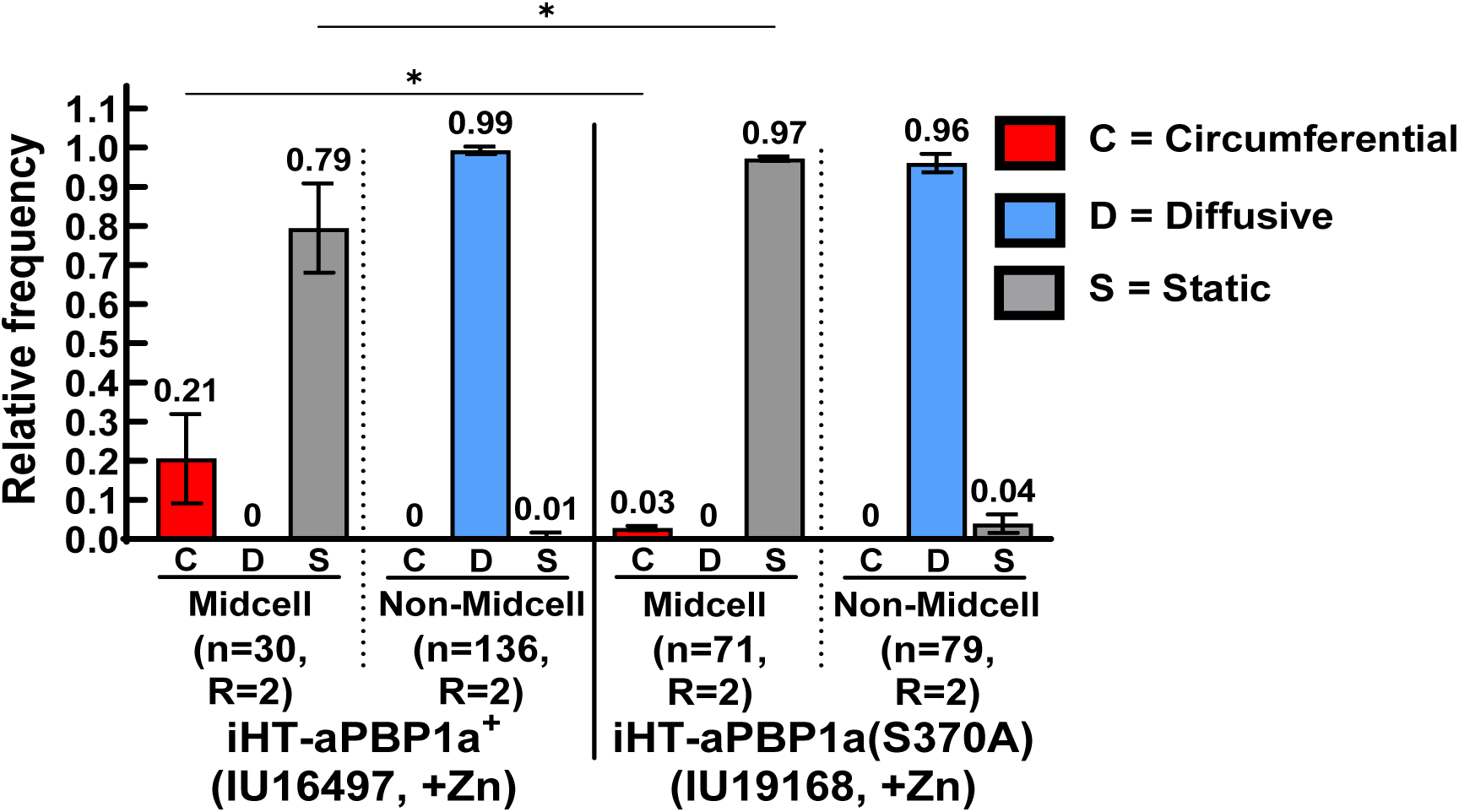
Circumferential movement of aPBP1a reflected active PG synthesis. Sm-TIRFm was performed at 1 FPS as described in *Materials and Methods* on strains *iht-pbp1a* // P_Zn_-*iht-pbp1a* (IU16497) and *iht-pbp1a*(S370A) // P_Zn_-*iht-pbp1a*(S370A) (IU19168) with 0.25 mM Zn inducer added. Movement patterns of HT-labeled molecules at sites of PG synthesis (midcell) or elsewhere in the cell (non-midcell) were determined. The layout is the same as Figure 1B (see legend for details). A two-way ANOVA with Tukey’s multiple comparison test was used to compare relative frequencies of motion types among strains. \**P* < 0.05.

Other data were consistent with aPBP1a not being a constant component of the PG elongasome. Notably, the velocity of iHT-aPBP1a in a merodiploid *iht-pbp1a*//P_Zn_-*iht-pbp1a* Δ*murZ* strain was similar to that of iHT-aPBP1a in a *murZ*^+^ merodiploid strain (Fig. S20*B*; Movie S20). Growth rate, protein amount, localization and relative frequency of circumferential molecules of iHT-aPBP1a were also similar in the Δ*murZ* mutant and *murZ*^+^ strain (Fig. S19 and S20*C*). Therefore, unlike iHT-bPBP2b and iHT-RodA (Fig. 4*C* and 4*D*), the circumferential velocity of iHT-aPBP1a was not decreased when PG precursor amounts were reduced. In addition, we tested whether the relative frequency of circumferential iHT-aPBP1a was changed in a *mpgA*(Y488D) Δ*pbp2b* mutant, which lacks the core PG elongasome (80, 82). Reduced catalytic activity of MpgA(Y488D) bypasses the requirement for the PG elongasome, but *mpgA*(Y488D) Δ*pbp2b* mutations are synthetically lethal with Δ*pbp1a* (80), possibly indicating that aPBP1a can substitute for the PG elongasome. Consistent with this notion, iHT-aPBP1a dynamics were the same in a *mpgA*(Y488D) Δ*pbp2b* merodiploid strain as in a WT strain (Fig. S20*C*; Movie S21).

### MpgA displays subdiffusive movement restricted to midcell

We investigated the dynamics of iHT-MpgA (previously MltG(*Spn*)) muramidase, which has been genetically linked to elongasome PG synthesis (80, 82). At an image acquisition rate of 1 FPS, the majority of iHT-MpgA molecules (≈87%) appeared to move diffusively, but seemed to be constrained mainly to the midcell region of cells (Fig. 1*B*; Movie S22), consistent with a previous report that sfGFP-MpgA localizes at midcell (34). The remaining ≈13% of iHT-MpgA molecules were static at midcell (Fig. 1*B*). There was also a small fraction (<1%) of iHT-MpgA molecules that appeared to display rapid (>50 nm/s) movement of very short duration at midcell (Fig. 3*A*, 3*B*, and S21*A*). However, this motion was difficult to track and at the limit of the criterion used for continuous motion (six consecutive frames).

Consequently, we increased the data acquisition rate to 10 FPS. At this higher rate, iHT-MpgA did not exhibit the processive, unidirectional movement observed for the elongasome components and for aPBP1a (Fig. 1*A* and S21*B*). Instead, iHT-MpgA moved erratically, mainly in the midcell region (see below; Fig. S21*B*; Movie S23). Measured at 10 FPS, the mean velocity of iHT-MpgA moving in short, directional stretches was relatively fast (≈327 nm/s) compared to circumferential movement of other proteins (<20 nm/s), but slower than diffusing molecules in non-midcell regions (>1,000 nm/s) (Fig. S21*C*). These distinctive subdiffusive dynamics did not support persistent association of MpgA with the PG elongasome or aPBP1a during PG synthesis.

Given these unusual dynamics, we confirmed the functionality of the iHT-MpgA fusion construct. The growth and cell morphology of strains expressing iHT-MpgA were similar to those of WT, and iHT-MpgA showed expected midcell localization (Fig. S2). Additionally, full-length iHT-MpgA fusion protein was expressed at 171 ± 11% the amount of WT MpgA, with only minimal degradation products observed (Fig. S6). A mutant expressing partially active MpgA(Y488D) suppressed the essentiality of the pneumococcal PG elongasome in transformation assays (80). However, the *iht*-*mpgA* construct failed to suppress Δ*pbp2b*, Δ*rodA*, Δ*mreC*, or Δ*rodZ* in transformation assays (Table S5), indicating iHT-MpgA possessed muramidase activity. Unlike Δ*mpgA* mutants, which accumulate suppressor mutations that inactivate aPBP1a (80); no *pbp1a* mutations were detected in the *iht-mpgA* fusion strain by Sanger DNA sequencing. These results indicated that the iHT-MpgA fusion protein was substantially active and that its dynamics accurately reflected those of WT MpgA.

### Core PG elongasome components and aPBP1a display different diffusion patterns from MpgA

We next investigated diffusion dynamics by performing sm-TIRFm at 20 FPS (*Materials and Methods*). Individual molecular trajectories were tracked (Fig. S22) and used to calculate mean-squared displacements (MSD) and diffusion coefficients (Fig. 6) (106, 107). Diffusion coefficients were between 0.040 and 0.055 µm^2^/s, comparable to values determined previously for PG synthesis proteins in *S. pneumoniae*, *E. coli* and *B. subtilis* (56, 100, 108, 109). Consistent with sm-TIRFm imaging at 1 FPS (Fig. 1*B*), single molecules displayed rapid diffusive motion in non-midcell regions and much slower, processive motion at midcell, where static molecules were also observed (Fig. S22; Movies S24 to S30). MSD curves generated for diffusively moving molecules were not linear and plateaued as time increased, with alpha values <1, which is indicative of some form of subdiffusion (Fig. 6). Subdiffusion is commonly due to diffusion within a confined space, but can also be caused by membrane crowding or interactions with other molecules (110). For bacterial membrane proteins, this confined space is often the surface area of the cell membrane (110).

**Fig. 6.**
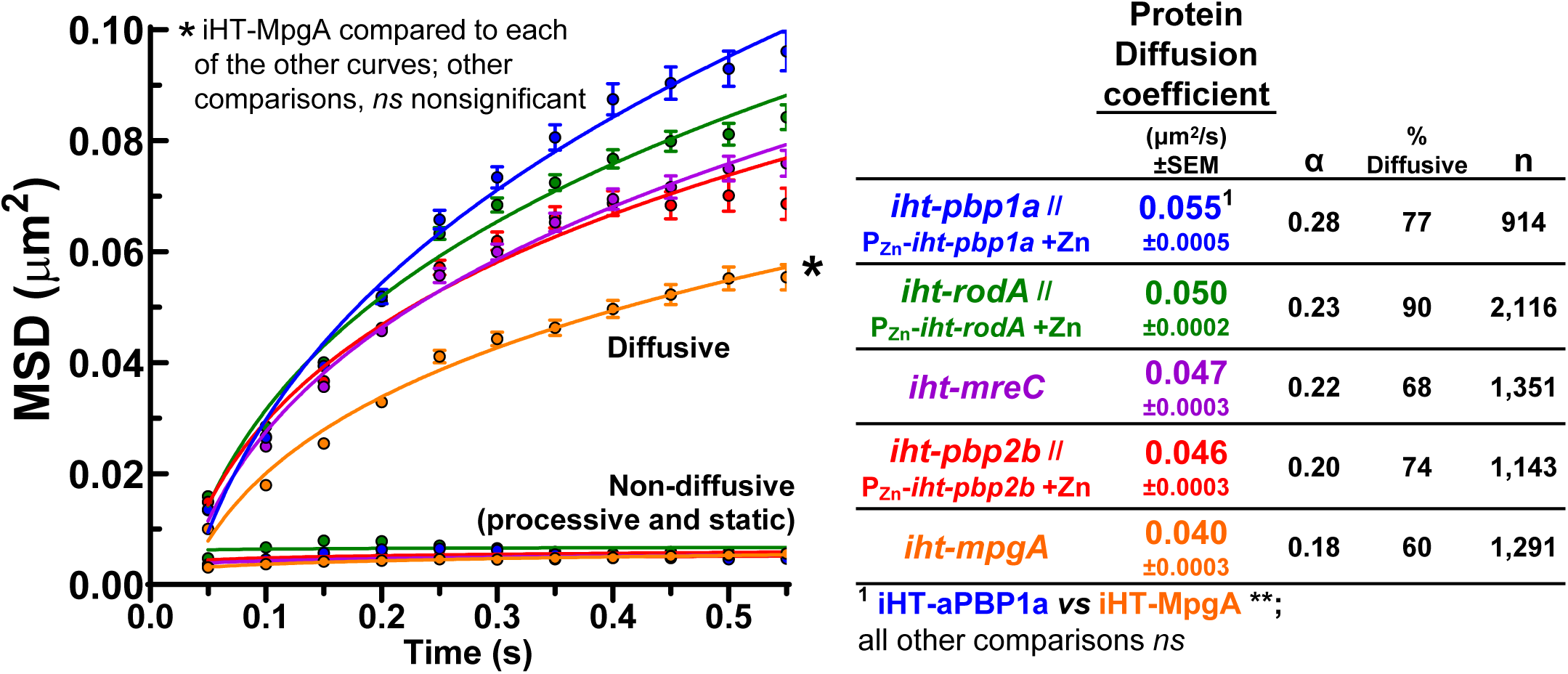
Elongation PG synthesis-associated proteins exhibited different patterns of confined diffusion. Sm-TIRFm was performed at 20 FPS on strains *iht-pbp2b* // P_Zn_-*iht-pbp2b* (IU16553), *iht-rodA* // P_Zn_-*iht-rodA* (IU16496), *iht-mreC* (IU16344), *iht-pbp1a* // P_Zn_-*iht-pbp1a* (IU16497) and *iht-mpgA* (IU15997) as described in *Materials and Methods*. 0.25 mM Zn inducer was added to strains IU16553, IU16496 and IU16497. Mean square displacements (MSD) were calculated for diffusive and non-diffusive trajectories. Circles show MSD values, and error bars represent SEM. Lines represent MSD curves fit to the data as described in *Materials and Methods*. The MSD curves were compared using a two-sample Kolmogorov-Smirnov test. \**P* < 0.05; *ns*, nonsignificant. To the right of the graph are the mean diffusion coefficients (± SEM) of diffusive trajectories, alpha values of diffusive trajectories (α), the percent of total trajectories that were classified as diffusive (% diffusive), and the total number of trajectories (n) analyzed from 2 biological replicates. α values < 1 indicate that molecules exhibited diffusion (or subdiffusion) as discussed in the text. The diffusion coefficients were compared between strains using a Brown-Forsythe and Welch’s ANOVA with a Games-Howell multiple comparison test. \*\**P* < 0.01; *ns*, nonsignificant. See Figure S22 for representative fields of cells containing multiple single-molecule trajectories.

Single-molecule trajectories of diffusively moving PG elongasome core components iHT-bPBP2b, iHT-RodA, and iHT-MreC covered most of the cell area, while non-diffusive processive and static molecules located to midcell, where active PG synthesis occurred (Fig. S22; Movies S24 to S26), consistent with results acquired at 1 FPS (Fig. S8). This suggests that the subdiffusive behavior of the core elongasome proteins was due to confinement within the cell membrane. The diffusion coefficients of iHT-bPBP2b, iHT-RodA, and iHT-MreC were similar (≈0.045 µm^2^/s) and not statistically different (Fig. 6). The MSD curves of diffusively moving iHT-bPBP2b, iHT-RodA, and iHT-MreC molecules plateaued in the same range (≈0.075 µm^2^), indicating that these core elongasome components were confined to a similarly sized area (Fig. 6) (111).

Diffusively moving iHT-aPBP1a molecules appeared to have a slightly greater diffusion coefficient (≈0.055 µm^2^/s) compared to those of the core components of the PG elongasome (Fig. 6); however, this apparent difference was not statistically significant. iHT-aPBP1a molecules also appeared to have a higher plateau in MSD plots (≈0.1 µm^2^) (Fig. 6), consistent with diffusion over the whole cell (Fig. S22 and Movie S27); however, again, this modest difference was not statistically different. Again, non-diffusive, processively moving and static iHT-aPBP1a molecules were mostly confined to midcell (Fig. S22), as was observed by sm-TIRFm at 1 FPS (Fig. 5 and S20*A*). Interestingly, we also observed some diffusive iHT-aPBP1a molecules that stopped moving, paused for a short time (<1 s), and then resumed diffusive movement (Movie S28). This pausing behavior was not readily detectable for diffusing components of the PG elongasome. We conclude that diffusing aPBP1a molecules move similarly to components of the core PG elongasome in pneumococcal cells, but there is an intermediate paused state during diffusion of aPBP1a.

Finally, diffusion analysis confirmed the unusual pattern of confined movement of iHT-MpgA molecules at midcell. A substantial fraction of diffusing iHT-MpgA molecules appeared to be confined to midcell, rather than over whole cells, like iHT-bPBP2b, iHT-RodA, iHT-MreC, and iHT-aPBP1a (Fig. S22; Movie S29). This conclusion was corroborated by manually counting the location of iHT-MpgA molecules from data (1 FPS) in Figure 1*B*. This analysis showed that 58% (65/113) of subdiffusive iHT-MpgA molecules were located within 125 nm of midcell septa or equatorial rings (Fig. S8, right). By contrast, no diffusing iHT-MreC molecules were detected at midcell (Fig. S8). Confined subdiffusion of iHT-MpgA is consistent with the low plateau (≈0.05 µm^2^) in the MSD plot, which was statistically different from those of iHT-bPBP2b, iHT-RodA, iHT-MreC, and iHT-aPBP1a (Fig. 6). In addition, the diffusion coefficient (≈0.04 µm^2^/s) of iHT-MpgA was statistically lower than diffusing iHT-aPBP1a over the body of cells (Fig. 6), consistent with the subdiffusive velocity of IHT-MpgA detected by sm-TIRFm at 10 FPS (Fig. S21C).

Similar to iHT-aPBP1a, iHT-MpgA subdiffusive molecules were observed transitioning between diffusive motion and paused states (Movie S30). Together, these results indicate that the confined, subdiffusive movement of MpgA molecules is fundamentally different from that of components of the pneumococcal core PG elongasome and aPBP1a.

## DISCUSSION

This paper shows that the core PG elongasome moves circumferentially around the midcell of dividing *S. pneumoniae* cells (Fig. 1). The velocity of core elongasome members bPBP2b, RodA, and MreC was statistically the same (≈11 nm/sec) (Fig. 3), which is slightly slower than the velocity of the bPBP2x and FtsW (≈20 nm/sec) components of the septal PG synthase (34). The processive motion of elongasome members was driven by PG synthesis and was independent of FtsZ treadmilling (Fig. 4 and 5), and this motion was not dependent on stage of cell division (Fig. S10*B*). Strikingly, the majority of bPBP2b or RodA molecules moved circumferentially in cells that severely underproduced bPBP2b (≈15% of WT) or likely RodA (Fig. 1*B*). In contrast, in cells expressing nearly WT protein levels, a minority of bPBP2b, RodA, and MreC molecules either moved circumferentially and synthesized PG at midcell or exhibited nonmoving static behavior at midcell (Fig. 1*B* and S8). The majority of core elongasome components moved diffusively in the membrane over cell bodies, where PG synthesis was not detected by FDAA labeling of exponentially growing cells (Fig. 2 and S9).

These observations indicate that pneumococcal elongation PG synthesis results from processive, circumferential movement of PG elongasomes that are confined to a narrow zone at midcell (Fig. 7 and S23). This pattern contrasts with the elongasome movement guided by short MreB filaments and driven by PG synthesis that occurs over the body of many rod-shaped bacteria (16). MreB-guided PG synthesis results in a dense mesh of concentrically oriented PG glycan strands along the inner, sidewall surface of *B. subtilis* cells (112). The circumferential movement of the pneumococcal PG elongasome anticipates that peripheral PG glycan strands may likewise be concentrically aligned in *S. pneumoniae* cells. These observations also indicate that in exponentially growing WT cells, the core elongasome components are present in excess, with only a minority of bPBP2b, RodA, and MreC engaged in active PG synthesis at midcell rings (Fig. 2 and S9). In a new study of competence induction, processive, bidirectional movement of GFP-bPBP2b and RodA-mNeonGreen at midcell was detected by ensemble-TIRFm in *S. pneumoniae* laboratory strain R800 (113), which shows significant differences in shape and the timing of cell division compared to the progenitor D39 strains used here (19, 33). The velocities of bPBP2b or RodA were not changed by competence induction (113).

**Fig. 7.**
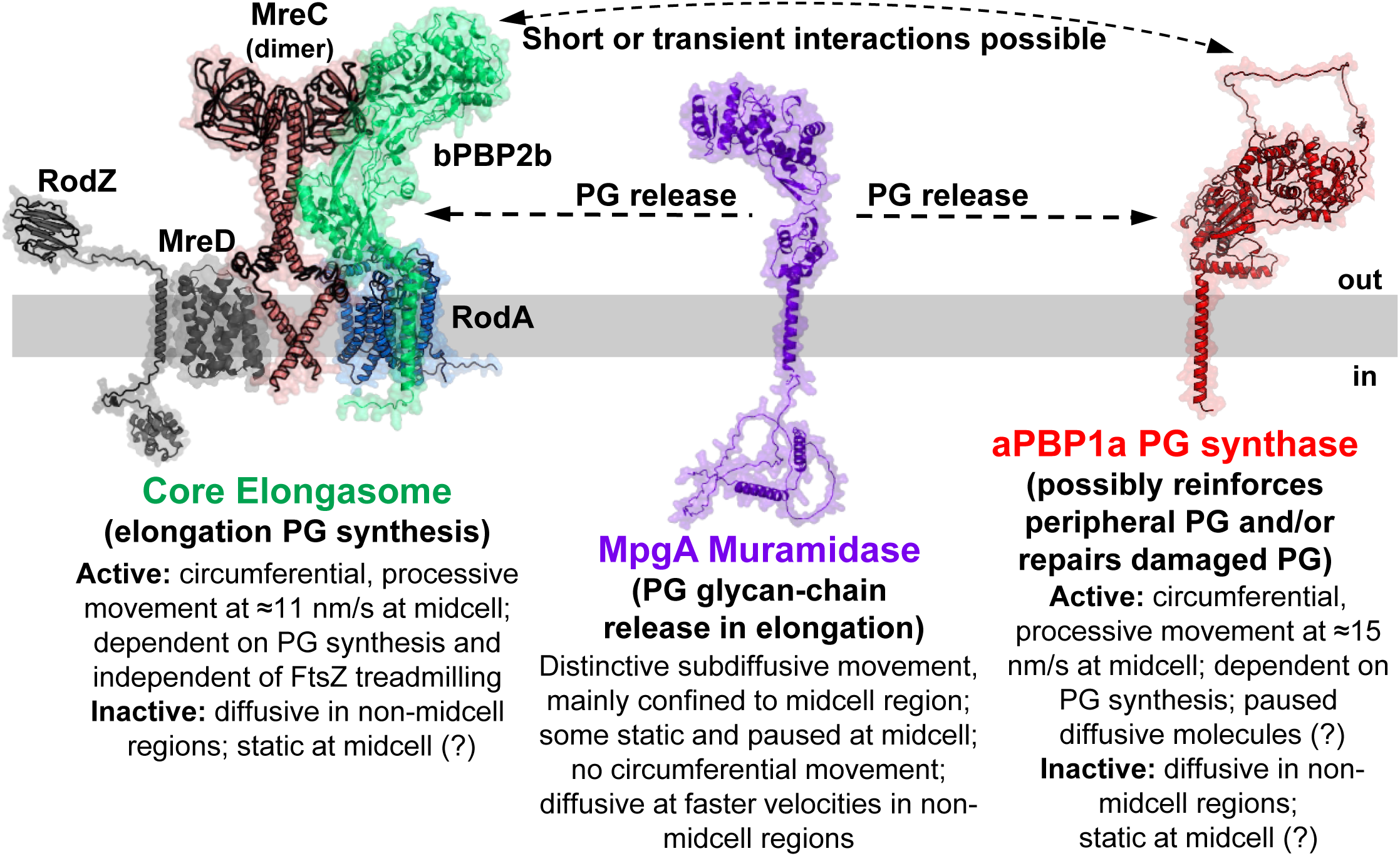
Summary model of the composition and dynamics of core elongasome components (bPBP2b, RodA, and MreC) and proteins linked to PG elongation synthesis (aPBP1a and MpgA) in growing *S. pneumoniae* cells. The circumferential, processive movement of bPBP2b, RodA, and MreC reported here likely applies to the other core elongasome components RodZ and MreD, which were not determined in this study. In early, predivisional cells, the core PG elongasome is located in the FtsZ-ring at the equator midcell of a daughter cell. In later divisional cells, the core PG elongasome is in the outer ring of the midcell septal annular disk. The motion of MreC was shown to be the same in early or late divisional cells. Class A aPBP1a also displays circumferential, processive motion at midcell that depends on PG synthesis, but its dynamics indicate that aPBP1a is not a persistent member of the core PG elongasome, although short interactions are possible. In contrast, MpgA moves in a distinctive type of subdiffusion in the midcell region. Components of the core elongasome, aPBP1a, and MpgA also move diffusively outside of midcell rings. There are also nonmoving, static molecules (> 6 s) of elongasome proteins and aPBP1a, mainly in midcell regions. It is not clear whether static molecules are synthesizing PG or are in a non-synthesizing transition state (see *Discussion*). In addition, diffusing molecules of aPBP1a and MpgA were infrequently detected pausing (< 1s) before resuming motion. See text for additional information.

The results presented here further raise the issue of what directs circumferential movement of elongasome components in tracks at midcell. The velocity of elongasome components was not changed when treadmilling was greatly impeded in an *ftsZ*(GTPase) mutant (Fig. 4*A*). Independence of FtsZ treadmilling fits the observation that FtsZ was not detected in the outer midcell ring where peripheral PG synthesis occurs later in division (Fig. S1*B*) (32). The similar velocity of MreC in predivisional cells when elongasomes are still located in the FtsZ-ring as in later-divisional cells when the elongasome is in outer midcell ring (Fig. S10*B*) is also consistent with independence of FtsZ treadmilling. In addition, when RodA or bPBP2b was depleted, no unassociated bPBP2b or RodA was detected being driven at the rate of treadmilling FtsZ (≈34 nm/sec; Fig. S17*D* and S17*E*), as occurs for components of the septal PG synthase of *E. coli* (100). Moreover, a severe decrease in FtsZ treadmilling speed led to only a marginal (≈17%) decrease in the speed of the septal PG synthase in *S. pneumoniae* (34) and no decrease in *S. aureus* (55). In *B. subtilis*, a slightly greater decrease (≈40%) in the speed of the septal PG synthase when FtsZ treadmilling was moderately or greatly reduced has recently been attributed to indirect effects (56). Thus, all evidence supports the conclusion that the movement and speed of the septal and elongasome PG synthases in WT *S. pneumoniae* cells are driven by PG synthesis, independent of FtsZ treadmilling.

It remains unknown what determines the tracks for the movement of the pneumococcal septal or elongasome PG synthases in the inner and outer midcell rings, respectively (Fig. S1). The muropeptide-crosslinked PG glycan strands to which new PG is added may provide a rigid structure that directs processive PG synthesis in a linear direction. Moreover, there appears to be a limited number of available sites in midcell rings for septal and elongasome PG synthases to load and synthesize PG. As noted above, only a minority of elongasome components are engaged in PG synthesis in WT cells (Fig. 1*B*, 2, and S23). However, when the cellular amount of bPBP2b was reduced to ≈15% of the WT level, the majority of bPBP2b molecules moved processively to synthesize PG (Fig. 1*B*), consistent with a limited number of spots on rings for active PG synthase complexes. Interestingly, ≈26% of circumferentially moving bPBP2b or MreC molecules reversed direction within the timeframe of these experiments. Direction reversal was also reported for processively moving members of the septal PG synthases of *S. aureus* (55) and *B. subtilis* (114). This directional transitioning may be indicative of PG synthase complexes stopping and reinitiating synthesis of a new glycan strand in the opposite direction (55, 114).

We also observed nonmoving, static molecules of bPBP2b, RodA, and MreC expressed at WT concentrations, primarily at midcell and to a lesser (≈5-fold) extent elsewhere in cells (Fig. 1*B* and S8). When elongasome PG synthesis was halted by expression of catalytically inactive proteins, circumferential movement also stopped and molecules became static (Fig. 5, S13*C*, and S13*D*) (69). To our knowledge, there is no direct evidence that PBPs static for several seconds are actively synthesizing PG, as suggested for aPBPs in *E. coli* (71, 74). To the contrary, when PG synthases stop synthesis, they stop moving (Fig. 5, S13*C*, and S13*D*) (34, 55, 56, 115). Therefore, we think it more likely that static PG synthases at midcell in WT cells may be incompletely assembled complexes or complexes waiting to assume an available site for PG synthesis, possibly stabilized by noncovalent binding of PBP TP domains to acceptor peptides in PG (56, 109). Consistent with this idea, we observed transitions of MreC molecules from the processive to static states (<5%), and *vice versa* (<1%). The mechanisms that arrange and limit the number of active pneumococcal PG elongasomes remain to be determined.

The processive movement of pneumococcal Class A aPBP1a strikingly contrasts with the diffusive motion for Class A PBPs reported previously in *E. coli* and *B. subtilis* (71, 74). Earlier IFM studies colocalized aPBP1a with elongasome components bPBP2b and MreC (35). Instead of diffusive motion, aPBP1a molecules at midcell moved circumferentially or were static (Fig. 5 and S20), and processive movement was abolished in a *pbp1a*(TP) mutant (Fig. 5). Processively moving aPBP1a also reversed direction infrequently (<9%). However, several results indicate that aPBP1a is not a persistent member of the core PG elongasome. The duration of aPBP1a processive motion was considerably shorter (≈10 s) than that of the PG elongasome components (≈23 s) (Fig. 3*B*). In addition, the relative frequency of circumferential motion of aPBP1a was unchanged in a suppressed Δ*pbp2b mpgA*(Y488D) mutant that lacks an intact PG elongasome (Fig. S20*C*). Like iHT-bPBP2b, iHT-aPBP1a was underproduced (10% of WT) when expressed from its native chromosomal locus (Fig. 1*B* and S5*B*). But, unlike iHT-bPBP2b, circumferentially moving aPBP1a was not the majority species at the low expression level (Fig. 1*B*). In addition, unlike bPBP2b and RodA, aPBP1a processive motion was not decreased in a Δ*murZ* mutant (Fig. S20*B* and S20*C*), which may reflect a different kinetic dependence for Lipid II substrate. Outside of midcell, aPBP1a moved diffusively in the membrane over the whole length of cells (Fig. 5 and S22). Together, these results indicate that aPBP1a is not a persistent member of the pneumococcal core PG elongasome (85), although shorter or transient interactions are possible (Fig. 7).

aPBPs have been proposed to play roles in normal PG synthesis and in repair of damaged PG (1, 39, 73, 116). In rod-shaped bacteria, it was postulated that when diffusing aPBPs encounter damaged PG, they cease moving and become static for relatively long times (≈5 s) (71, 74). For reasons discussed above, long static PG synthases may not be synthesizing PG. We also detected another aPBP1a motion away from midcell in exponentially growing cells. Diffusing aPBP1a molecules infrequently paused for short (<1 s) intervals, before resuming diffusive motion (Movie S28). It remains to be determined whether static or paused aPBP1a molecules are repairing damaged PG in *S*. *pneumoniae* cells.

In Gram-positive bacteria, the PG in the septal annular disk has a core of concentric ordered PG glycan strands covered by a dense mesh of randomly oriented PG strands that face the cell membrane (112). It has been postulated that aPBPs may synthesize this layer of randomly oriented, inner-facing PG strands (39), although remodeling by PG hydrolases followed by resynthesis could potentially contribute to this disordered pattern. Circumferential, processive movement suggests that aPBP1a synthesis may result in concentric, ordered PG glycan strands, rather than randomly oriented strands. Glycan strands synthesized by the shorter tracks of aPBP1a motion are presumably shorter than those synthesized by the more processive core PG elongasome. These shorter strands may be remodeled and crosslinked into the longer strands to provide additional strength to the peripheral PG layer. The dynamics of the other pneumococcal Class A PBPs, aPBP2a and aPBP1b, remains to be determined. The circumferential movement of an aPBP potentially has another implication to bacteria that elongate only from their poles, such as *Rhizobiales* species, including *Agrobacterium tumefaciens* (117, 118). Polar elongasome complexes in these bacteria lack bPBP:SEDS PG synthases, and PG elongation is carried out only by a single essential aPBP (117, 118), which may move circumferentially during PG elongation.

Finally, the subdiffusional, confined motion of the MpgA muramidase at midcell was unexpected. IFM showed that MpgA (formerly MltG(*Spn*)), like aPBP1a, colocalized with MreC to the outer ring in later-divisional cells (80). MpgA is an essential muramidase that cleaves newly synthesized PG glycan chains 7 disaccharides from the undecaprenol membrane anchor (82). A LysM domain in MpgA correctly places the cleavage point in the glycan chain. Therefore, MpgA is thought to act as a PG release factor that frees newly synthesized PG for crosslinking into existing PG (82). Instead of moving circumferentially, most MpgA molecules moved at subdiffusion speeds (≈300 nm/sec) that were slower than that of freely diffusing MpgA (>1,000 nm/sec). However, instead of moving over the whole body of cells, most MpgA molecules were confined to regions of PG synthesis at midcell (Fig. 1*B*, S*8*, and S22). Therefore, MpgA functions separately from the core PG elongasome and from aPBP1a (Fig. 7). Only a fraction (≈15%) of MpgA molecules were static, mostly at midcell (Fig. 1*B* and S*8*), and we detected subdiffusing MpgA molecules that paused (<1 s) and then resumed movement (Movie S30), similar to aPBP1a. Whether static or paused MpgA are carrying out PG strand cleavage requires further study. Likewise, proteins that interact with and regulate MpgA activity (119) and the mechanism that confines MpgA movement to midcell await discovery.

## MATERIALS AND METHODS

Bacterial strains used were unencapsulated (Δ*cps*) derivatives of *Streptococcus pneumoniae* (*Spn*) serotype 2 strain D39W and are listed in Table S1. IU1945 and IU1824 were used as parent strains (Table S1). Detailed methods are described in *Supplemental Materials and Methods*, including: bacterial strain construction and growth conditions; ectopic expression and depletion conditions; cell labeling with HaloTag ligand; 2D-epifluorescence microscopy and analysis; sm-TIRFm sample preparation; sm-TIRFm imaging; sm-TIRFm image analysis; single-molecule tracking and diffusion analysis; 3D-structured illumination microscopy and analysis; quantitative western blotting; transformation assay.

## ACKNOWLEDGEMENTS

We thank laboratory members and Jie Xiao (Johns Hopkins) for discussions about this work; John D. Richardson and Ziyun A. Ye for technical assistance with some experiments, Jim Powers (Indiana University Bloomington) for advice about light microscopy; Mike VanNieuwenhze (Indiana University Bloomington) for FDAA reagents; Luke Lavis (Janelia Lab) for Fluor JF549; Reinhold Brückner, and Regine Hakenbeck (Kaiserlautern University) for anti-bPBP2x antibody; and Suzanne Walker and David Z. Rudner (Harvard Medical School) for antibodies against pneumococcal PG synthesis proteins. This work was supported by NIH brant R35GM131767 (to MEW), NSF grant MCB1027504 (to SLS); NIH grant RO1AI148752 (to Suzanne Walker); NIH grant T32 GM109825 (to AJP); NIH grant F31AI138430 (to MML); NIH grants T32 GM007753 and F30 AI156972 (to JEP), and NIH equipment grant S10OD024988 to the Indiana University Bloomington (IUB) Light Microscopy Imaging Center.

## AUTHOR CONTRIBUTIONS

AJP, MML, KEB, SLS, and MEW contributed to the conception or design of this study. AJP, MML, KEB, MAT, JEP SLS, HCTT, and MEW contributed to the acquisition, analysis, and interpretation of the data. KEB, HCTT, and MEW contributed to the writing of the manuscript with input from the other authors.

## DATA AVAILABILITY

The data that support the findings of this study are presented in the paper, including the Supplemental Information and Dataset S1.

