## Supplemental Tables and Figures for "Elongasome core proteins and class A PBP1a display zonal, processive movement at the midcell of *Streptococcus pneumoniae*"

Running title: Dynamics of pneumococcal peripheral PG synthesis

Key words: processive movement of PG synthesis proteins; PBP2b:RodA:MreC elongasome dynamics; Class A PBP circumferential movement; MpgA muramidase confined subdiffusion; diffusion of non-active PG synthesis proteins

[2]Corresponding author:

Malcolm E. Winkler
Department of Biology
Indiana University Bloomington
1001 East Third Street
Bloomington, Indiana USA 47405


**This file contains:**

Supplemental Materials and Methods

Supplemental Tables S1 to S5

Supplemental Legends for Movies S1 to S30

Supplemental References

Supplemental Figures S1 to S23 with Legends

### SUPPLEMENTAL MATERIALS AND METHODS

**Bacterial strain construction and growth conditions.** Bacterial strains used were unencapsulated (Δ*cps*) derivatives of *Streptococcus pneumoniae* (*Spn*) serotype 2 strain D39W and are listed in Table S1 (1). IU1945 and IU1824 were used as parent strains (Table S1). Linear DNA amplicons synthesized by overlapping fusion PCR were transformed into competent pneumococcal cells and grown on trypticase soy agar II plates (BD BBL, 221261) containing 5% (v/v) defibrinated sheep blood (TSAII-BA) and incubated at 37°C in an atmosphere of 5% $CO_2$ as described previously (2). Primers used to synthesize amplicons are listed in Table S2. For antibiotic selection, TSAII-BA plates contained 250 µg/mL kanamycin, 150 µg/mL spectinomycin, 0.3 µg/mL erythromycin, 250 µg/mL streptomycin, 2.5 µg/mL chloramphenicol, or 0.25 µg/mL tetracycline, as needed. Ectopic expression of various genes was achieved with a $P_{Zn}$ zinc-inducible promoter in the ectopic *bgaA* site (3). Zn inducer contained 0.4 or 0.5 mM $ZnCl_2$ and one-tenth the concentration of $MnSO_4$ (Zn + [1/10] Mn) and was added to TSAII-BA plates or BHI broth for gene induction. Mn was included with Zn to prevent zinc toxicity (4-6). During construction of strains IU12272, IU12345, IU15419, IU16134, IU16136, IU16182, IU16184, IU16202, IU16204, IU16232, IU16236, IU16239, IU16252 and IU16281, 0.4 mM Zn inducer was added to cells throughout transformation, plating, single colony isolation, and storage in brain heart infusion broth (BHI; BD Bacto 237500). During construction of IU11258, 0.5 mM Zn inducer was added to cells throughout transformation, plating, single colony isolation, and storage in BHI broth.

For overnight cultures, frozen glycerol stocks were inoculated into 4 mL BHI broth, serially diluted, and incubated statically for 12-16 h at 37°C in an atmosphere of 5% $CO_2$.

0.4 mM Zn inducer was added to overnight cultures and incubation times were ≤12 h when performing experiments with strains IU11258 (Δ*pbp2b* // $P_{Zn}$-*pbp2b*), IU12345 (Δ*mreC* // $P_{Zn}$-*mreC*), IU16136 (Δ*rodA* // $P_{Zn}$-*rodA*), IU16202 (*iht-rodA* Δ*pbp2b* // $P_{Zn}$-*pbp2b*), IU16204 (*iht-pbp2b* Δ*rodA* // $P_{Zn}$-*rodA*), IU16232 (*iht-pbp2b*(S391A) // $P_{Zn}$-*pbp2b*), IU16239 (*iht-rodA*(D283A) // $P_{Zn}$-*rodA*), and IU16281 (*iht-pbp2b* Δ*mreC* // $P_{Zn}$-*mreC*). Growth was monitored by $OD_{620}$ as described previously (7). Unless noted, experiments were performed in C+Y media adjusted to pH 7.1 by adding 500 µL of 1 M HCl to 43.9 mL of C+Y (8,9). 2-4 mL overnight cultures in BHI and still in exponential phase ($OD_{620}$ = 0.1–0.4) were collected by centrifugation (21,100 X *g* for 5 min at room temperature (RT)), washed in 1 mL C+Y with centrifugation, resuspended in 4 mL fresh C+Y, and diluted to $OD_{620}$ = 0.003-0.005 in 5 mL fresh C+Y, with or without Zn inducer. Cultures were incubated statically at 37°C with 5% $CO_2$, and growth was monitored every 45-60 m by measuring $OD_{620}$.

**Ectopic expression and depletion conditions.** Ectopic expression of iHT-bPBP2b (IU16553), iHT-RodA (IU16496) or iHT-aPBP1a (IU16497). Overnight cultures were grown in BHI broth without Zn, centrifuged, washed with centrifugation in C+Y without Zn, and diluted into C+Y with indicated concentrations of Zn inducer. For single-molecule total internal reflection fluorescence microscopy (sm-TIRFm) and two-dimensional epifluorescence microscopy (2D-FM), HT labeling and washing were done in C+Y containing the specified concentration of Zn inducer. Agarose pads also contained the indicated concentration of Zn inducer. Western blotting was performed as described below.

Ectopic expression of FtsZ(D214A) (IU16091, *iht*-$L_6$-*pbp2b ftsZ*$^+$ // Δ*bgaA*::*tet*-$P_{Zn}$-*ftsZ*(D214A)). Overnight cultures of IU16091 and control strain IU15928 were grown in BHI broth without Zn, centrifuged, washed with centrifugation in C+Y, and diluted into C+Y with 0.25 mM Zn inducer. HT labeling and washing were done in C+Y with 0.25 mM Zn inducer, and agarose pads also contained 0.25 mM Zn inducer.

Expression of iHT-bPBP2b(S391A) (IU16232, *iht*-$L_6$-*pbp2b*(S391A) // Δ*bgaA*::*tet*-$P_{Zn}$-*pbp2b*$^+$) and iHT-RodA(D283A) (IU16239, *iht*-$L_6$-*rodA*(D283A) // Δ*bgaA*::*tet*-$P_{Zn}$-*rodA*$^+$). Overnight cultures of IU16232 and IU16239 were grown in BHI broth with 0.4 mM Zn inducer for 12-13 h. For sm-TIRFm, cultures were washed in C+Y without Zn and diluted to $OD_{620}$ ≈0.036 in C+Y lacking Zn (defined as the start of depletion). Cultures were grown for 1 h and labeled with HT-ligand and prepared for sm-TIRFm as described elsewhere in *Materials and Methods*. Imaging occurred 3 h into depletion. For sm-TIRFm of IU16232 with Zn inducer, overnight cultures were diluted to $OD_{620}$ = 0.005 in C+Y with 0.25 mM Zn inducer added. At $OD_{620}$ ≈0.1, cultures were labeled and prepared for sm-TIRFm as described above using C+Y with 0.25 mM Zn inducer. For 2D-FM and western blotting, overnight cultures were washed in C+Y without Zn and diluted to $OD_{620}$ = 0.003 in C+Y with or without 0.2 mM Zn inducer. Samples were processed for western blotting at 3.5 h, and cells were labeled and imaged for 2D-FM at 3.5 and 4 h.

Depletion of MreC (IU16281). Overnight cultures of IU16281 were grown in BHI broth with 0.4 mM Zn inducer for ≈12 h, washed with centrifugation in C+Y without Zn, diluted to $OD_{620}$ ≈0.030 in C+Y with 0.2 mM Zn inducer, and grown for 1 h. For sm-TIRFm, cultures were centrifuged and resuspended in C+Y without Zn (defined as the start of depletion) and immediately HT labeled and prepared for sm-TIRFm. Imaging occurred 3 h into depletion. For 2D-FM and western blotting, after 1 h growth in C+Y with 0.2 mM Zn inducer, cultures were washed with centrifugation in C+Y without Zn and diluted to $OD_{620}$ = 0.005 in C+Y with or without Zn. At $OD_{620}$ = 0.1-0.2, samples were either HT labeled and imaged with 2D-FM or processed for western blotting as described below.

Depletion of bPBP2b (IU11258 and IU16202) or RodA (IU16136 and IU16204). Overnight cultures of IU11258, IU16202, IU16136 and IU16204 were grown in BHI broth with 0.4 mM Zn inducer for ≈12 h. For sm-TIRFm of IU16202 and IU16204, cultures were washed with centrifugation in C+Y without Zn, and diluted to OD620 ≈0.036 in C+Y lacking Zn (defined as the start of depletion). Cultures were grown for 1.5 h then HT labeled and prepared for sm-TIRFm. Imaging occurred 3 h into depletion. For 2D-FM and western blotting of IU11258, IU16202, IU16136 and IU16204, overnight cultures were washed with centrifugation in C+Y without Zn and diluted to $OD_{620}$ = 0.003 in C+Y with or without 0.2 mM Zn. At 3 h, samples were HT labeled and imaged with 2D-FM as described. For western blotting, samples were processed at 3 or 4 h.

**Cell labeling with halotag-ligand.** Cells were grown in C+Y at 37° C with 5% $CO_2$ until $OD_{620}$ = 0.08-0.15. For saturation labeling of HT-fusion proteins with HT-ligand, 500 μL of culture was incubated with a final concentration of 500 nM HT-TMR (Promega #G8252). For single molecule labeling, JF549 ligand (gift of Luke Lavis, Janelia) was added to 500 μL culture to the final concentrations listed in Table S4. Unless noted, samples were incubated at 37° C in the dark with shaking at 500 rpm for 15 m. Labeled cells were centrifuged at 21,100 x *g* for 2.5 m at RT and washed with centrifugation 3 times in 500 μL fresh C+Y to remove unbound ligand. Cell pellets were resuspended in

500 µL of fresh C+Y for sm-TIRFm. For 2D-FM, cell pellets were resuspended in 25-100 µL of fresh C+Y.

**2D-epifluorescence microscopy (2D-FM) and analysis.** 1 µL of cells were placed on a glass slide with coverslip and imaged on a Nikon Eclipse E-400 epifluorescence phase-contrast microscope with a Nikon Intensilight C-HGFI epifluorescence illuminator. A CoolSNAP HQ2 CCD camera (Photometrics) was used to capture images, and analyzed with Nikon NIS-Elements BR imaging software. Phase-contrast images were collected with a 20-50 ms exposure time, and HT images were collected using a Texas Red-HYQ filter (Ex 532-587 nm, Em 560 nm) with a 1 s exposure time. Additional image processing was completed with ImageJ (10). Demographs were constructed using Microbe J as described previously (11). Length and width measurements of cells were made from phase-contrast images using Nikon NIS-Elements BR software.

**TIRFm sample preparation.** Samples were prepared for TIRFm as described previously, with some differences (11). Glass coverslips were soaked in an acid-ethanol mixture (25% concentrated HCl, 25% $H_2O$, 50% ethanol) for 12-24 h, washed with gentle agitation 5-10 times in Type I (Barnsted) $H_2O$, and dried with lens paper. Glass slides were cleaned with 70% ethanol and gene frames (1.0 cm x 1.0 cm) (Thermo-Fisher # AB0576) were attached. 1.5% (w/v) agarose (Sigma BioReagent A9414) in C+Y was melted and agarose pads were made within Gene Frames as described previously (12). Where noted, Zn inducer was added to C+Y prior to melting agarose. 1.2 µL of HT-labeled cells were spread on a coverslip, briefly air-dried (<3 m) and placed cell side down onto the agarose pad. Samples were equilibrated to RT for 10 m, then warmed to 37°C for at least 20 m prior to TIRFm. The TIRFm objective and stage were warmed to 37° C for at least 20 m prior to imaging.

**TIRFm imaging.** TIRFm was performed in the Indiana University Light Microscopy Imaging Center on a DeltaVision OMX-SR microscope (GE Healthcare), with an Apo N 60X/1.49 TIRF objective (Olympus) and PCO.edge 4.2 sCMOS cameras (PCO). Laser lines were 488 nm, with emission filter 500–550 for GFP or 561 nm with emission filter 609–654 for HT. Differential Interference Contrast microscopy (DICm) images were collected during imaging to determine cell locations. For an image acquisition rate of 1 frame per s (FPS), DICm exposure time was 9 ms at 10% laser power (T), GFP exposure time was 45 ms at 10% T, and HT exposure time was 45 ms at 100% T for 180 s. For 10 FPS imaging, DICm exposure time was 9 ms at 10% T and HT exposure time was 45 ms at 100% T for 30 s. For 20 FPS imaging, used for single-molecule tracking with diffusion analysis, DICm exposure time was 3 ms at 10% T and HT exposure time was 24 ms at 100% T for 100 s. Images were acquired in sequential mode, and channels were aligned using SoftWoRx (GE Healthcare).

**TIRFm image analysis.** Data were processed and analyzed using FIJI (10). DICm channels were inverted to show dark cell bodies with light cell boundaries. Some images required manual alignment of channels. To account for drift of cells during imaging, the HyperstackReg plugin was used to perform rigid-body registration on the DICm channel, which was then applied to the other channels of the image (13).

Movement pattern analysis was completed on 1 FPS images to determine the relative frequency of different types of single-molecule motion of HT-fusion proteins. For each field, each cell was examined by direct visualization for the entirety of the imaging duration

(180 s). HT-fusion molecules were identified by eye with molecules having a diameter of approximately 3-6 pixels and signal intensity greater than ≈1.2-fold above background. Only fields where >80% of cells displayed zero or one labeled molecules were analyzed. All cells in a given field were analyzed unless only partially visible (for example, cells on the edge of the field). Movements of HT-fusion proteins were classified as circumferential, diffusive, static, or transitional. Circumferential molecules were defined as particles moving in one direction for 6 or more consecutive frames and with a linear velocity ≥5 nm/s as determined by kymograph analysis (described below). No more than one circumferential particle was scored for each cell. Static molecules were defined as particles appearing motionless for 6 or more consecutive frames, or moving with a linear velocity <5 nm/s as determined by kymograph analysis. No more than one static particle was scored for each cell. Diffusive molecules were defined as particles that moved, but not in a consistent direction, for 6 or more non-consecutive frames within a period of 90 s. No more than one diffusive particle was scored for each cell. Transitional molecules were defined as 6 or more frames of one motion type directly followed by 6 or more frames of another motion type. More than one type of movement can be seen within a cell without being classified as transitional. Each particle analyzed was determined to be either within an area of the cell where PG synthesis was occurring (midcell) or not (non-midcell). The minimum duration of six frames (when imaged at a rate of 1 FPS) was based on previous work (11). The linear velocity cutoff of 5 nm/s to distinguish between circumferentially moving and static (i.e., slow or non-moving) molecules was based on a molecule moving at 5 nm/s for the minimum duration of 6 frames (6 s). This molecule would move a total distance of 30 nm, less than half of a pixel width of the system (effective pixel size 78.6

nm). We were not confident in distinguishing movements below this cutoff from the intrinsic movement of the imaging system.

Circumferential velocities of single molecules were determined using kymograph analysis as described previously (11) and further detailed here. Proper measurement of circumferential velocities required removal of the spatial scale from images before analysis. Kymographs were generated using the FIJI line tool and reslice function. Circumferential velocities were calculated using V = P / [T * *tan*(A)], where V is the circumferential velocity (in nm/s), P is the pixel size (78.6 nm), T is the time between frames (in s), *tan* is the tangent function, and A is the angle (in radians) of the molecule path measured from the kymograph. Circumferential durations were measured from kymographs by drawing a vertical line starting from the beginning of the circumferential molecule path to the end. The duration (in s) equals the length of the line (in frames, represented as pixels in the kymograph) divided by the image acquisition rate (in FPS). Circumferential distances of single molecules were calculated by multiplying the velocity by the duration for a given molecule. Primary velocity, duration, and distance data are compiled in Supplemental Dataset S1.

**Single-molecule tracking with diffusion analysis.** Single-molecule tracking with diffusion analysis was performed on images acquired at 20 FPS. Molecules were identified and localized within each frame, and localizations were linked across frames to construct single molecule trajectories (or tracks) using the ImageJ plugin TrackMate (14). The TrackMate Laplacian of Gaussian detector (LoG) was used for particle localization with the following settings: spot radius was 500 nm; threshold was 3 (dimensionless); and sub-localization box was on. In certain cases, spurious spots were generated at the edges of a frame and were removed using an appropriate spot filter in the X or Y position. Spot linking across frames to generate particle trajectories was performed using the simple Linear Assignment Problem (LAP) tracker with the following settings: linking max distance was 300 nm; gap-closing max distance was 0; and gap-closing max frame gap was 0. Trajectories shorter than 5 frames were manually discarded, along with any erroneous trajectories that were external to a cell outline. Multiple trajectories were typically located within a single cell. Trajectories of 5 or more frames were partitioned into diffusive and non-diffusive groups according to two empirically determined criteria: trajectories with displacement > 0.13 μm AND velocity standard deviation > 0.63 μm/s comprised the diffusive group, while the remaining trajectories comprised the non-diffusive group, which included processive/circumferentially moving and static molecules. Trajectory displacement was defined as the distance between the starting point of a trajectory and the end point of a trajectory. Velocity standard deviation was defined as the standard deviation of all the instantaneous velocities for a trajectory. Instantaneous velocities were defined as the distance a particle moved in a trajectory from one frame to the next (in μm) divided by the time between frames (0.05 s for 20 FPS imaging).

These trajectories were imported into MATLAB, and the mean square displacement (MSD) at a given time-lag Δt (in frames) was computed for each trajectory according to the equation:

$$MSD(\Delta t) = \frac{1}{N}\sum_{i>j}\big(x(i) - x(j)\big)^2 + \big(y(i) - y(j)\big)^2,$$

where the sum is taken over indices $i = 0,1, \ldots, T$ and $j = 0,1, \ldots, T$ such that $\Delta t = i - j$, $x$ and $y$ are particle locations for a given frame $i$ or $j$, and $N = T - \Delta t + 1$, where $T$ is the total number of frames of a given trajectory (15). This MSD calculation was done automatically for each trajectory using MSDanalyzer (16), a MATLAB class compatible with imported TrackMate trajectories. The plotted MSD values (circles in Fig. 6) are the result of averaging MSD values of all trajectories for a given time-lag (ensemble average). These ensemble averaged MSD values for time-lags 0.05 to 0.55 s were then fitted to the anomalous MSD diffusion equation MSD = $4Dt^{\alpha} + D_0$ to obtain MSD curves (curved lines in Fig. 6), diffusion coefficients (D; $\mu m^2/s$), alpha ($\alpha$), which in the case of confined diffusion (subdiffusion) $\alpha < 1$, and $D_0$, which accounts for the localization uncertainty inherent in the imaging conditions (17). The anomalous MSD diffusion equation was used to capture the perturbative effects of cell curvature, membrane inhomogeneity, protein-protein interactions, as well as the confined space of a bacterial cell which necessarily deviate a protein's diffusion from the special case $\alpha = 1$ corresponding to Brownian motion. Diffusion coefficients were compared between strains using Brown-Forsythe and Welch's ANOVA with a Games-Howell multiple comparisons test (GraphPad Prism v10).

**3D-structured illumination microscopy (3D-SIM) and analysis.** Cells were grown, labeled and fixed as described in sections 4.6 and 4.7 of (18). Briefly, overnight cultures were diluted to $OD_{620}$ = 0.003 in fresh BHI broth (IU16553 was supplemented with 0.3 mM Zn inducer). If HT labeling was required (IU15928 and IU16553 in Fig. 2), at $OD_{620} \approx$ 0.07, 600 µL of cells were labeled with 500 nM JF549 HT-ligand for 15 m, as described above. Cells were washed twice with centrifugation with 1 mL BHI broth at 37°C, resuspended in 600 µL BHI broth, and incubated in the dark for 20 m.

At $OD_{620} \approx 0.23$ (Fig. S9, IU9965) or immediately following the HT labeling procedure (Fig. 2, IU15928 and IU16553), 600 µL culture was transferred to a 0.22 µm microcentrifuge tube filter (Corning Costar Spin-X, CLS8160), the media was removed by vacuum filtration, and 250 µL BHI broth at 37° C was added to the filter for 3 s and then filtered away. 500 µL PBS at 37°C was added and then filtered away. The filter was removed from the vacuum hose and 250 µL BHI broth with 400 µM HADA or 125 µM TADA was pipetted up and down approximately 10X to resuspend cells. Cells were incubated at 37°C in the dark for 2.5 m, then reattached to the vacuum pump and the media was filtered away. 500 µL PBS at RT was added to the filter and immediately filtered away. This wash step was repeated, then the filter was removed from the vacuum pump. 600 µL of 4% (v/v) paraformaldehyde was pipetted up and down on the filter ≈10X to resuspend cells, and the cells were transferred to a 2 mL tube. During fixation, cells were incubated in the dark for 15 m at RT, and then incubated on ice in the dark for 45 m.

After fixation, cells were centrifuged at 16,100 × *g* for 5 m at 4°C, washed twice with centrifugation with 600 µL ice-cold PBS, and resuspended in 200 µL GTE buffer (50 mM glucose, 20 mM Tris-HCl, pH 7.5, 1 mM EDTA). Cells were centrifuged 16,100 × *g* for 5 m at 4°C, the GTE buffer was removed, and the pellets were centrifuged again and residual buffer was removed. Pellets were air dried for ≈1 m and resuspended in 3 µL Vectashield Hardset Antifade (Vector Laboratories, H-1400) with vortexing. 1.2 µL of cells was placed on a 12 mm/1.5 round coverslip (EMS, 72230-01) and a glass microscope slide place on top. The slide was incubated for 15 m in the dark at RT to allow the mounting media to cure. 3D-SIM images were acquired and processed as described in section 4.12 of (18). Reconstructed images in this work consist of 15 plane *Z*-stacks (each 0.125 µm thick) representing a depth of 1.875 µm and fully encompassing horizontal cells.

**Quantitative western blotting.** Western blotting with anti-HT, anti-MreC, and anti-bPBP2b antibodies was performed as described previously with some modifications (18). Briefly, overnight cultures were diluted to $OD_{620}$ = 0.003-0.005 in C+Y with Zn inducer where indicated, and incubated at 37°C in 5% $CO_2$. When cells reached $OD_{620}$ = 0.15-0.2, 2 mL of culture was centrifuged at 4°C at 16,100 × *g* for 5 m, washed with 4°C PBS, centrifuged again, and the supernate was removed. Pellets were placed on dry ice for 15 m, thawed at RT for 5 m, and resuspended in 80 µL SEDS lysis buffer (0.1% (v/v) deoxycholate, 150 mM NaCl, 0.2% (v/v) SDS, 15 mM EDTA, pH 8.0). Samples were vortexed vigorously before incubation at 37°C with 300 rpm shaking for 15 m, with brief vortex mixing every 5 m. Protein concentration was determined using the DC protein assay kit (Bio-Rad, 5000116). Samples were diluted 1:1 with 2x Laemmli sample buffer (Bio-Rad, 1610737) containing 5% (v/v) β-mercaptoethanol, incubated at 95°C for 10 m, and vortexed briefly.

Unless noted otherwise, 2.5 µg lysate was loaded onto a 4-15% gradient SDS-PAGE gel (Bio-Rad) and run for 1 h at 150 V. Samples were transferred to a nitrocellulose membrane at 350 mA for 90 m, then blocked in 6 mL PBS with 0.1% (w/v) Tween 20 (Sigma, P1379-500ML) (PBST) containing 50 mg/mL skim milk (BD Difco, 232100) for 20 m with gentle rocking. Blots were rinsed briefly in 5 mL PBST, then incubated for 1 h with gentle rocking in PBST with either mouse anti-HT (1:600 dilution; Promega, G9211), rabbit anti-MreC (1:1,400 dilution; (19-21)), or rabbit anti-bPBP2b (1:10,000 dilution; (18)). Blots were rinsed briefly twice in 5 mL PBST, and washed for 15 m in 5 mL PBST. Blots were incubated in PBST with either ECL anti-rabbit IgG horseradish peroxidase (HRP) antibody (1:10,000 dilution; GE Healthcare, NA934V) or anti-mouse HRP antibody

(1:1,200 dilution; Bio-Rad, 1706516) for 1 h with gentle rocking. Blots were sequentially washed in 5 mL PBST for 5 m, 5 m, 15 m, 5 m, and 5 m, and then incubated with Amersham ECL western blotting detection reagent (GE Healthcare, RPN2106) for 1 m and imaged with an IVIS imaging system (Xenogen), using a 1 m exposure time as described previously (22).

Western blotting with anti-aPBP1a and anti-MpgA antibodies was performed as described above with some modifications. Cells were grown, washed, and frozen as described above. Cell pellets were resuspended in SEDS lysis buffer according to the cell density of the culture at the time of harvest. For an $OD_{620}$ = 0.160, pellets were resuspended in 80 µL SEDS lysis buffer. Cultures with higher or lower $OD_{620}$ were resuspended in proportionally more or less SEDS lysis buffer, respectively. Samples were lysed as described above. Protein concentration was not determined. Samples were diluted in Laemmli buffer and processed as described above. Samples were loaded onto the gel (e.g., Fig. S5), and electrophoresis and transfer to nitrocelluose were done as described above. TotalStainQ (Azure Biosystems, AC2227) total protein stain was used to determine the relative amount of total protein in each lane and was used for normalization. The blot was washed for 5 m with 5 mL water, then washed for 5 m with 6 mL TotalStainQ solution. Blots were rinsed 3X for 3 m each with 6 mL TotalQ Wash solution, and then imaged with an Azure Biosystems 600 imaging system with 524 nm excitation and 572 nm emission wavelengths. The blot was washed for 5 m with 5 mL water before blocking as described above. Blotting and washes were performed as described above, but used either a rabbit anti-aPBP1a antibody (1:10,000 dilution) (20) or a rabbit anti-MpgA (1:7,000 dilution) primary antibody generated using full-length *S.*

*pneumoniae* D39 MpgA (amino acid M1 to N551) expressed in *E. coli* and purified as described in (23). No band was detected in a Δ*mpgA* strain (IU7325) using this anti-MpgA antibody (Fig. S6). An anti-rabbit IR Dye® 800CW (1:14,000 dilution; Li-Cor, 926-32211) was used as the secondary antibody. After the final wash in PBST, the blot was washed twice for 5 m each in 5 mL PBS and immediately imaged with 784 nm excitation and 832 nm emission wavelengths using the Azure Biosystem 600. Quantitation of relative protein amounts was performed as described previously (24).

**Transformation assays.** Transformations were performed as previously described (5,6). Amplicons containing Δ*rodA*::$P_c$-*erm,* Δ*pbp2b*<>*aad9*, Δ*mreCD*<>*aad9*, Δ*rodZ*<>*aad9*, Δ*pbp1b*::$P_c$-*aad9*, or Δ*pbp1b*::$P_c$-*erm* deletion mutations with ≈1 kb of flanking chromosomal DNA for homologous recombination were synthesized by PCR using primers and templates listed in Table S2. All transformation experiments were performed with no added DNA as a negative control, and with Δ*pbp1b* amplicons containing the same antibiotic selection as a positive control for competence efficiency and colony size comparison. Each transformation experiment was performed independently twice.

### SUPPLEMENTAL TABLES S1 TO S5

**Table S1.** *Streptococcus pneumoniae* D39W strains used in this study

| Strain number | Genotype (construction)[a b] | Antibiotic resistance[c] | Reference or source |
|---|---|---|---|
| IU1751 | R6 Δ*mreCD*<>*aad9* | Spc$^{R}$ | (21) |
| IU1824 | D39 Δ*cps rpsL1* | Str$^{R}$ | (1) |
| IU1945 | D39 Δ*cps* | None | (1) |
| IU6726 | D39 Δ*cps rpsL1* Δ*pbp1a*::P$_{C}$-[*kan-rpsL*$^{+}$] | Kan$^{R}$ | (6) |
| IU6741 | D39 Δ*cps rpsL1* Δ*pbp1a* | Str$^{R}$ | (6) |
| IU7325 | D39 Δ*cps rpsL1* Δ*pbp1a* Δ*mltG*::P$_{C}$-[*kan-rpsL*$^{+}$] | Kan$^{R}$ Str$^{S}$ | (6) |
| IU7397 | D39 Δ*cps* Δ*pbp2b*<>*aad9* //Δ*bga*::*kan*-t1t2-P$_{fcsk}$-*pbp2b*$^{+}$ | Spc$^{R}$ Kan$^{R}$ | (2) |
| IU7614 | D39 Δ*cps rpsL1 ftsZ*$^{+}$-P$_{C}$-[*kan-rpsL*$^{+}$] | Kan$^{R}$ | (6) |
| IU8122 | D39 Δcps Δ*bgaA*::*tet*-P$_{Zn}$-RBS$^{ftsA}$-*ftsZ*$^{+}$ | Tet$^{R}$ | (25) |
| IU8872 | D39 Δ*cps* Δ*bgaA*::*tet*-P$_{Zn}$-*mpgA*$^{+}$ | Tet$^{R}$ | (6) |
| IU8980 | D39 Δ*cps rpsL1* P$_{c}$-[*kan-rpsL*$^{+}$]-*mpgA*$^{+}$ | Kan$^{R}$ | (6) |
| IU9023 | D39 Δ*cps rpsL1* P$_{c}$-[*kan-rpsL*$^{+}$]-*pbp2b*$^{+}$ | Kan$^{R}$ | (11) |
| IU9613 | D39 Δ*cps rpsL1 rodZ*$^{+}$//Δ*bgaA*::*tet*-P$_{Zn}$-*rodZ*$^{+}$ | Str$^{R}$ Tet$^{R}$ | (20) |
| IU9760 | D39 Δ*cps rpsL1 mpgA*(Y488D) | Str$^{R}$ | (6) |
| IU9783 | D39 Δ*cps rpsL1 mpgA*(Y488D) Δ*pbp2b*<>*aad9* | Spc$^{R}$ Str$^{R}$ | (6) |
| IU9931 | D39 Δ*cps* Δ*rodZ*<>*aad9*//Δ*bgaA*::*tet*-P$_{Zn}$-RBS$^{ftsA}$-*rodZ*$^{+}$ | Spc$^{R}$ Tet$^{R}$ | (6) |
| IU9965 | D39 Δ*cps rpsL1 sfgfp*-L$_{1}$-*pbp2b* | Str$^{R}$ | (11) |
| IU9985 | D39 Δ*cps rpsL1 ftsZ*-L$_{2}$-*sfgfp* | Str$^{R}$ | (11) |
| IU10103 | D39 Δ*cps rpsL1* P$_{c}$-[*kan-rpsL*$^{+}$]-*mreC*$^{+}$ | Kan$^{R}$ | (20) |
| IU10220 | D39 Δ*cps rpsL1* Δ*bgaA::tet*-P$_{Zn}$-*mreC*$^{+}$ | Tet$^{R}$ Str$^{R}$ | (20) |
| IU10651 | D39 Δ*cps* Δ*mreCD*<>*aad9 mpgA*(Y488D) | Spc$^{R}$ | (6) |
| IU10922 | D39 Δ*cps* Δ*bgaA*::*tet*-P$_{Zn}$-*rodA*$^{+}$ | Tet$^{R}$ | (6) |
| IU10943 | D39 Δ*cps rpsL1 mpgA*(Y488D) markerless Δ*rodA*::P$_{c}$-*erm* | Str$^{R}$ Erm$^{R}$ | (6) |
| IU10945 | D39 Δ*cps rpsL1* Δ*rodA*::P$_{c}$-[*kan-rpsL*$^{+}$] *mpgA*(Y488D) | Kan$^{R}$ | (6) |
| IU11157 | D39 Δ*cps rpsL1 isfgfp*-L$_{1}$-*pbp2x* | Str$^{R}$ | (11) |
| IU11246 | D39 Δ*cps rpsL1* Δ*bgaA*::*tet*-P$_{Zn}$-*pbp2b*$^{+}$ | Str$^{R}$ Tet$^{R}$ | (25) |
| IU11258 | D39 Δ*cps rpsL1* Δ*pbp2b*<>*aad9* // Δ*bgaA*::*tet*-P$_{Zn}$-*pbp2b*$^{+}$ | Str$^{R}$ Spc$^{R}$ Tet$^{R}$ | (25) |
| IU12272 | D39 Δ*cps rpsL1* Δ*mreC*::P$_{c}$-[*kan-rpsL*$^{+}$] // Δ*bgaA::tet*-P$_{Zn}$-*mreC*$^{+}$ | Kan$^{R}$ Tet$^{R}$ | (20) |
| IU12345 | D39 Δ*cps rpsL1* Δ*mreC* // Δ*bgaA::tet*-P$_{Zn}$-*mreC* | Str$^{R}$ Tet$^{R}$ | (20) |
| IU13536 | D39 Δ*cps rpsL1* Δ*murZ* | Str$^{R}$ | (11) |

| Strain number | Genotype (construction)[a b] | Antibiotic resistance[c] | Reference or source |
|---|---|---|---|
| IU13680 | D39 Δ*cps* Δ*pbp1b*::$P_c$-*aad9* | Spc$^R$ | (20) |
| IU14290 | D39 Δ*cps rpsL1 pbp1a*-$L_0$-*ht*-$P_C$-*erm* | Str$^R$ Erm$^R$ | (11) |
| IU14738 | D39 Δ*cps rpsL1 iht*-$L_6$-*mapZ* | Str$^R$ | (11) |
| IU14850 | D39 Δ*cps rpsL1* Δ*bgaA*::*tet*-$P_{Zn}$-*ftsZ*(D214A) | Tet$^R$ Str$^R$ | (11) |
| IU15419 | D39 Δ*cps rpsL1* Δ*pbp2b::*$P_c$-[*kan-rpsL*$^+$] //Δ*bgaA::tet*-$P_{Zn}$-*pbp2b*$^+$ (IU11246 x Δ*pbp2b::*$P_c$-[*kan-rpsL*$^+$] fusion amplicon) | Kan$^R$ Tet$^R$ | This study |
| IU15433 | D39 Δ*cps rpsL1 rodA*-$L_0$-*ht*-$P_C$-*erm* (IU1824 x *rodA*-$L_0$-*ht*-$P_C$-*erm* fusion amplicon) | Str$^R$ Erm$^R$ | This study |
| IU15906 | D39 Δ*cps rpsL1 mreC*-$L_0$-*ht*-$P_c$-*erm* (IU1824 x *mreC*-$L_0$-*ht*-$P_c$-*erm* fusion amplicon) | Erm$^R$ | This study |
| IU15907 | D39 Δ*cps rpsL1* $P_c$-[*kan-rpsL*$^+$]-*rodA*$^+$ | Kan$^R$ | (20) |
| IU15928 | D39 Δ*cps rpsL1 iht*-$L_6$-*pbp2b* markerless | Str$^R$ | (18) |
| IU15970 | D39 Δ*cps rpsL1 iht*-$L_6$-*rodA* | Str$^R$ | This study |
| IU15997 | D39 Δ*cps rpsL1 iht*-$L_6$-*mpgA* markerless (IU8980 x *iht*-$L_6$-*mpgA* fusion amplicon) | Str$^R$ | This study |
| IU16044 | D39 Δ*cps rpsL1 ftsZ*-$L_2$-*sfgfp* $P_c$-[*kan-rpsL*$^+$]-*pbp2b*$^+$ (IU9985 x $P_c$-[*kan-rpsL*$^+$]-*pbp2b*$^+$ amplicon from IU9023) | Kan$^R$ | This study |
| IU16054 | D39 Δ*cps rpsL1 rodA*$^+$ // Δ*bgaA*::*tet*-$P_{Zn}$-*rodA*$^+$ (IU1824 x Δ*bgaA*::*tet*-$P_{Zn}$-*rodA*$^+$ amplicon from IU10922) | Str$^R$ Tet$^R$ | This study |
| IU16056 | D39 Δ*cps rpsL1 ftsZ*-$L_2$-*sfgfp iht*-$L_6$-*pbp2b* (IU16044 x *iht*-$L_6$-*pbp2b* amplicon from IU15928) | Str$^R$ | This study |
| IU16078 | D39 Δ*cps rpsL1 iht*-$L_6$-*pbp2b* Δ*murZ*::$P_c$-[*kan-rpsL*$^+$] (IU15928 x Δ*murZ*::$P_c$-[*kan-rpsL*$^+$] amplicon from K767) | Kan$^R$ | This study |
| IU16080 | D39 Δ*cps rpsL1 iht*-$L_6$-*rodA* Δ*murZ*::$P_c$-[*kan-rpsL*$^+$] (IU15970 x Δ*murZ*::$P_c$-[*kan-rpsL*$^+$] amplicon from K767) | Kan$^R$ | This study |
| IU16091 | D39 Δ*cps rpsL1 iht*-$L_6$-*pbp2b* Δ*bgaA*::*tet*-$P_{Zn}$-*ftsZ*(D214A) (IU15928 x Δ*bgaA*::*tet*-$P_{Zn}$-*ftsZ*(D214A) amplicon from IU14850) | Str$^R$ Tet$^R$ | This study |
| IU16110 | D39 Δ*cps rpsL1 iht*-$L_6$-*pbp2b* Δ*murZ* (IU16078 x Δ*murZ* amplicon from IU13536) | Str$^R$ | This study |

| Strain number | Genotype (construction)[a b] | Antibiotic resistance[c] | Reference or source |
|---|---|---|---|
| IU16112 | D39 Δ*cps rpsL1 iht*-$L_6$-*rodA* Δ*murA1*<br>(IU16080 x Δ*murA1* amplicon from IU13536) | $Str^R$ | This study |
| IU16134 | D39 Δ*cps rpsL1* Δ*rodA*::$P_c$-[*kan-rpsL*$^+$] // Δ*bgaA*::*tet*-$P_{Zn}$-*rodA*$^+$<br>(IU16054 x Δ*rodA*::$P_c$-[*kan-rpsL*$^+$] amplicon from IU10945) | $Kan^R$ $Tet^R$ | This study |
| IU16136 | D39 Δ*cps rpsL1* Δ*rodA*::$P_c$-*cat* // Δ*bgaA*::*tet*-$P_{Zn}$-*rodA*$^+$<br>(IU16054 x Δ*rodA*::$P_c$-*cat* fusion amplicon) | $Cm^R$ $Str^R$ $Tet^R$ | This study |
| IU16182 | D39 Δ*cps rpsL1* $P_c$-[*kan-rpsL*$^+$]-*rodA*$^+$ Δ*pbp2b*<>*aad9* // Δ*bgaA*::*tet*-$P_{Zn}$-*pbp2b*$^+$<br>(IU11258 x $P_c$-[*kan-rpsL*$^+$]-*rodA*$^+$ amplicon from IU15907) | $Kan^R$ $Tet^R$ $Spc^R$ | This study |
| IU16184 | D39 Δ*cps rpsL1* $P_c$-[*kan-rpsL*$^+$]-*pbp2b*$^+$ Δ*rodA*::$P_c$-*cat* // Δ*bgaA*::*tet*-$P_{Zn}$-*rodA*$^+$<br>(IU16136 x $P_c$-[*kan-rpsL*$^+$]-*pbp2b*$^+$ amplicon from IU9023) | $Kan^R$ $Cm^R$ $Tet^R$ | This study |
| IU16202 | D39 Δ*cps rpsL1 iht-$L_6$-rodA* Δ*pbp2b*<>*aad9* // Δ*bgaA*::*tet*-$P_{Zn}$-*pbp2b*$^+$<br>(IU16182 x *iht-$L_6$-rodA* amplicon from IU15970) | $Str^R$ $Spc^R$ $Tet^R$ | This study |
| IU16204 | D39 Δ*cps rpsL1 iht*-$L_6$-*pbp2b* Δ*rodA*::$P_c$-*cat* // Δ*bgaA*::*tet*-$P_{Zn}$-*rodA*$^+$<br>(IU16184 x *iht*-$L_6$-*pbp2b* amplicon from IU15928) | $Str^R$ $Cm^R$ $Tet^R$ | This study |
| IU16232 | D39 Δ*cps rpsL1 iht*-$L_6$-*pbp2b*(S391A) // Δ*bgaA*::*tet*-$P_{Zn}$-*pbp2b*$^+$<br>(IU15419 x *iht*-$L_6$-*pbp2b*(S391A) fusion amplicon) | $Str^R$ $Tet^R$ | This study |
| IU16236 | D39 Δ*cps rpsL1 rodA*(D283A) // *ΔbgaA::tet-$P_{Zn}$-rodA*$^+$<br>(IU16134 x *rodA*(D283A) fusion amplicon) | $Str^R$ $Tet^R$ | This study |
| IU16239 | D39 Δ*cps rpsL1 iht*-$L_6$-*rodA*(D283A)// *ΔbgaA*::*tet*-$P_{Zn}$-*rodA*$^+$<br>(IU16134 x *iht-$L_6$-rodA*(D283A) fusion amplicon) | $Str^R$ $Tet^R$ | This study |
| IU16252 | D39 Δ*cps rpsL1* [*kan-rpsL*$^+$]-*pbp2b*$^+$ Δ*mreC* // Δ*bgaA::tet*-$P_{Zn}$-*mreC*$^+$<br>(IU12345 x [*kan-rpsL*$^+$]-*pbp2b*$^+$ amplicon from IU9023) | $Kan^R$ $Tet^R$ | This study |

| Strain number | Genotype (construction)[a b] | Antibiotic resistance[c] | Reference or source |
|---|---|---|---|
| IU16281 | D39 Δ*cps rpsL1 iht*-L$_6$-*pbp2b* Δ*mreC //* Δ*bgaA::tet*-P$_{Zn}$-*mreC*$^+$<br>(IU16252 x *iht*-L$_6$-*pbp2b* amplicon from IU15928) | Str$^R$ Tet$^R$ | This study |
| IU16320 | D39 Δ*cps rpsL1 iht*-L$_6$-*pbp1a*<br>(IU6726 x *iht*-L$_6$-*pbp1a* fusion amplicon) | Str$^R$ | This study |
| IU16344 | D39 Δ*cps rpsL1 iht*-L$_6$-*mreC*<br>(IU10103 x *iht*-L$_6$-*mreC* fusion amplicon*)* | Str$^R$ | This study |
| IU16496 | D39 Δ*cps rpsL1 iht*-L$_6$-*rodA* // Δ*bgaA*::*kan*-P$_{Zn}$-*iht*-L$_6$-*rodA*<br>(IU15970 x Δ*bgaA*::*kan*-P$_{Zn}$-*iht*-L$_6$-*rodA* fusion amplicon) | Str$^R$ Kan$^R$ | This study |
| IU16497 | D39 Δ*cps rpsL1 iht*-L$_6$-*pbp1a* // Δ*bgaA*::*kan*-P$_{Zn}$-*iht*-L$_6$-*pbp1a*<br>(IU16320 x Δ*bgaA*::*kan*-P$_{Zn}$-*iht*-L$_6$-*pbp1a* fusion amplicon) | Str$^R$ Kan$^R$ | This study |
| IU16553 | D39 Δ*cps rpsL1 iht*-L$_6$-*pbp2b* // Δ*bgaA*::*kan*-P$_{Zn}$-*iht*-L$_6$-*pbp2b* | Str$^R$ Kan$^R$ | (18) |
| IU17594 | D39 Δ*cps rpsL1 mpgA*(Y488D) Δ*pbp2b*<>*aad9* // *ftsZ*$^+$-P$_c$-[*kan-rpsL*$^+$]<br>(IU9783 x *ftsZ*$^+$-P$_c$-[*kan-rpsL*$^+$] amplicon from IU7614) | Spc$^R$ Kan$^R$ | This study |
| IU17603 | D39 Δ*cps rpsL1 mpgA*(Y488D) Δ*pbp2b*<>*aad9 ftsZ*-L$_2$-*sfgfp*<br>(IU17594 x *ftsZ*-L$_2$-*sfgfp* amplicon from IU9985) | Spc$^R$ Str$^R$ | This study |
| IU18410 | D39 Δ*cps rpsL1 mpgA*(Y488D) Δ*pbp2b*<>*aad9 ftsZ*-L$_2$-*sfgfp* Δ*bgaA*::*kan*-P$_{Zn}$-*iht*-L$_6$-*pbp1a*<br>(IU17603 x Δ*bgaA*::*kan*-P$_{Zn}$-*iht*-L$_6$-*pbp1a* amplicon from IU16497) | Spc$^R$ Kan$^R$ Str$^R$ | This study |
| IU19018 | D39 Δ*cps rpsL1* Δ*murZ*::P$_C$-*erm iht*-L$_6$-*pbp1a* // Δ*bgaA*::*kan*-P$_{Zn}$-*iht*-L$_6$-*pbp1a*<br>(IU16497 x Δ*murZ*::P$_C$-*erm* amplicon from E767) | Erm$^R$ Kan$^R$ Str$^R$ | This study |
| IU19110 | D39 Δ*cps rpsL1 iht-pbp1a*(S370A)<br>(IU6726 x *iht-pbp1a*(S370A) fusion amplicon) | Str$^R$ | This study |
| IU19168 | D39 Δ*cps rpsL1 iht*-L$_6$-*pbp1a*(S370A) // Δ*bgaA*::*kan*-P$_{Zn}$-*iht*-L$_6$-*pbp1a*(S370A)<br>(IU19110 x Δ*bgaA*::*kan*-P$_{Zn}$-*iht*-L$_6$-*pbp1a*(S370A) fusion amplicon) | Kan$^R$ Str$^R$ | This study |
| E193 | D39 Δ*cps* Δ*pbp1b*::P$_c$-*erm* | Erm$^R$ | (21) |
| E767 | D39 Δ*cps* Δ*murZ*::P$_c$-*erm* | Erm$^R$ | (24) |
| K49 | D39 Δ*cps* Δ*mreC*::P$_c$-[*kan-rpsL*$^+$] | Kan$^R$ | (20) |

| Strain number | Genotype (construction)[a b] | Antibiotic resistance[c] | Reference or source |
|---|---|---|---|
| K767 | D39 Δ*cps* Δ*murZ*::$P_c$-[*kan-rpsL*$^+$] | Kan$^R$ | (11) |

[a]Amino-acid sequences of linkers: $L_0$ (GSAGSAAGSG); $L_1$ (LEGSG); $L_2$ (KLDIEFLQ); $L_6$ (LEGSGQGPGSGQGSG). Linker DNA sequences are codon optimized for *Spn* as described previously (11).

[b]$P_c$-*erm* and $P_c$-[*kan-rpsL*$^+$] cassettes described previously (26).

[c]Antibiotic resistance markers: Erm$^R$, erythromycin; Kan$^R$, kanamycin; Spc$^R$, spectinomycin; Str$^R$, streptomycin; Cm$^R$, chloramphenicol; Tet$^R$, tetracycline.

**Table S2.** Oligonucleotide primers used for construction of *S. pneumoniae* strains

| Primer | Sequence (5' to 3') | Template[a] | Amplicon Produced |
|---|---|---|---|
| **For construction of IU15419 (Δ*pbp2b::*$P_c$-[*kan-rpsL*$^+$])** | | | |
| TT452 | GGAGGGTTGGCTGTGGGTGGCTACAAGAAC | D39 | Upstream of *pbp2b* + 60 bp 5' *pbp2b* |
| TT1173 | CATTATCCATTAAAAATCAAACGGATCCTATAA ATTAAGCCGAATCGGAATCGAA | | |
| kanrpsL forward | TAGGATCCGTTTGATTTTTAATGGATAATG | $P_c$-[*kan-rpsL*$^+$] cassette | $P_c$-[*kan-rpsL*$^+$] |
| kanrpsL reverse | GGGCCCCTTTCCTTATGCTTTTG | | |
| TT1174 | CAAAAGCATAAGGAAAGGGGCCCCTAACAAAT GGTGTAGGACCTTCCATTG | D39 | 72 bp of 3' *pbp2b* + downstrea m *pbp2b* |
| TT352 | TGAAGGACTGGAAAGACCACTGCACCTTCT | | |
| **For construction of IU15433 (*rodA*-$L_0$-*ht*-$P_c$-*erm*)** | | | |
| TT494 | TAGGCTTGGGACTTATGATCTTGCCGATTGT | D39 | 5' flanking region |
| TT504 | CGGAGCCAGCGGAACCTTTAATTTGTTTTAATA CAACCTTTTTCCGTT | | |
| TT505 | GGAAAAAGGTTGTATTAAAACAAATTAAAGGTT CCGCTGGCTCCGCT | IU14290 | Middle: $L_0$-*ht*-$P_c$-*erm* |
| TT498 | CTACTTTTACCATGATTTTCTCCTTATTTCCTCC CGTTAAATAATAGATAACTATTAAAA | | |
| TT499 | TTATCTATTATTTAACGGGAGGAAATAAGGAGA AAATCATGGTAAAAGTAGCAGTTATG | D39 | 3' flanking region |
| TT495 | GACGGCCACCTTGATTTAAAATGTCTTCC | | |
| **For construction of IU15906 (*mreC*-$L_0$-HT-$P_c$-*erm*)** | | | |
| TT406 | GCGTGGCTATCAGGGTGGAAAAGGCAGAA | D39 | 5' flanking region |
| ML89 | GCCAGAACCAGCAGCGGAGCCAGCGGAACCT GAATTCCCCACTAATTCTATCACATCTAC | | |
| JQ179 | GGTTCCGCTGGCTCCGCTGCTGGTTCTGGC | IU14290 | $L_0$-*ht*-$P_c$-*erm* |
| JQ184 | TTATTTCCTCCCGTTAAATAATAGATAACTAT | | |
| ML90 | AGTTATCTATTATTTAACGGGAGGAAATAATGA GACAGTTGAAGCGAGTTGGAGTATTTT | D39 | 3' flanking region |
| P107 | CAAAAGCATAAGGAAAGGGGCCCCTGGATGCT GGTAATCGAACAGAGG | | |
| **For construction of IU15997 (*iht*-$L_6$-*mpgA*)** | | | |
| P1348 | TCTTCTTGCAGCCTTGAAAGAGGTGGCAGT | D39 | Upstream of *mpgA* |
| AJP460 | CAAAAAAATTTCCAAACCTTTTTTATCCATAAGT TTTTCCTCCTTGTTGATAATCCATAA | | |
| AJP405 | GATAAAAAAGGTTTGGAAATTTTTTTGGCTTCT GCTGAAATTGGTACTGGTTTTCCATTT | IU14738 | *iht*-$L_6$ |
| YT104 | ACCAGAACCTTGACCAGATCCTGGTCCTTG | | |

| Primer | Sequence (5' to 3') | Template[a] | Amplicon Produced |
|---|---|---|---|
| AJP461 | CAAGGACCAGGATCTGGTCAAGGTTCTGGTTT GAGTGAAAAGTCAAGAGAAGAAGAGAAA | D39 | *mpgA*' |
| TT412 | CGACCAAGGAAGCAATGGTCAACAACTCAT | | |
| **For construction of IU16136 (Δ*rodA*::$P_c$-*cat*)** | | | |
| P1543 | CAGGCCGTACTCTTCTGTCCTCTTTACTTCC | D39 | 5' flanking upstream of *rodA* |
| P1545 | CATTATCCATTAAAAATCAAACGGATCCTAAGC CACCACACCGATGACCA | | |
| kanrpsL forward | TAGGATCCGTTTGATTTTTAATGGATAATG | IU11119[b] | $P_c$-*cat* |
| kanrpsL reverse | GGGCCCCTTTCCTTATGCTTTTG | | |
| P1546 | CAAAAGCATAAGGAAAGGGGCCCTCGATGAGT TACCAGACTAATCTAGCTGAA | D39 | 3' flanking downstream of *rodA* |
| P1544 | CGGGTGTTCAAGCTCTCTGGCTTCATTTTC | | |
| **For construction of IU16232 (*iht*-$L_6$-*pbp2b*(S391A))** | | | |
| TT452 | GGAGGGTTGGCTGTGGGTGGCTACAAGAAC | IU15928 | upstream *iht*-$L_6$-*pbp2b* (S391A) |
| AJP462 | TGGTCGCCGCCTTGACAACAGCACCTGGAACA AAGACATTGGT | | |
| AJP463 | ACCAATGTCTTTGTTCCAGGTGCTGTTGTCAAG GCGGCGACCAT | D39 | *pbp2b* (S391A) to downstream |
| TT352 | TGAAGGACTGGAAAGACCACTGCACCTTCT | | |
| **For construction of IU16236 (*rodA*(D283A))** | | | |
| P1543 | CAGGCCGTACTCTTCTGTCCTCTTTACTTCC | D39 | Upstream to *rodA* (D283A) |
| AJP464 | GCAATAACCGTAAAAATCATAGCTGACTCTCGA ACTGGGATAAGCA | | |
| AJP465 | TTATCCCAGTTCGAGAGTCAGCTATGATTTTTA CGGTTATTGCAG | D39 | *rodA* (D283A) to downstream |
| P1544 | CGGGTGTTCAAGCTCTCTGGCTTCATTTTC | | |
| **For construction of IU16239 (*iht*-$L_6$-*rodA*(D283A))** | | | |
| P1543 | CAGGCCGTACTCTTCTGTCCTCTTTACTTCC | IU15970 | Upstream to *iht*-$L_6$-*rodA* (D283A) |
| AJP464 | GCAATAACCGTAAAAATCATAGCTGACTCTCGA ACTGGGATAAGCA | | |
| AJP465 | TTATCCCAGTTCGAGAGTCAGCTATGATTTTTA CGGTTATTGCAG | D39 | *iht*-$L_6$-*rodA* (D283A) to downstream |
| P1544 | CGGGTGTTCAAGCTCTCTGGCTTCATTTTC | | |
| **For construction of IU16320 (*iht*-$L_6$-*pbp1a*)** | | | |
| P234 | CCCTTGTGTTCATAGCGAGGATAAGCA | D39 | |

| Primer | Sequence (5' to 3') | Template[a] | Amplicon Produced |
|---|---|---|---|
| TT1218 | AAAAATTTCCAAACCTTTTTATCCATCATCTTGTTTTACCACCTAATAAATGTTCTTTG | | Upstream of *pbp1a* |
| TT1219 | AGAACATTTATTAGGTGGTAAAACAAGATGATGGATAAAAAAGGTTTGGAAATTTTTTTG | IU14738 | *iht*-$L_6$ |
| TT1239 | TAGGCGCAGAATCGTTGGTTTGTTACCAGAACCTTGACCAGATCCTGGTCCTTG | | |
| AJP399 | CAAGGACCAGGATCTGGTCAAGGTTCTGGTAACAAACCAACGATTCTGCGCCTA | D39 | *pbp1a* and downstream |
| P235 | AGGCAAGCCTGCAACCATGGTCTTGAAA | | |
| **For construction of IU16344 (*iht*-$L_6$-*mreC*)** | | | |
| P104 | AATGAGACGTGTTGCCATTGCAGG | D39 | Upstream of *mreC* |
| TT1232 | AAAAATTTCCAAACCTTTTTATCCATATCCCTACCTTTATATCAAAAACTGTTACAGTA | | |
| TT1233 | TGTAACAGTTTTTGATATAAAGGTAGGGATATGGATAAAAAAGGTTTGGAAATTTTTTTG | IU14738 | *iht*-$L_6$ |
| TT1234 | GACATATTTTGATTTTTTAAAACGGTTACCAGAACCTTGACCAGATCCTGGTCCTTGTCC | | |
| TT1235 | ACCAGGATCTGGTCAAGGTTCTGGTAACCGTTTTAAAAAATCAAAATATGTCATTATTGT | D39 | *mreC* + *mreD*' |
| TT1236 | CCAAGCCTATAACAAAACAATAGACTAGGTAGAGATACTCTG | | |
| **For construction of IU16496 (Δ*bgaA*::*kan*-$P_{Zn}$-*iht*-$L_6$-*rodA*)** | | | |
| P146 | TGGCCATTCATCGCTGGTCGTGCTGAAAT | IU13393[c] | 5' Δ*bgaA*::*kan*-$P_{Zn}$ |
| TT1260 | AAATTTCCAAACCTTTTTATCCATTACATCGCTTCCTCTCTATCTTCCTTGTTA | | |
| TT1261 | GGAAGATAGAGAGGAAGCGATGTAATGGATAAAAAAGGTTTGGAAATTTTTTTG | IU15970 | *iht*-$L_6$-*rodA* |
| TT897 | AACTGGTTTATGAGAAAGTAAGTTCTTTTATTTAATTTGT TTTAATACAACCTTTTTCCG | | |
| TT898 | AAAAGGTTGTATTAAAACAAATTAAATAAAAGAACTTACT TTCTCATAAACCAGTTGCTG | D39 | 3' *bgaA locus* |
| CS121 | GCTTTCTTGAGGCAATTCACTTGGTGC | | |
| **For construction of IU16497 (Δ*bgaA*::*kan*-$P_{Zn}$-*iht*-$L_6$-*pbp1a*)** | | | |
| P146 | TGGCCATTCATCGCTGGTCGTGCTGAAAT | IU13393[c] | 5' Δ*bgaA*::*kan*-$P_{Zn}$ |
| TT1260 | AAATTTCCAAACCTTTTTATCCATTACATCGCTTCCTCTCTATCTTCCTTGTTA | | |
| TT1261 | GGAAGATAGAGAGGAAGCGATGTAATGGATAAAAAAGGTTTGGAAATTTTTTTG | IU16320 | *iht*-$L_6$-*pbp1a* |
| BR64 | CAGCAACTGGTTTATGAGAAAGTAAGTTCTTTTATGGTTGTGCTGGTTGAGGATTCTG | | |
| BR63 | GAATCCTCAACCAGCACAACCATAAAAGAACTTACTTTCTCATAAACCAGTTGCTGC | D39 | 3' *bgaA locus* |

| Primer | Sequence (5' to 3') | Template[a] | Amplicon Produced |
|---|---|---|---|
| CS121 | GCTTTCTTGAGGCAATTCACTTGGTGC | | |
| **For construction of IU17594 (*ftsZ*$^+$-$P_c$-[*kan-rpsL*$^+$])** | | | |
| TT165 | AGTGGTGCCGATATGGTCTTCATCACTGCT | IU7614 | *ftsZ*$^+$-$P_c$-[*kan-rpsL*$^+$] |
| TT166 | TCATTGGGAGAGCCGGTTCCTGTGAAGAAT | | |
| **For construction of IU17603 (*ftsZ*-L2-*sfgfp*)** | | | |
| TT165 | AGTGGTGCCGATATGGTCTTCATCACTGCT | IU9985 | *ftsZ*-L2-*sfgfp* |
| TT166 | TCATTGGGAGAGCCGGTTCCTGTGAAGAAT | | |
| **For construction of IU18410 (Δ*bgaA*::*kan*-$P_{Zn}$-*iht*-$L_6$-*pbp1a*)** | | | |
| TT657 | CGCCCCAAGTTCATCACCAATGACATCAAC | IU16497 | Δ*bgaA*::*kan*-$P_{Zn}$-*iht*-$L_6$-*pbp1a* |
| CS121 | GCTTTCTTGAGGCAATTCACTTGGTGC | | |
| **For construction of IU19018 (Δ*murZ*::$P_C$-*erm*)** | | | |
| P1554 | GATTTTGTGGTACGACGGGCATGTATAGCG | E767 | Δ*murZ*::$P_C$-*erm* |
| P1555 | TGAACCTGAAATCCCCCTGTAACCAGAACT | | |
| **For construction of IU19110 (*iht-pbp1a*(S370A))** | | | |
| P234 | CCCTTGTGTTCATAGCGAGGATAAGCA | IU16320 | 5' *iht-pbp1a* (S370A) |
| TT1529 | CTGTGATCGGTTTCATAGTTGCTCCCCAGTCGCGGTTTGT | | |
| TT1530 | ACAAACCGCGACTGGGGAGCAACTATGAAACCGATCACAG | D39 | 3' *iht-pbp1a* (S370A) |
| TT627 | TTGTTGTGACAGATGAGTCTGGCAAGGAGC | | |
| **For construction of IU19168 (Δ*bgaA*::*kan*-$P_{Zn}$-*iht-pbp1a*(S370A))** | | | |
| TT657 | CGCCCCAAGTTCATCACCAATGACATCAAC | IU16497 | 5' Δ*bgaA*::*kan*-$P_{Zn}$-*iht-pbp1a* (S370A) |
| TT1529 | CTGTGATCGGTTTCATAGTTGCTCCCCAGTCGCGGTTTGT | | |
| TT1530 | ACAAACCGCGACTGGGGAGCAACTATGAAACCGATCACAG | IU16497 | 3' Δ*bgaA*::*kan*-$P_{Zn}$-*iht-pbp1a* (S370A) |
| CS121 | GCTTTCTTGAGGCAATTCACTTGGTGC | | |
| **For transformation assays** | | | |
| P222 | CGTTCGTGTGGCGCTGCTTCAAATTGTT | E193 | Δ*pbp1b*::$P_c$-*erm* |
| P522 | AACGGCAACCACCAAAGGAGAAACCAAGGA | | |
| P104 | AATGAGACGTGTTGCCATTGCAGG | IU1751 | Δ*mreCD* <>*aad9* |
| P107 | TGTCGCTTTCTCAGCAGCAAGACT | | |
| TT452 | GGAGGGTTGGCTGTGGGTGGCTACAAGAAC | IU7397 | Δ*pbp2b* <>*aad9* |
| TT352 | TGAAGGACTGGAAAGACCACTGCACCTTCT | | |
| TT329 | CAACTGATATAGTTGGAAGTGAGGAGTCCATTTCCC | IU9931 | Δ*rodZ* <>*aad9* |
| P1385 | ACAACACCTGCAATGGCCACACGTTGCTTT | | |
| P1543 | CAGGCCGTACTCTTCTGTCCTCTTTACTTCC | IU10943 | Δ*rodA*::$P_c$-*erm* |
| P1544 | CGGGTGTTCAAGCTCTCTGGCTTCATTTTC | | |
| P222 | CGTTCGTGTGGCGCTGCTTCAAATTGTT | IU13680 | Δ*pbp1b*::$P_c$-*aad9* |
| P522 | AACGGCAACCACCAAAGGAGAAACCAAGGA | | |

[a]Genomic DNA of D39W was used as templates for PCR reactions, except for the $P_c$-[*kan-rpsL*$^+$ ] and $P_c$-*erm* cassettes (26).

[b]Genotype of IU11119 is D39 Δ*cps ezrA*-$L_0$-*sfgfp*-$P_c$ –*cat* (11).

[c]Genotype of IU13393 is D39 Δ*cps* Δ*bgaA*::*kan*-$P_{Zn}$-*murZ* (24).

**Table S3.** Cellular amounts of iHT-tagged proteins relative to untagged WT proteins

| Genotype (strain #) | mM Zn inducer | % tagged protein amount relative to untagged WT amount (mean ± SD)[a] | n |
|---|---|---|---|
| *iht-pbp2b* (IU15928) | 0 | 15 ±5 | 3 |
| *iht-pbp2b* // $P_{Zn}$-*iht-pbp2b* (IU16553) | 0 | 13 ±1 | 2 |
| | 0.25 | 82 ±6 | 2 |
| *iht-pbp1a* (IU16320) | 0.25 | 10 ±10 | 2 |
| *iht-pbp1a* // $P_{Zn}$-*iht-pbp1a* (IU16497) | 0 | 9 ±3 | 3 |
| | 0.25 | 101 ±41 | 3 |
| *iht-pbp1a*(S370A) // $P_{Zn}$-*iht-pbp1a*(S370A) (IU19168) | 0 | 4 ±2 | 2 |
| | 0.25 | 78 ±50 | 2 |
| *iht-pbp1a* // $P_{Zn}$-*iht-pbp1a* Δ*murZ* (IU19018) | 0 | 8 ±9 | 2 |
| | 0.25 | 162 ±26 | 2 |
| *mpgA*(Y488D) Δ*pbp2b* *ftsZ-sfgfp pbp1a* // $P_{Zn}$-*iht-pbp1a* (IU18410) | 0 | <1[b] | 3 |
| | 0.25 | 32 ±2[b] | 2 |
| *iht-mreC* (IU16344) | 0 | 82 ±18 | 4 |
| *iht-rodA* (IU15970) | 0 | nd[c] | |
| *iht-rodA* // $P_{Zn}$-*iht-rodA* (IU16496) | 0 | nd[c] | |
| | 0.25 | nd[c] | |
| *iht-mpgA* (IU15997) | 0 | 171 ±11 | 2 |

[a]Determined by quantitative western blotting as described in *Materials and Methods*. WT untagged protein amounts were determined in strain IU1824 and defined as 100%.

[b]iHT-aPBP1a protein amount only; untagged aPBP1a was also expressed (see Fig. S20*C*).

[c]nd, no data due to lack of antibody to native RodA(*Spn*).

**Table S4.** HT-JF549 ligand concentrations used to perform single molecule (sm-) TIRFm

| Protein labeled | Strain | [HT-JF549] (pM) |
|---|---|---|
| iHT-bPBP2b | IU15928 | 100 |
| | IU16553 | 60-100 |
| | IU16091 | 100 |
| | IU16056 | 100 |
| | IU16110 | 100 |
| | IU16232 | 100 |
| | IU16281 | 100 |
| | IU16204 | 120-150 |
| iHT-RodA | IU15970 | 100-120 |
| | IU16496 | 100-120 |
| | IU16112 | 100 |
| | IU16239 | 100 |
| | IU16202 | 120-150 |
| iHT-aPBP1a | IU16320 | 50-100 |
| | IU16497 | 50-100 |
| | IU18410 | 100 |
| | IU19018 | 100 |
| | IU19168 | 100 |
| iHT-MreC | IU16344 | 30-50 |
| iHT-MpgA | IU15997 | 30-60 |

**TABLE S5.** iHT-MpgA is active based on lack of suppression of PG elongasome mutations in transformation assays

| **Amplicon** (30 ng) | **Selection** | **Recipient strain**[a] | | |
|---|---|---|---|---|
| | | D39 Δ*cps rpsL1* (**WT**, IU1824) | ***iht-mpgA*** (IU15997) | ***mpgA*(Y488D)**[b] (IU9760) |
| | | **# of colonies after 22 h** | | |
| Δ*rodA*::$P_c$-*erm* | Erythromycin (0.3 µg/mL) | 0 | 0 | >500 |
| Δ*pbp1b*::$P_c$-*erm* | | >500 | >500 | >500 |
| No DNA | | 0 | 0 | 0 |
| Δ*mreCD<>aad9* | Spectinomycin (150 µg/mL) | 0 | 0 | >500 |
| Δ*pbp2b<>aad9* | | 0 | 0 | >500 |
| Δ*rodZ<>aad9* | | 0 | 0 | >500 |
| Δ*pbp1b*::$P_c$-*aad9* | | >500 | >500 | >500 |
| No DNA | | 0 | 0 | 0 |

[a]Recipient strains in D39 Δ*cps rpsL1* (IU1824) genetic background and amplicons were obtained as described in Table S1 and S2. Transformation assays and visualization of colonies normalized to 1 mL of transformation mixture were performed as described in *Materials and Methods*. All transformation experiments were performed with Δ*pbp1b* amplicons containing the same antibiotic selections as the positive control for detection of colonies. Each transformation experiment was performed 2 times independently with similar results.

[b]Y488D is an MpgA variant previously characterized to be partially defective (6).

### SUPPLEMENTAL LEGENDS FOR MOVIES S1 TO S30

**bPBP2b movies:** S1, S4, S6, S7, S8, S10, S11, S13, S14, S24

**RodA movies:** S2, S5, S9, S12, S15, S25

**MreC movies:** S3, S26

**aPBP1a movies:** S16, S17, S18, S19, S20, S21, S27, S28

**MpgA movies:** S22, S23, S29, S30

**Movie S1.** Single molecules of iHT-bPBP2b displayed processive circumferential motion at midcell. Diffusively moving and static molecules were also observed. Sm-TIRFm was performed on strain IU15928 (*iht-pbp2b*) as described in *Materials and Methods*. White arrow points to a cell with a circumferentially moving molecule that changes direction. A diffusively moving molecule can also be seen in this cell. Cyan arrow points to a cell with a static molecule. Scale bar = 1 μm. Images were taken at 1 s intervals. Movie is shown at 8 FPS.

**Movie S2.** Single molecules of iHT-RodA displayed processive circumferential motion at midcell. Sm-TIRFm was performed on strain IU15970 (*iht-rodA*) as described in *Materials and Methods*. White arrow points to a cell with a circumferentially moving molecule. Scale bar = 1 μm. Images were taken at 1 s intervals. Movie is shown at 8 FPS.

**Movie S3.** Single molecules of iHT-MreC displayed processive circumferential motion at midcell. Diffusively moving and static molecules were also observed. Sm-TIRFm was performed on strain IU16344 (*iht-mreC*) as described in *Materials and Methods*. White arrow points to a cell with a circumferentially moving molecule. Cyan arrow points to a cell with static, diffusive, and circumferentially moving molecules. Scale bar = 1 μm. Images were taken at 1 s intervals. Movie is shown at 8 FPS.

**Movie S4.** When iHT-bPBP2b was expressed near WT levels, ≈70% of molecules displayed diffusive motion. Sm-TIRFm was performed on strain IU16553 (*iht-pbp2b* // $P_{Zn}$-*iht-pbp2b*) in C+Y medium containing 0.25 mM Zn inducer as described in *Materials and Methods*. Cyan arrow points to a cell with a static molecule. White arrows point to cells with circumferentially moving molecules. Diffusive molecules are seen in several cells. Scale bar = 1 μm. Images were taken at 1 s intervals. Movie is shown at 8 FPS.

**Movie S5.** When iHT-RodA was expressed near WT levels, ≈70% of molecules displayed diffusive motion. Sm-TIRFm was performed on strain IU16496 (*iht-rodA* // $P_{Zn}$-*iht-rodA*) in C+Y medium containing 0.25 mM Zn inducer as described in *Materials and Methods*. Scale bar = 1 μm. Images were taken at 1 s intervals. Movie is shown at 8 FPS.

**Movie S6.** Circumferential velocity of iHT-bPBP2b molecules did not decrease when FtsZ(D214A) was overexpressed. Sm-TIRFm was performed on strain IU16091 (*iht-pbp2b* // $P_{Zn}$-*ftsZ*(D214A)) in C+Y medium containing 0.25 mM Zn inducer as described in *Materials and Methods*. Scale bar = 1 μm. Images were taken at 1 s intervals. Movie is shown at 8 FPS.

**Movie S7.** Circumferential velocity of FtsZ-sfGFP (green) was over two-fold faster than that of iHT-bPBP2b (red). Sm-TIRFm was performed on strain IU16056 (*iht-pbp2b ftsZ-sfgfp*) as described in *Materials and Methods*. White arrows denote cells, where both nascent FtsZ-sfGFP rings and circumferentially moving iHT-bPBP2b molecules can be seen. Scale bar = 1 μm. Images were taken at 1 s intervals. Movie is shown at 8 FPS.

**Movie S8.** iHT-bPBP2b circumferential velocity decreased in a Δ*murZ* background. Sm-TIRFm was performed on strain IU16110 (*iht-pbp2b* Δ*murZ*) as described in *Materials* *and Methods*. Scale bar = 1 µm. Images were taken at 1 s intervals. Movie is shown at 8 FPS.

**Movie S9.** iHT-RodA circumferential velocity decreased in a Δ*murZ* background. Sm-TIRFm was performed on strain IU16112 (*iht-rodA* Δ*murZ*) as described in *Materials and Methods*. Scale bar = 1 µm. Images were taken at 1 s intervals. Movie is shown at 8 FPS.

**Movie S10.** iHT-bPBP2b(S391A) molecules were static at midcell or moved diffusively, but did not move circumferentially. Sm-TIRFm was performed on strain IU16232 (*iht-pbp2b*(S391A) // $P_{Zn}$-*pbp2b*) in C+Y medium without Zn as described in *Materials and Methods*. Scale bar = 1 µm. Images were taken at 1 s intervals. Movie is shown at 8 FPS.

**Movie S11.** iHT-bPBP2b(S391A) molecules did not move circumferentially, even in the presence of WT bPBP2b. Sm-TIRFm was performed on strain IU16232 (*iht-pbp2b*(S391A) // $P_{Zn}$-*pbp2b*) in C+Y medium containing 0.25 mM Zn inducer as described in *Materials and Methods*. Scale bar = 1 µm. Images were taken at 1 s intervals. Movie is shown at 8 FPS.

**Movie S12.** iHT-RodA(D283A) molecules were static at midcell or moved diffusively, but did not move circumferentially. Sm-TIRFm was performed on strain IU16239 (*iht-rodA*(D283A) // $P_{Zn}$-*rodA*) in C+Y medium without Zn as described in *Materials and Methods*. Scale bar = 1 µm. Images were taken at 1 s intervals. Movie is shown at 8 FPS.

**Movie S13.** iHT-bPBP2b molecules did not move circumferentially when MreC was depleted. Sm-TIRFm was performed on strain IU16281 (*iht-pbp2b* Δ*mreC* // $P_{Zn}$-*mreC*) after 3 h of growth in C+Y medium lacking Zn as described in *Materials and Methods*. Scale bar = 1 µm. Images were taken at 1 s intervals. Movie is shown at 8 FPS.

**Movie S14.** iHT-bPBP2b molecules shifted largely from circumferential to diffusive movement when RodA was depleted. Sm-TIRFm was performed on strain IU16204 (*iht-pbp2b ∆rodA* // $P_{Zn}$-*rodA*) after 3 h of growth in C+Y medium without Zn as described in *Materials and Methods*. White arrows point to cells with circumferentially moving molecules. Scale bar = 1 µm. Images were taken at 1 s intervals. Movie is shown at 8 FPS.

**Movie S15.** iHT-RodA molecules shifted largely from circumferential to diffusive movement when bPBP2b was depleted. Sm-TIRFm was performed on strain IU16202 (*iht-rodA ∆pbp2b* // $P_{Zn}$-*pbp2b*) after 3 h of growth in C+Y medium without Zn as described in *Materials and Methods*. Scale bar is 1 µm. Images were taken at 1 s intervals. Movie is shown at 8 FPS.

**Movie S16.** iHT-aPBP1a displayed processive circumferential motion at midcell. Sm-TIRFm was performed on strain IU16497 (*iht-pbp1a* // $P_{Zn}$-*iht-pbp1a*) in C+Y containing 0.25 mM Zn inducer as described in *Materials and Methods*. White arrow points to a cell with a circumferentially moving molecule. Scale bar = 1 µm. Images were taken at 1 s intervals. Movie is shown at 8 FPS.

**Movie S17.** iHT-aPBP1a displayed processive circumferential motion at midcell. Sm-TIRFm was performed on strain IU16320 (*iht-pbp1a*) as described in *Materials and Methods*. White arrow points to a cell with a circumferentially moving molecule. Scale bar = 1 µm. Images were taken at 1 s intervals. Movie is shown at 8 FPS.

**Movie S18.** The majority of iHT-aPBP1a molecules moved diffusively, even when underexpressed compared to WT. Sm-TIRFm was performed on strain IU16320 (*iht-*

*pbp1a*) as described in *Materials and Methods*. Scale bar = 1 µm. Images were taken at 1 s intervals. Movie is shown at 8 FPS.

**Movie S19.** Circumferential movement was largely eliminated when aPBP1a was catalytically inactivated. Sm-TIRFm was performed on strain IU19168 (*iht-pbp1a*(S370A)//$P_{Zn}$-*iht-pbp1a*(S370A)) as described in *Materials and Methods*. Scale bar = 1 µm. Images were taken at 1 s intervals. Movie is shown at 8 FPS.

**Movie S20.** iHT-aPBP1a showed similar single-molecule dynamics in a Δ*murZ* mutant and a *murZ*$^+$ strain. Sm-TIRFm was performed on strain IU19018 (*iht-pbp1a*//$P_{Zn}$-*iht-pbp1a* Δ*murZ*) as described in *Materials and Methods*. Scale bar = 1 µm. Images were taken at 1 s intervals. Movie is shown at 8 FPS.

**Movie S21.** iHT-aPBP1a single-molecule dynamics were the same in a *mpgA*(Y488D) Δ*pbp2b* merodiploid strain as in a WT strain. Sm-TIRFm was performed on strain IU18410 (*mpgA*(Y488D) Δ*pbp2b ftsZ-sfgfp pbp1a*//$P_{Zn}$-*iht-pbp1a*) as described in *Materials and Methods*. Scale bar = 1 µm. Images were taken at 1 s intervals. Movie is shown at 8 FPS.

**Movie S22.** Approximately 80% of iHT-MpgA molecules moved subdiffusively and were confined to midcell. Sm-TIRFm was performed on strain IU15997 (*iht-mpgA*) as described in *Materials and Methods*. Scale bar = 1 µm. Images were taken at 1 s intervals. Movie is shown at 8 FPS.

**Movie S23.** iHT-MpgA molecules displayed non-processive motion at midcell. Sm-TIRFm was performed on strain IU15997 (*iht-mpgA*) as described in *Materials and Methods*. Scale bar is 1 µm. Images were taken at 0.1 s intervals. Movie is shown at 10 FPS.

**Movie S24.** Imaging iHT-bPBP2b at 20 FPS revealed processive, circumferentially, moving molecules, static molecules, and rapidly diffusing molecules, consistent with imaging at 1 FPS. Sm-TIRFm was performed on strain IU16553 (*iht-pbp2b* // $P_{Zn}$-*iht-pbp2b*) in C+Y medium containing 0.2 mM Zn inducer as described in *Materials and Methods*. Scale bar = 1 µm. Images were taken at 0.05 s intervals. Movie is shown at 20 FPS.

**Movie S25.** Imaging iHT-RodA at 20 FPS revealed processive, circumferentially, moving molecules, static molecules, and rapidly diffusing molecules, consistent with imaging at 1 FPS. Sm-TIRFm was performed on strain IU16496 (*iht-rodA* // $P_{Zn}$-*iht-rodA*) in C+Y medium containing 0.2 mM Zn inducer as described in *Materials and Methods*. Scale bar = 1 µm. Images were taken at 0.05 s intervals. Movie is shown at 20 FPS.

**Movie S26.** Imaging iHT-MreC at 20 FPS revealed processive, circumferentially, moving molecules, static molecules, and rapidly diffusing molecules, consistent with imaging at 1 FPS. Sm-TIRFm was performed on strain IU16344 (*iht-mreC*) in C+Y medium containing 0.2 mM Zn inducer as described in *Materials and Methods*. Scale bar = 1 µm. Images were taken at 0.05 s intervals. Movie is shown at 20 FPS.

**Movie S27.** Imaging iHT-aPBP1a at 20 FPS revealed processive, circumferentially, moving molecules, static molecules, and rapidly diffusing molecules, consistent with imaging at 1 FPS. Sm-TIRFm was performed on strain IU16497 (*iht-pbp1a* // $P_{Zn}$-*iht-pbp1a*) in C+Y medium containing 0.2 mM Zn inducer as described in *Materials and Methods*. Scale bar = 1 µm. Images were taken at 0.05 s intervals. Movie is shown at 20 FPS.

**Movie S28.** iHT-aPBP1a molecules imaged at 20 FPS transitioned rapidly between diffusive motion and being static at midcell. Sm-TIRFm was performed on strain IU16497 (*iht-pbp1a* // $P_{Zn}$-*iht-pbp1a*) in C+Y medium containing 0.2 mM Zn inducer as described in *Materials and Methods*. Scale bar = 1 µm. Images were taken at 0.05 s intervals. Movie is shown at 20 FPS.

**Movie S29.** iHT-MpgA molecules imaged at 20 FPS were largely confined to midcell. Sm-TIRFm was performed on strain IU15997 (*iht-mpgA*) in C+Y containing 0.2 mM Zn inducer as described in *Materials and Methods*. Scale bar = 1 µm. Images were taken at 0.05 s intervals. Movie is shown at 20 FPS.

**Movie S30.** iHT-MpgA molecules imaged at 20 FPS transitioned rapidly between diffusive motion and being static at midcell. Sm-TIRFm was performed on strain IU15997 (*iht-mpgA*) in C+Y medium containing 0.2 mM Zn inducer as described in *Materials and Methods*. Scale bar = 1 µm. Images were taken at 0.05 s intervals. Movie is shown at 20 FPS.

FtsZ polymers drive the directional movement of sPG-synthesis enzymes via a Brownian ratchet mechanism. *Nat Commun*, **12**, 609.

### SUPPLEMENTAL FIGURES S1 TO S23 WITH LEGENDS

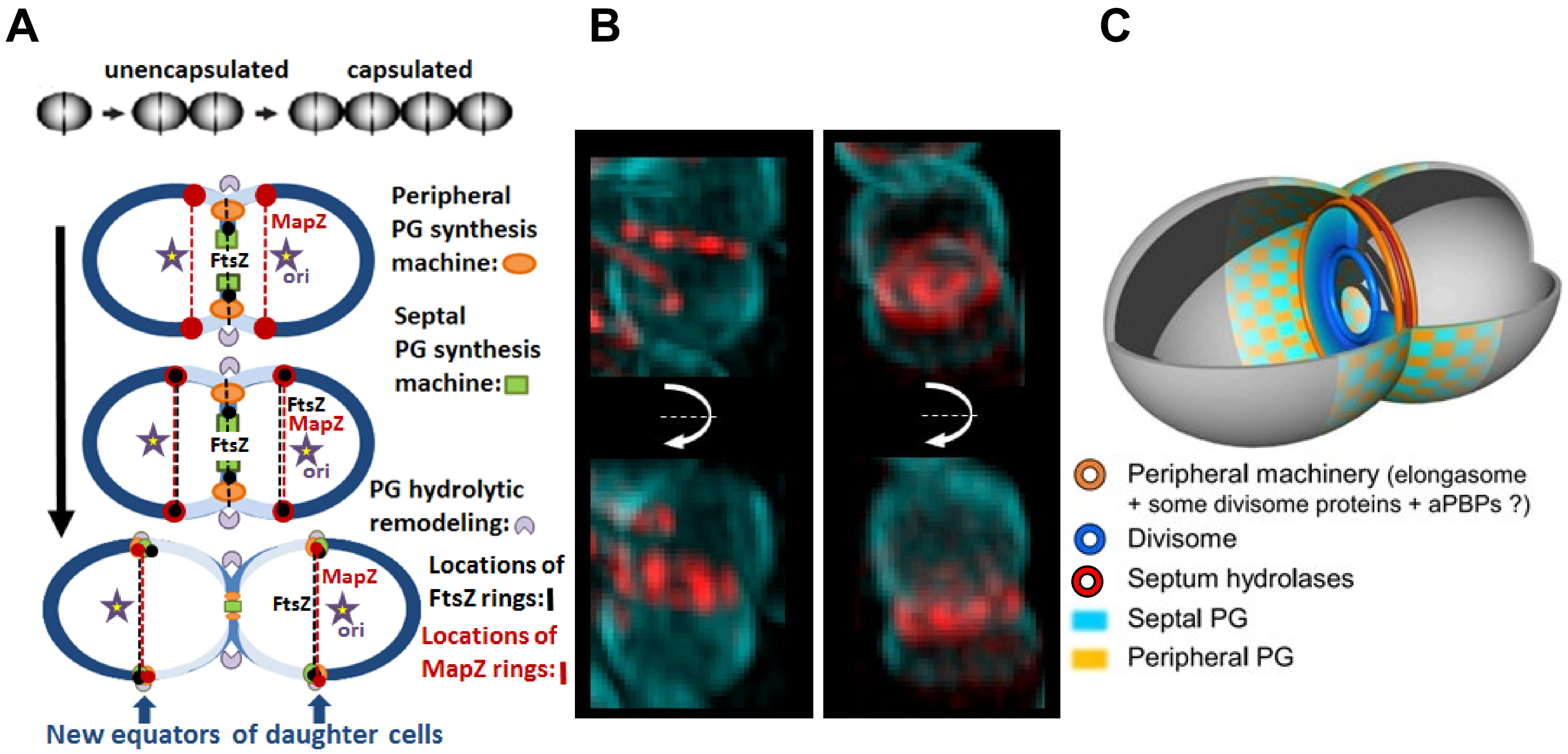
A
unencapsulated
capsulated
Peripheral PG synthesis machine:
Septal PG synthesis machine:
MapZ
FtsZ
ori
PG hydrolytic remodeling:
Locations of FtsZ rings:
Locations of MapZ rings:
New equators of daughter cells
B
C
Peripheral machinery (elongasome + some divisome proteins + aPBPs ?)
Divisome
Septum hydrolases
Septal PG
Peripheral PG

Fig. S1

**Fig. S1.** PG synthesis is confined to a zone at midcell in *Streptococcus pneumoniae*. (*A*) Diagram of a pneumococcal cell throughout cell division. Cell wall shown in blue, with existing PG (dark blue) and newly synthesized PG (light blue). Elongation (peripheral) PG synthesis machine (orange oval); septal PG synthesis machine (green rectangle); PG remodeling hydrolases (light purple circular sector); FtsZ rings (black dotted lines); and MapZ rings (red dotted lines). From reference (11). (*B*) 3D-SIM images of *S. pneumoniae* cells labeled with a long pulse of a blue FDAA (HADA) and a short pulse of a red FDAA (TADA) revealing inner and outer rings of PG synthesis during cell division. Images in the bottom row are rotations of images in the top row. From reference (18). (*C*) Summary model depicting the separation of the septal and elongation (peripheral) PG synthesis machines during cell division. Also shown is remodeling (checkerboard pattern) by PG hydrolases that mixes the PG produced by septal (light blue) and elongation (peripheral) PG synthesis (light orange). Septal annular disk, where leading-edge septal PG synthesis closes the ring (blue); elongation (peripheral) PG synthesis around the outer edge of the annular disk (orange ring); remodeling PG hydrolase, such as FtsEX:PcsB, at the outer edge of the annular disk (red ring). From reference (27).

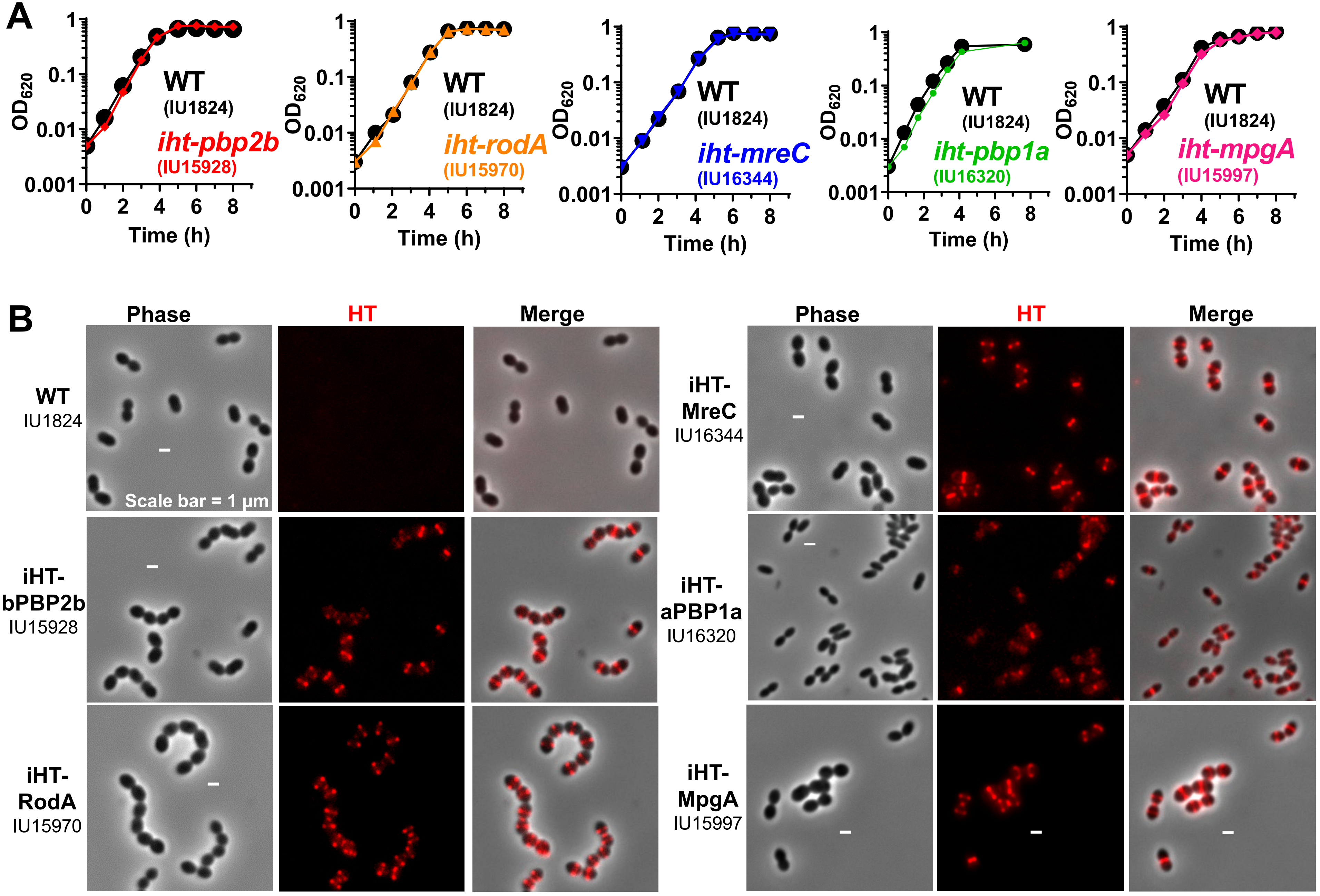
A
OD620
Time (h)
WT
(IU1824)
iht-pbp2b
(IU15928)
iht-rodA
(IU15970)
iht-mreC
(IU16344)
iht-pbp1a
(IU16320)
iht-mpgA
(IU15997)
B
Phase
HT
Merge
WT
IU1824
Scale bar = 1 μm
iHT-
bPBP2b
IU15928
iHT-
RodA
IU15970
iHT-
MreC
IU16344
iHT-
aPBP1a
IU16320
iHT-
MpgA
IU15997

Fig. S2

**Fig. S2.** Growth curve analysis and protein localization indicated fluorescent fusion proteins had nearly full function. (*A*) D39 Δ*cps rpsL1* (WT, black, IU1824); *iht-pbp2b* (red, IU15928); *iht-rodA* (gold, IU15970); *iht-mreC* (blue, IU16344); *iht-pbp1a* (green, IU16320); and *iht-mpgA* (IU15997, magenta) cells were grown in C+Y medium. Representative curves are shown from two biological replicates that gave similar results. (*B*) Cells were labeled with a saturating amount (500 nM) of HT-TMR ligand and viewed by 2D-FM. Representative images are shown from two or more biological replicates that gave similar results. Scale bar = 1 μm.

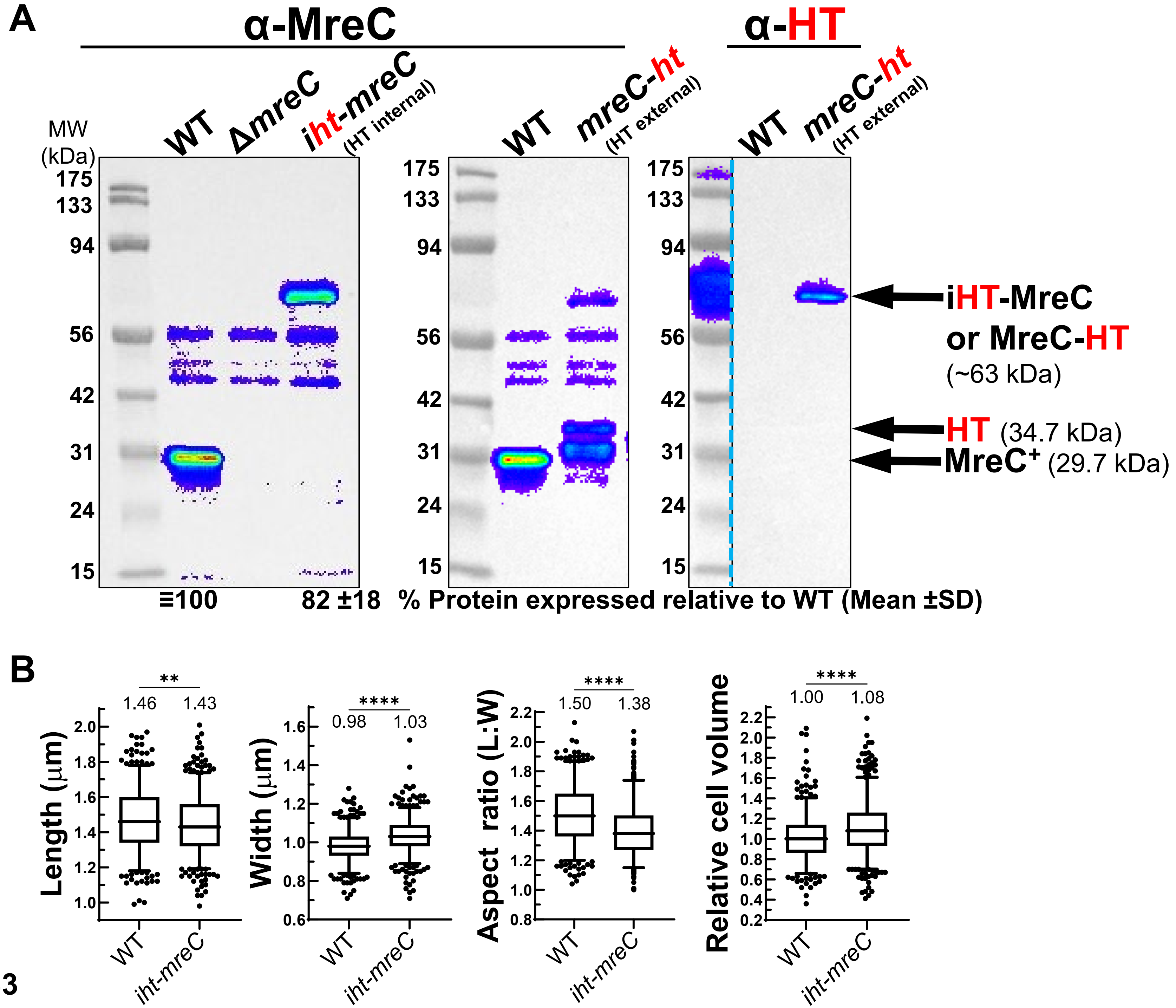
A
α-MreC
α-HT
MW (kDa)
WT
ΔmreC
iht-mreC (HT internal)
WT
mreC-ht (HT external)
WT
mreC-ht (HT external)
175
133
94
56
42
31
24
15
iHT-MreC or MreC-HT (~63 kDa)
HT (34.7 kDa)
MreC+ (29.7 kDa)
≡100
82 ±18
% Protein expressed relative to WT (Mean ±SD)
B
Length (µm)
**
1.46
1.43
Width (µm)
****
0.98
1.03
Aspect ratio (L:W)
****
1.50
1.38
Relative cell volume
****
1.00
1.08
WT
iht-mreC
Fig. S3

**Fig. S3.** Cells expressed iHT-MreC near WT levels and showed minimal morphological differences compared to WT. (*A*) Quantitative western blots were performed as described in *Materials and Methods* on lysates of strains: D39 Δ*cps rpsL1* (WT, IU1824); *mpgA*(Y488D) Δ*mreCD* (Δ*mreC*, IU10651); *iht-mreC* (intracellular HT, IU16344); and *mreC-ht* (extracellular HT, IU15906). Blots were probed with anti-MreC (left and middle) or anti-HT (right) antibody. Representative blots are shown from two biological replicates that gave similar results. Dotted blue line in the right blot denotes that some lanes were cropped from the image for clarity. (*B*) Box and whisker plots (whiskers, 5 and 95 percentile) of cell lengths, widths, aspect ratios (cell length to width), and relative cell volumes. Over 400 cells from three biological replicates were measured. A Mann-Whitney test was used to compare dimension parameters between strains. **$P$ < 0.01; ****$P$ < 0.0001.

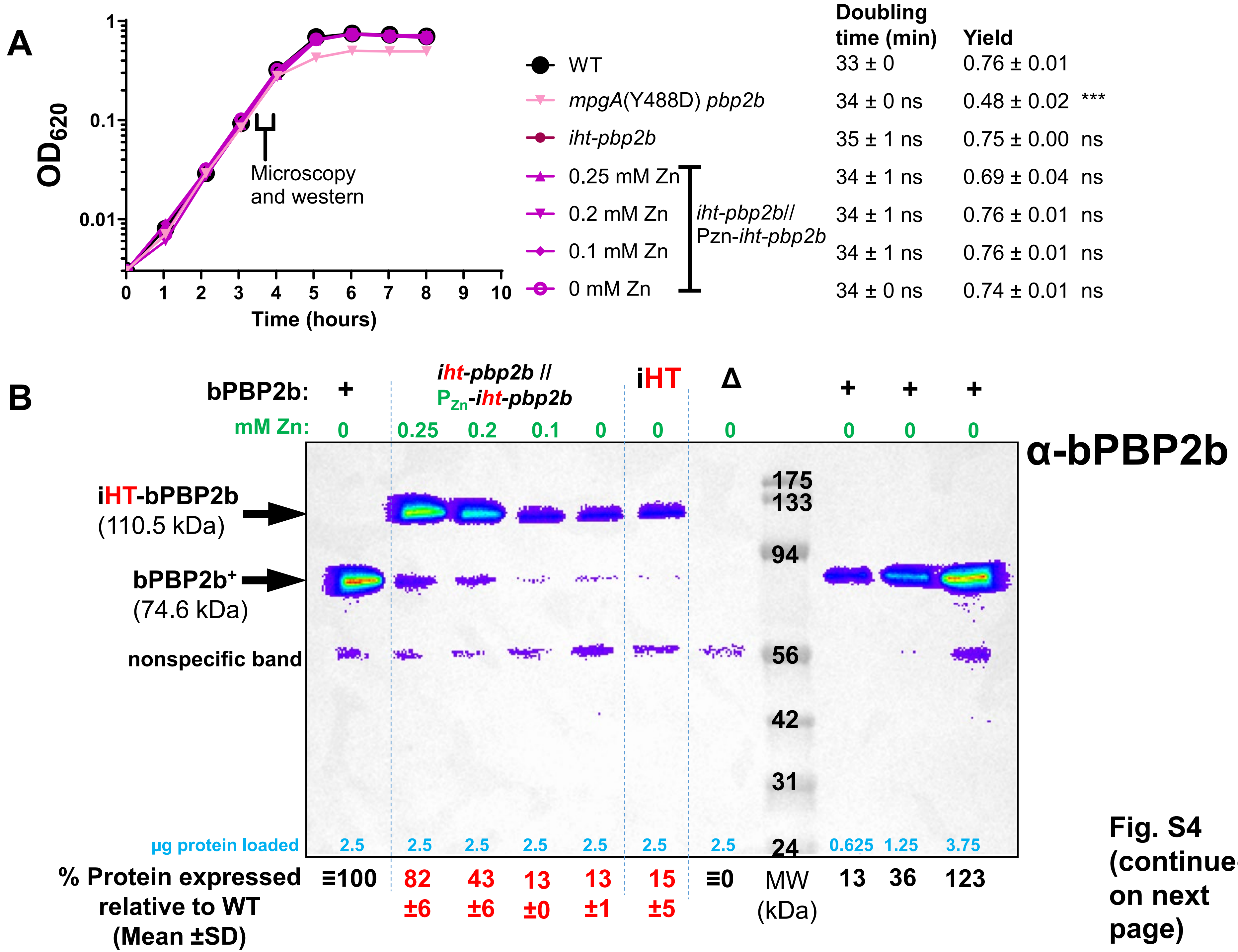


Fig. S4 (continued on next page)

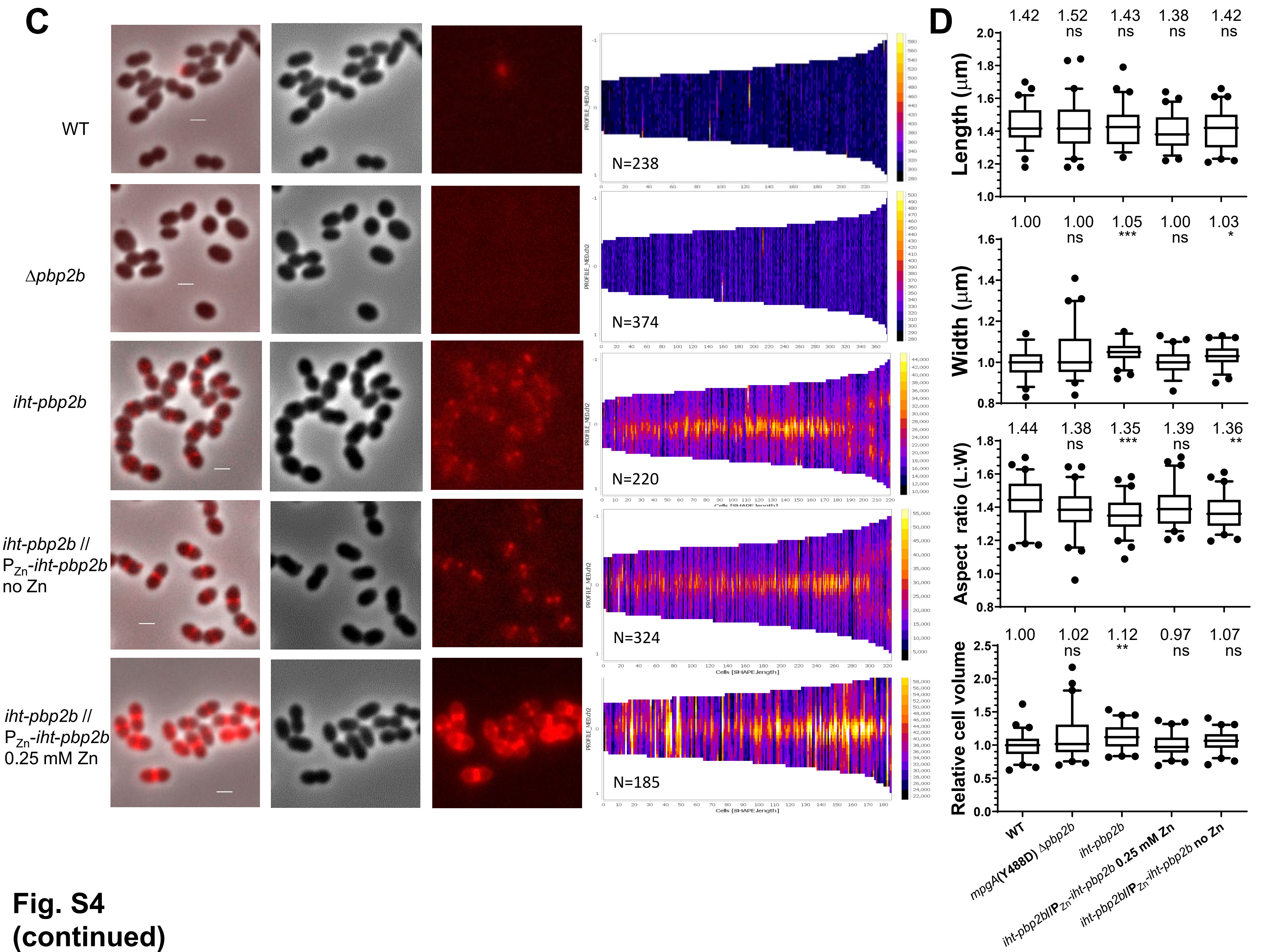


**Fig. S4 (continued)**

**Fig. S4.** iHT- bPBP2b was expressed from the native locus at 15 ± 5% (mean ± SD) of the amount of untagged WT bPBP2b. D39 Δ*cps rpsL1 pbp2b*⁺ (WT, indicated by +; IU1824); *iht-pbp2b* (iHT, IU15928); *iht-pbp2b* // $P_{Zn}$-*iht-pbp2b* (IU16553); and *mpgA*(Y488D) Δ*pbp2b* (indicated by Δ; IU9783) were grown in C+Y medium with the indicated concentrations of Zn inducer, and 2D-FM and western blot analysis were performed as described in *Materials and Methods*. (*A*) Representative growth curves of two or more biological replicates that gave similar results. Times of sample collection for microscopy and western blotting are indicated. The doubling times and growth yields (mean ± SEM) are indicated, and a one-way ANOVA with Dunnet's post-test was used to compare growth parameters among strains compared to WT. *ns* (nonsignificant); ****P* < 0.001. (*B*) Western blots of cells expressing bPBP2b or iHT-bPBP2b probed with anti-bPBP2b antibody. Amounts of added Zn inducer are shown at the top in green; µg total protein loaded are shown toward the bottom in blue; % iHT-bPBP2b expressed relative to untagged WT bPBP2b (mean ± SD) is shown at the bottom. A representative blot is shown from two biological replicates that gave similar results. (*C*) 2D-FM images of cells expressing bPBP2b or iHT-bPBP2b labeled with a saturating amount (500 nM) of HT-ligand. The fluorescence intensity of images for *iht-pbp2b* // $P_{Zn}$-*iht-pbp2b* +0.25 mM Zn inducer (IU16553) was lowered compared to that of the other strains to visualize localization. Scale bar = 1 µm. Demograph intensity scales (shown on the right of the demographs) are auto-adjusted based on the maximal intensity reading and "N" denotes total number of cells analyzed. (*D*) Box and whisker plots (whiskers, 5 and 95 percentile) of cell lengths, widths, aspect ratios (cell length to width), and relative cell volumes. n = 60 cells from two biological replicates. A Kruskal-Wallis ANOVA with Dunn's post-test was used to compare dimension parameters compared to WT values. *ns* (nonsignificant); **P* < 0.05; ***P* < 0.01; *** *P* < 0.001.

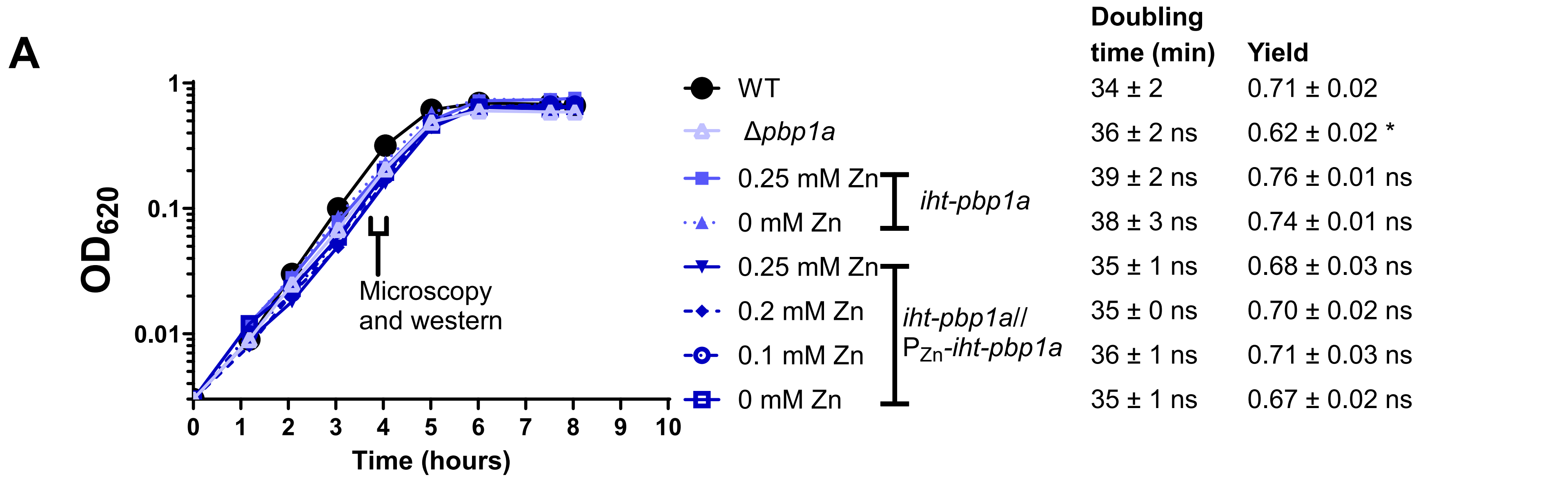


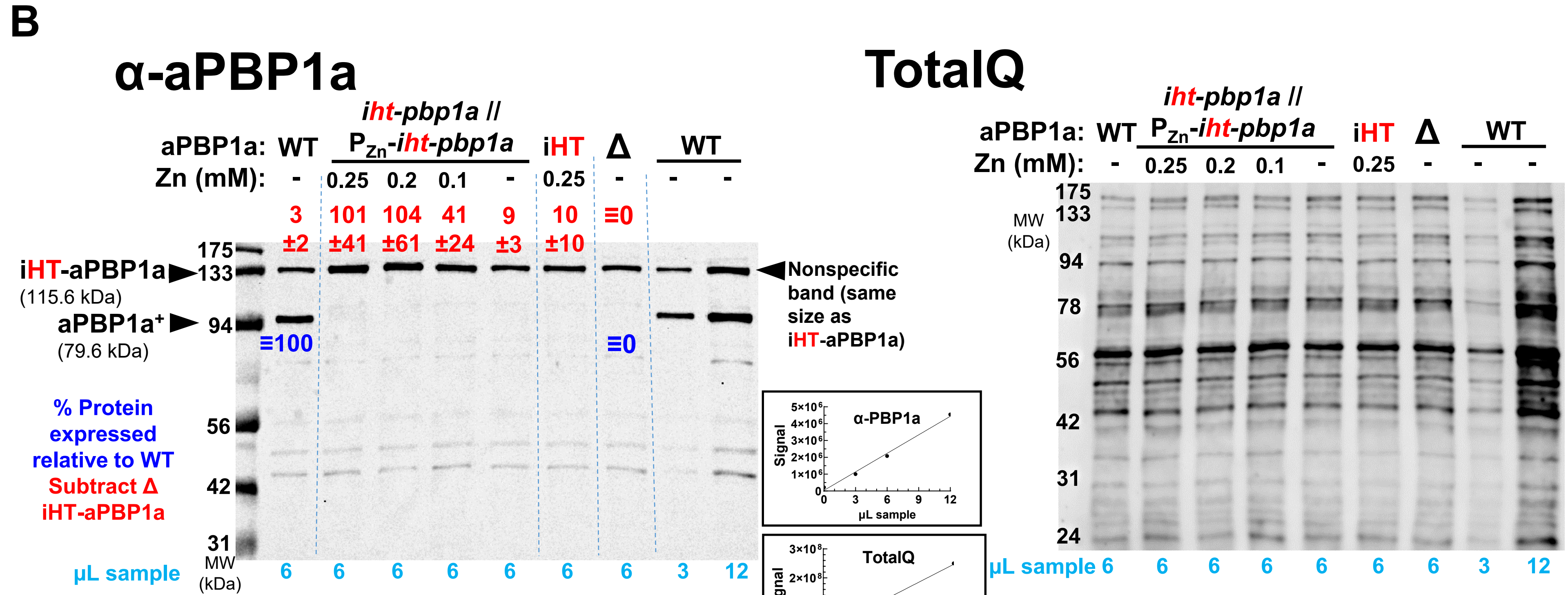


**Fig. S5**
**(continued on next page)**

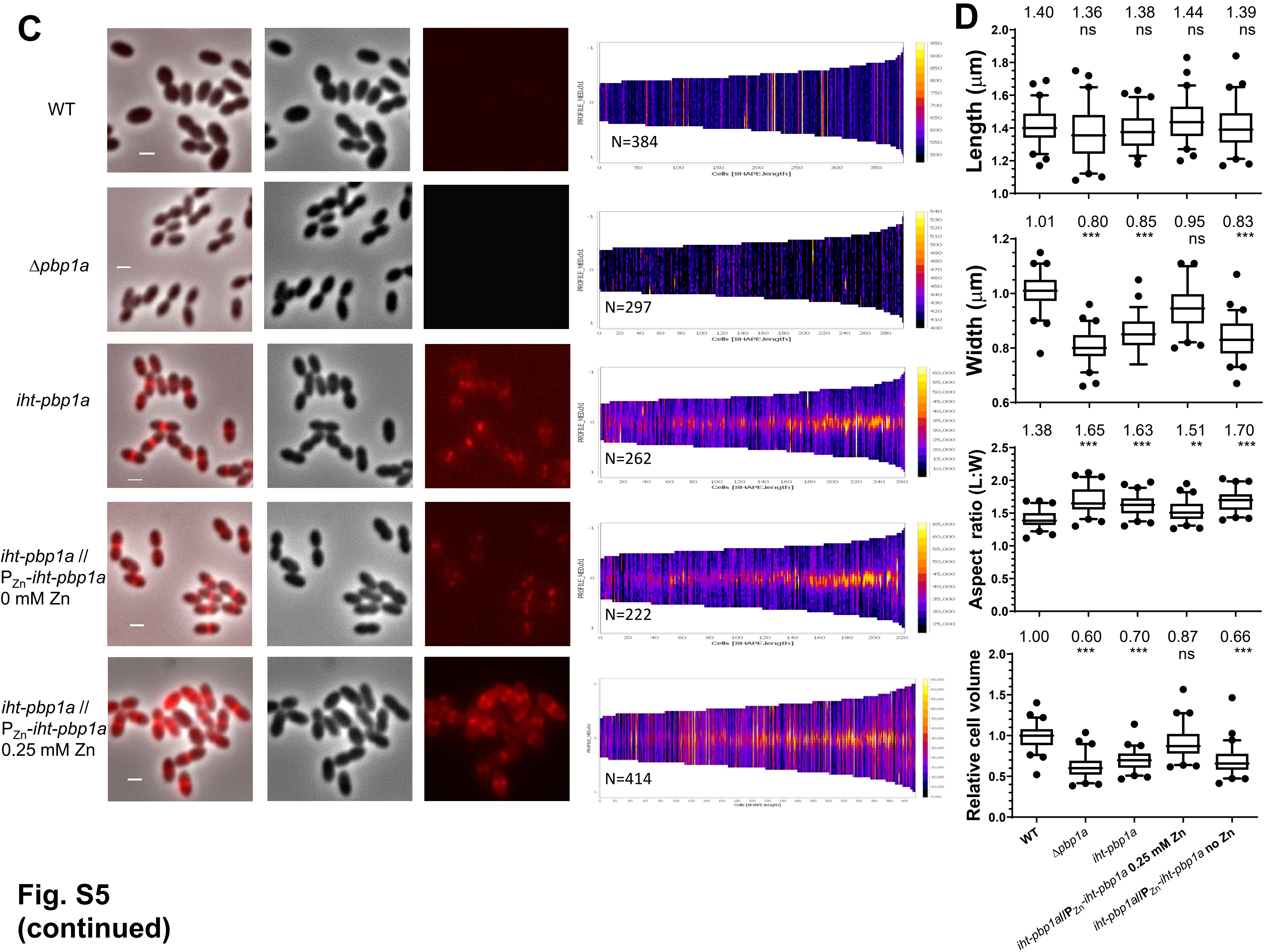


**Fig. S5 (continued)**

**Fig. S5.** The iHT-aPBP1a was expressed from the native locus at 10 ±10% (mean ± SD) of the amount of untagged WT aPBP1a. D39 Δ*cps rpsL1 pbp2b*⁺ (WT, indicated by +; IU1824); *iht-pbp1a* (iHT, IU16320); *iht-pbp1a* // $P_{Zn}$-*iht-pbp1a* (*iht-pbp1a* // $P_{Zn}$-*iht-pbp1a*, IU16497); and Δ*pbp1a* (indicated by Δ; IU6741) were grown in C+Y medium with the indicated concentrations of Zn inducer, and 2D-FM and western blot analysis were performed as described in *Materials and Methods*. (*A*) Representative growth curves of two or more biological replicates that gave similar results. Times of sample collection for microscopy and western blotting are indicated. The doubling times and growth yields (mean ± SEM) are indicated, and a one-way ANOVA with Dunnet's post-test was used to compare growth parameters among strains compared to WT. *ns* (nonsignificant); **P* < 0.05. (*B*) Western blots of cells expressing aPBP1a or iHT-aPBP1a probed with anti-aPBP1a antibody. Amounts of added Zn inducer are shown at the top; µL of sample loaded are shown at the bottom. Samples were taken at $OD_{620}$ ≈0.15 to 0.25 and resuspended in volumes of lysis buffer proportional to the final $OD_{620}$. Quantitation of relative aPBP1a amount for each lane was normalized with Total Q protein stain as described in *Materials and Methods*. Blue values in the blot show % aPBP1a expressed relative to untagged WT aPBP1a (mean ± SD), with signal from the Δ*pbp1a* lane set as zero. Red values at the top show % iHT-aPBP1a expressed relative to untagged WT aPBP1a, with background signal from the non-specific band in the Δ*pbp1a* lane subtracted. To the right of the western blot is the same blot stained with Total Q protein stain. Standard curves used to interpolate values for quantitation are shown between the blots. Best-fit lines were determined by linear regression with GraphPad Prism. A representative blot is shown from two biological replicates that gave similar results. (*C*) 2D-FM images of cells expressing aPBP1a or iHT-aPBP1a labeled with a saturating amount (500 nM) of HT-ligand. The fluorescence intensity of images for *iht-pbp1a* // $P_{Zn}$-*iht-pbp1a* +0.25 mM Zn inducer (IU16497) was lowered compared to that of the other strains to visualize localization. Scale bar = 1 µm. Demograph intensity scales (shown on the right of the demographs) were auto-adjusted based on maximal intensity reading, where N denotes total number of cells analyzed. (*D*) Box and whisker plots (whiskers, 5 and 95 percentile) of cell lengths, widths, aspect ratios (cell length to width), and relative cell volumes. n = 60 cells from two biological replicates. A Kruskal-Wallis test with Dunn's post-test was used to compare dimension parameters compared to WT values. *ns* (nonsignificant); ***P* < 0.01; ****P* < 0.001.

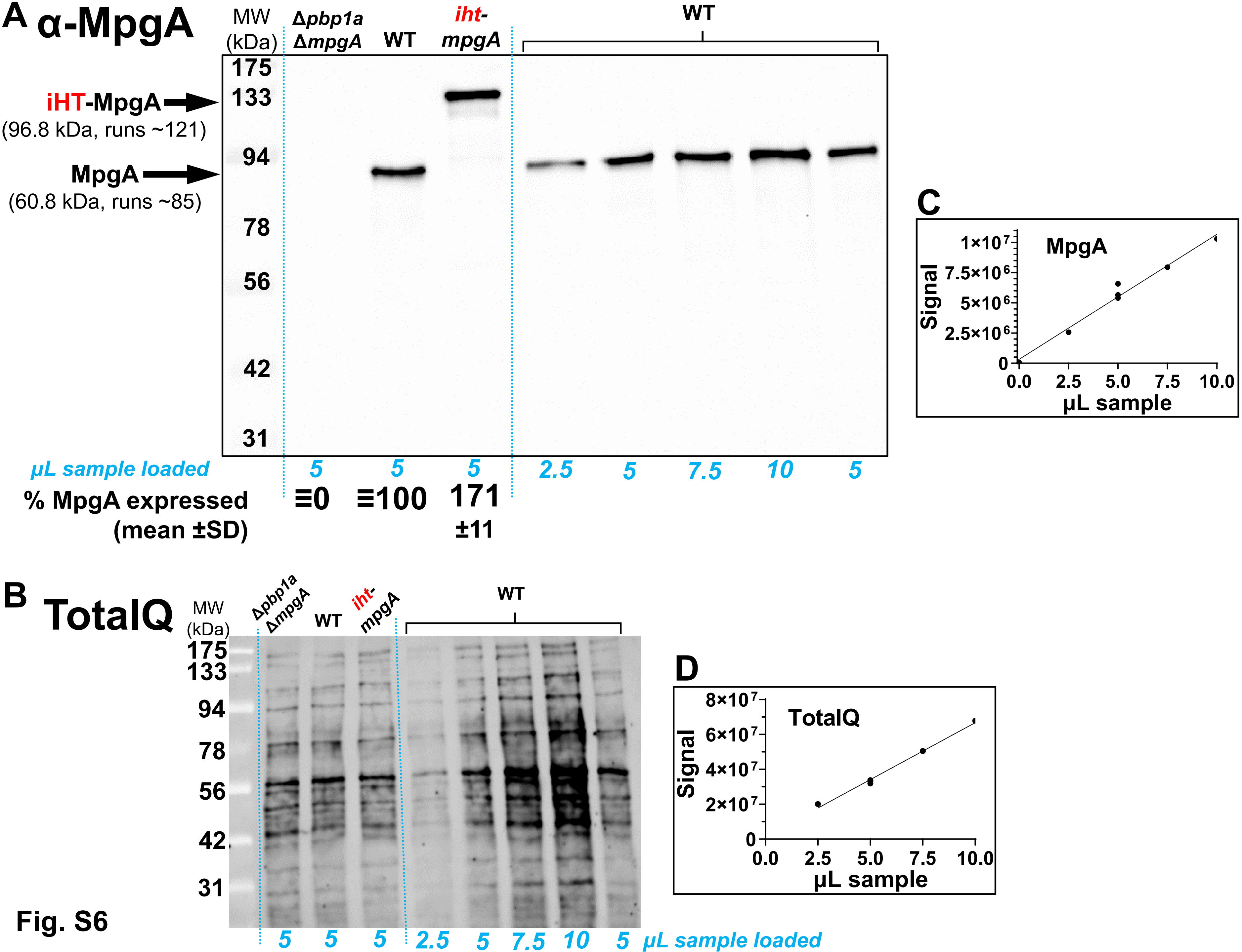
A α-MpgA
MW (kDa)
Δpbp1a ΔmpgA
WT
iht-mpgA
WT
175
133
94
78
56
42
31
iHT-MpgA
(96.8 kDa, runs ~121)
MpgA
(60.8 kDa, runs ~85)
μL sample loaded
5 5 5 2.5 5 7.5 10 5
% MpgA expressed (mean ±SD)
≡0
≡100
171 ±11
B TotalQ
MW (kDa)
Δpbp1a ΔmpgA
WT
iht-mpgA
WT
175
133
94
78
56
42
31
5 5 5 2.5 5 7.5 10 5 μL sample loaded
C
MpgA
Signal
1×10^7
7.5×10^6
5×10^6
2.5×10^6
0
0.0 2.5 5.0 7.5 10.0
μL sample
D
TotalQ
Signal
8×10^7
6×10^7
4×10^7
2×10^7
0
0.0 2.5 5.0 7.5 10.0
μL sample

Fig. S6

**Fig. S6.** iHT-MpgA was expressed from the native locus at 171 ± 11% (mean ± SD) of the amount of untagged WT MpgA. (*A*) D39 Δ*cps rpsL1 mpgA*$^+$ (WT, IU1824); *iht-mpgA* (IU15997); and Δ*pbp1a* Δ*mpgA* (IU7325) were grown in C+Y medium, and cell lysates were prepared and probed with anti-MpgA antibody in western blots as described in *Materials and Methods*. Amount of sample (µL) loaded is shown at the bottom in blue, % iHT-MpgA expressed relative to untagged WT MpgA (mean ± SD) is shown at the bottom in black. A representative blot is shown from two biological replicates that gave similar results. (*B*) Western blot shown in (*A*) stained with Total Q protein stain for normalization of total protein per lane. (*C* and *D*) Standard curves used to interpolate values of (*C*) MpgA and (*D*) Total Q signals for quantitation. Best-fit lines were determined by linear regression with GraphPad Prism.

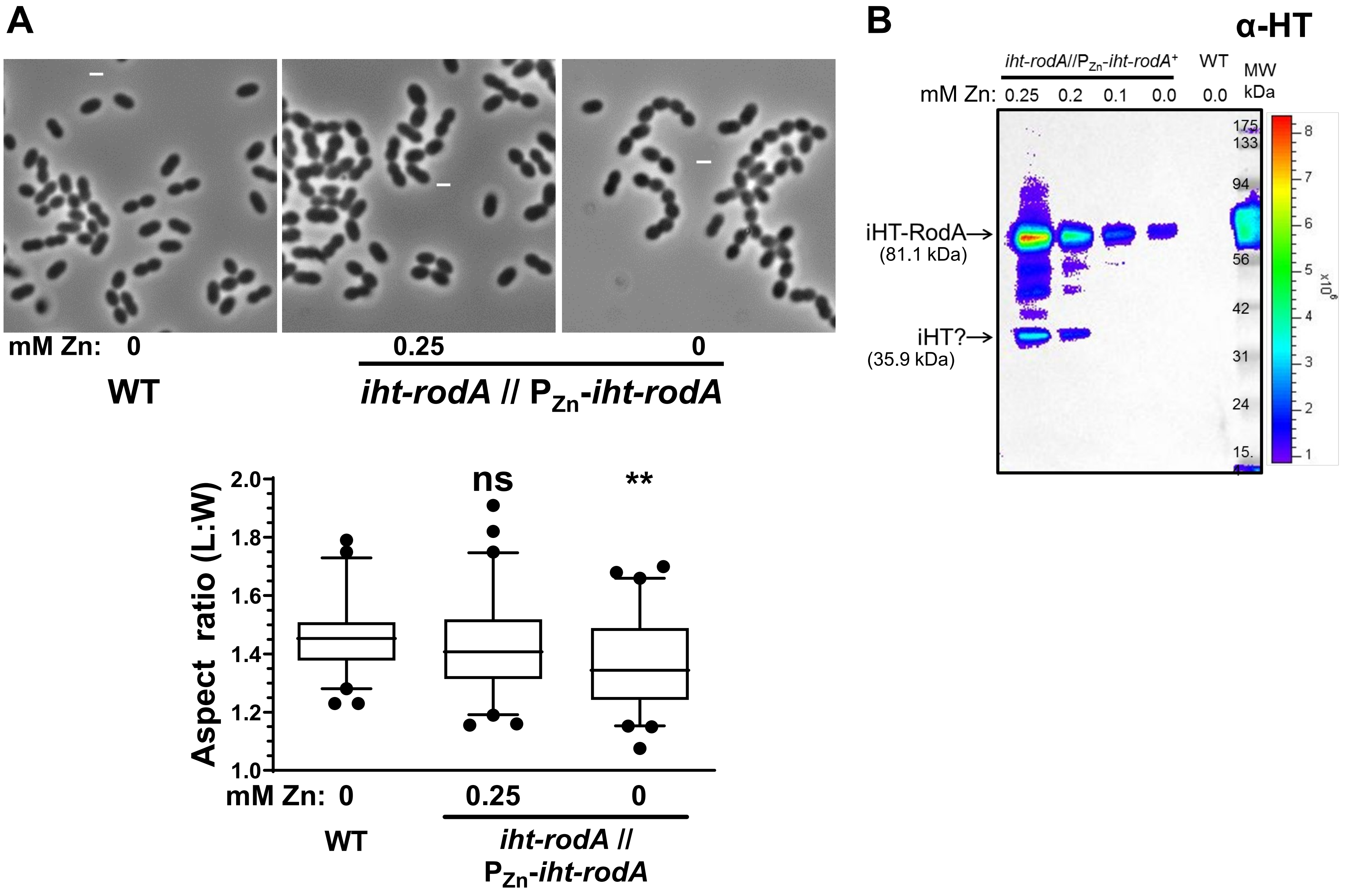
A
mM Zn: 0
0.25
0
WT
iht-rodA // PZn-iht-rodA
Aspect ratio (L:W)
2.0
1.8
1.6
1.4
1.2
1.0
ns
**
mM Zn: 0
0.25
0
WT
iht-rodA //
PZn-iht-rodA
B
α-HT
iht-rodA//PZn-iht-rodA+
WT
MW
kDa
mM Zn: 0.25 0.2 0.1 0.0 0.0
175
133
94
56
42
31
24
15.
iHT-RodA→
(81.1 kDa)
iHT?→
(35.9 kDa)
x10^6
8
7
6
5
4
3
2
1

Fig. S7

**Fig. S7.** Overexpression of iHT-RodA alleviated cell shape defects. 2D-phase contrast microscopy and western blotting were performed on WT (IU1824) and *iht-rodA* // $P_{Zn}$-*iht-rodA* (IU16496) as described in *Materials and Methods*. (*A*) Phase contrast images (upper) and box and whisker plot (whiskers, 5 and 95 percentile) of aspect ratio (lower) of IU1824 and IU16496 with or without 0.25 mM Zn inducer added. Scale bar = 1 μm. n = 60 cells from 2 biological replicates. A Kruskal-Wallis ANOVA with Dunn's post-test was used to compare aspect ratios to WT. *ns* (nonsignificant); $**P < 0.01$. (*B*) Western blot of cells expressing RodA or iHT-RodA probed with anti-HT antibody. Amount of added Zn inducer is shown at the top. A representative blot is shown from two biological replicates that gave similar results.

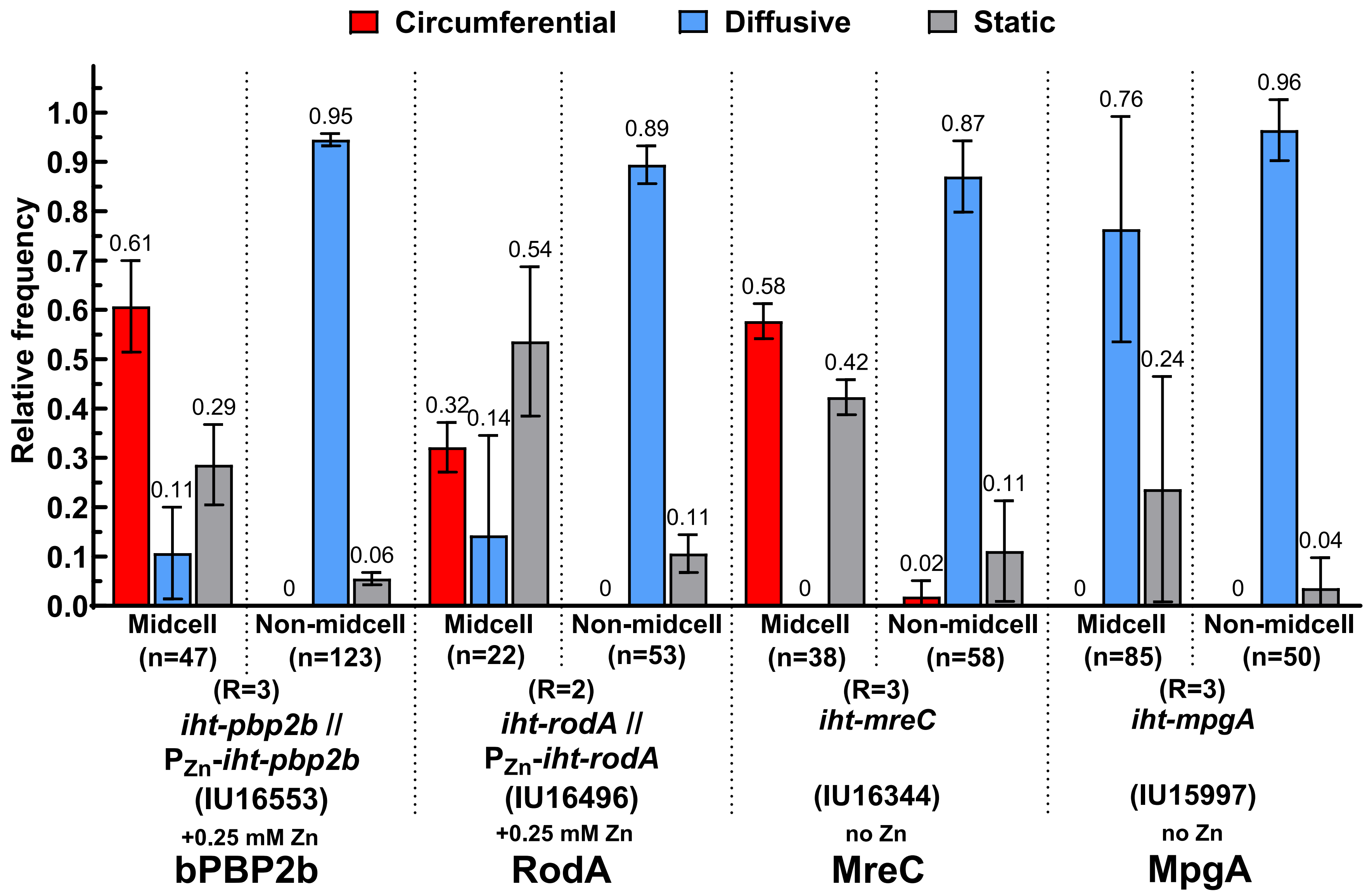


Data sorted from Figure 1*B*

**Fig. S8**

**Fig. S8.** Nearly all circumferential and static molecules of iHT-bPBP2b, iHT-RodA, and iHT-MreC, but not iHT-MpgA, were located at midcell. Data from Figure 1B were sorted into movement patterns of HT-labeled protein molecules at midcell, where PG synthesis occurred or elsewhere in cells (non-midcell). The layout is the same as Figure 1*B* (see legend for details). Strains: *iht-pbp2b* // $P_{Zn}$-*iht-pbp2b* (IU16553) + 0.25 mM Zn inducer; *iht-rodA* // $P_{Zn}$-*iht-rodA* (IU16496) + 0.25 mM Zn inducer; *iht-mreC* (IU16344); and *iht-mpgA* (IU15997). n = number of cells analyzed from R biological replicates.

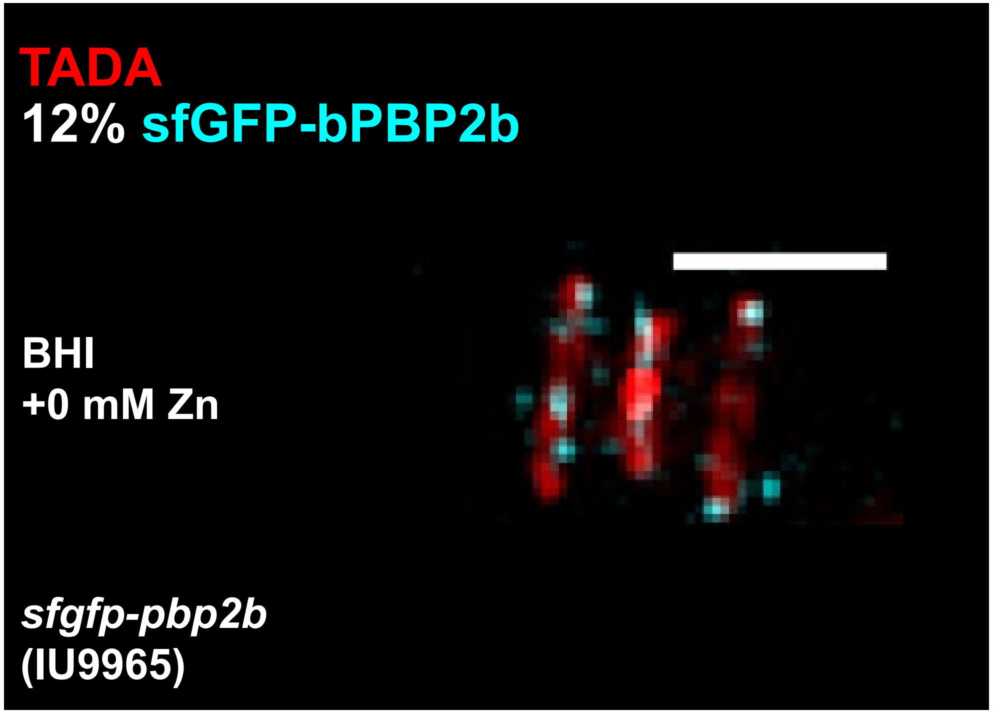


**Fig. S9.** sfGFP-bPBP2b (cyan) expressed at a low level (≈12% of WT) localized primarily to regions of active PG synthesis (red). 3D-SIM images of cells expressing sfGFP-bPBP2b (IU9965, cyan) labeled with 125 μM of the fluorescent D-amino acid TADA (red) for 2.5 m and fixed before observation as described in *Materials and Methods*. The percentage indicates the amount of sfGFP-bPBP2b expressed relative to untagged WT bPBP2b (18). Scale bar = 1 μm.

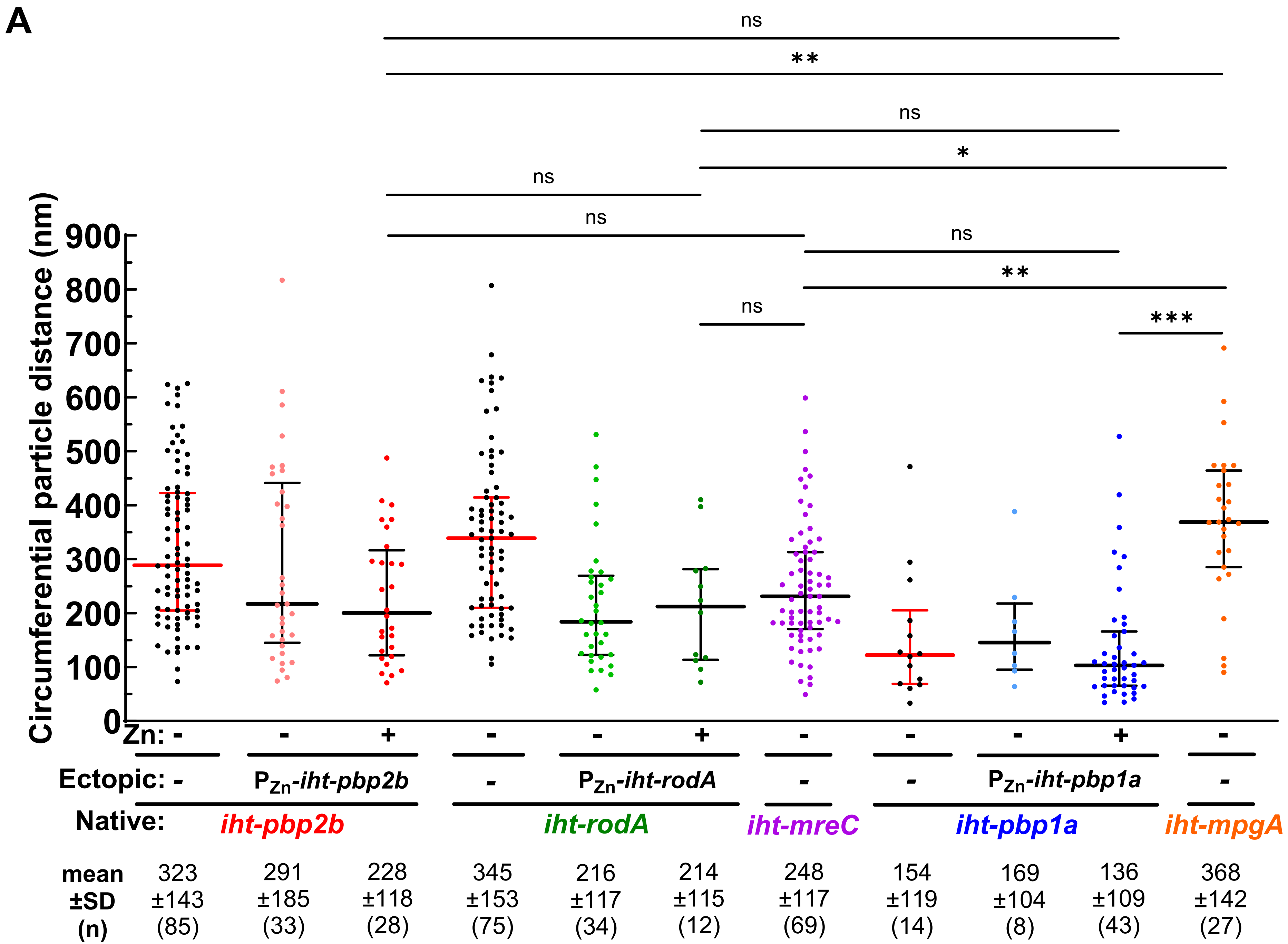


**Fig. S10** (continued on next page)

**B**

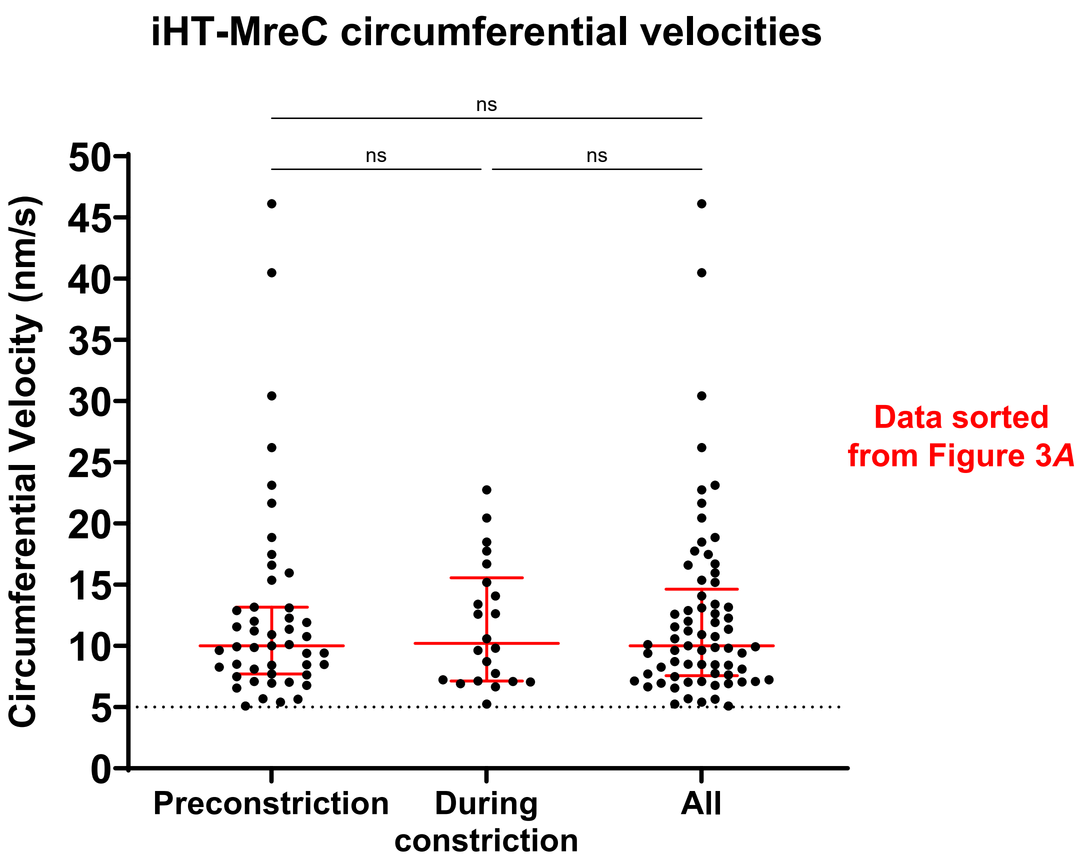


| | Preconstriction | During constriction | All |
|---|---|---|---|
| Mean | 12.8 | 11.7 | 12.5 |
| ±SD | ±8.5 | ±5.1 | ±7.5 |
| n | 47 | 22 | 69 |

**Fig. S10 (continued)**

**Fig. S10.** (*A*) Dot plot of distances traveled by circumferentially moving proteins. Sm-TIRFm at 1 FPS was performed as described in *Materials and Methods* for *iht-pbp2b* (IU15928); *iht-pbp2b* // $P_{Zn}$-*iht-pbp2b* (IU16553); *iht-rodA* (IU15970); *iht-rodA* // $P_{Zn}$-*iht-rodA* (IU16496); *iht-mreC* (IU16344); *iht-pbp1a* (IU16320); *iht-pbp1a* // $P_{Zn}$-*iht-pbp1a* (IU16497); and *iht-mpgA* (IU15997). For each circumferentially moving protein, the velocity (Figure 3*A*) was multiplied by the duration (Figure 3*B*) to determine the circumferential particle distance traveled. +Zn indicates that 0.25 mM Zn inducer was added. Black and red lines are median ± interquartile, and mean ± SD are indicated. n = total number of molecules analyzed from 2-5 biological replicates. A Kruskal-Wallis ANOVA with Dunn's post test was used to compare particle distances between strains. *ns* (nonsignificant); *$P < 0.05$; **$P < 0.01$; ***$P < 0.001$. (*B*) Circumferential velocities of iHT-MreC molecules were similar before and after cell constriction. Dot plot of circumferential velocities of iHT-MreC (IU16344) determined by sm-TIRFm at 1 FPS, replotted from Figure 3*A* and sorted based on cell constriction. Constriction was determined by visual assessment of cell outlines. Red lines are median ± interquartile, and mean ± SD are indicated. n = total molecules analyzed from 3 biological replicates. Dotted grey line indicates the minimum threshold velocity (5 nm/s). A Kruskal-Wallis ANOVA with Dunn's post test was used to compare velocities in preconstricted and constricted cells. *ns* (nonsignificant).

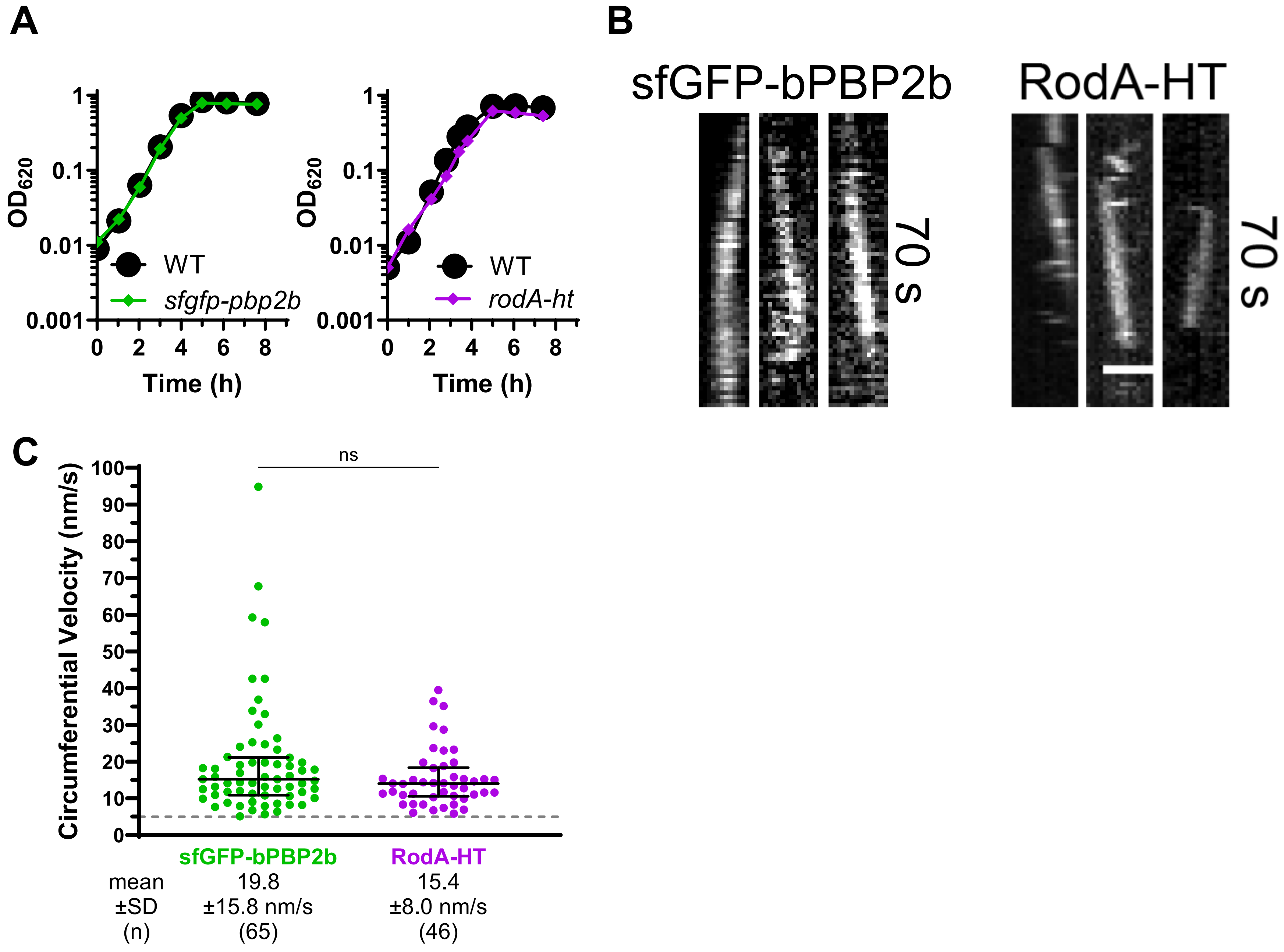
A
B
C
1
0.1
0.01
0.001
OD620
WT
sfgfp-pbp2b
rodA-ht
0 2 4 6 8
Time (h)
sfGFP-bPBP2b
RodA-HT
70 s
ns
Circumferential Velocity (nm/s)
100
90
80
70
60
50
40
30
20
10
0
sfGFP-bPBP2b
RodA-HT
mean ±SD (n)
19.8 ±15.8 nm/s (65)
15.4 ±8.0 nm/s (46)

**Fig. S11**

**Fig. S11.** sfGFP-bPBP2b and RodA-HT displayed similar circumferential movement. Strains expressing sfGFP-bPBP2b (IU9965) or RodA-HT (IU15433) were grown in C+Y medium and dynamics were determined by sm-TIRFm at 1 FPS. For RodA-HT, cells were labeled with 500 nM HT-ligand JF549. (*A*) Growth curve analysis of WT (black, IU1824); *sfgfp-pbp2b* (green, IU9965); or *rodA-ht* (purple, IU15433) grown in C+Y medium. Representative growth curves are shown from two biological replicates that gave similar results. (*B*) Representative kymographs of circumferentially moving molecules. Scale bar = 1 μm. (*C*) Dot plots of circumferentially moving molecules. Black lines are median ± interquartile, and mean ± SD are indicated for n molecules from three biological replicates for IU9965 and two replicates for IU15433. Velocities were compared using a Mann Whitney test. *ns* (nonsignificant).

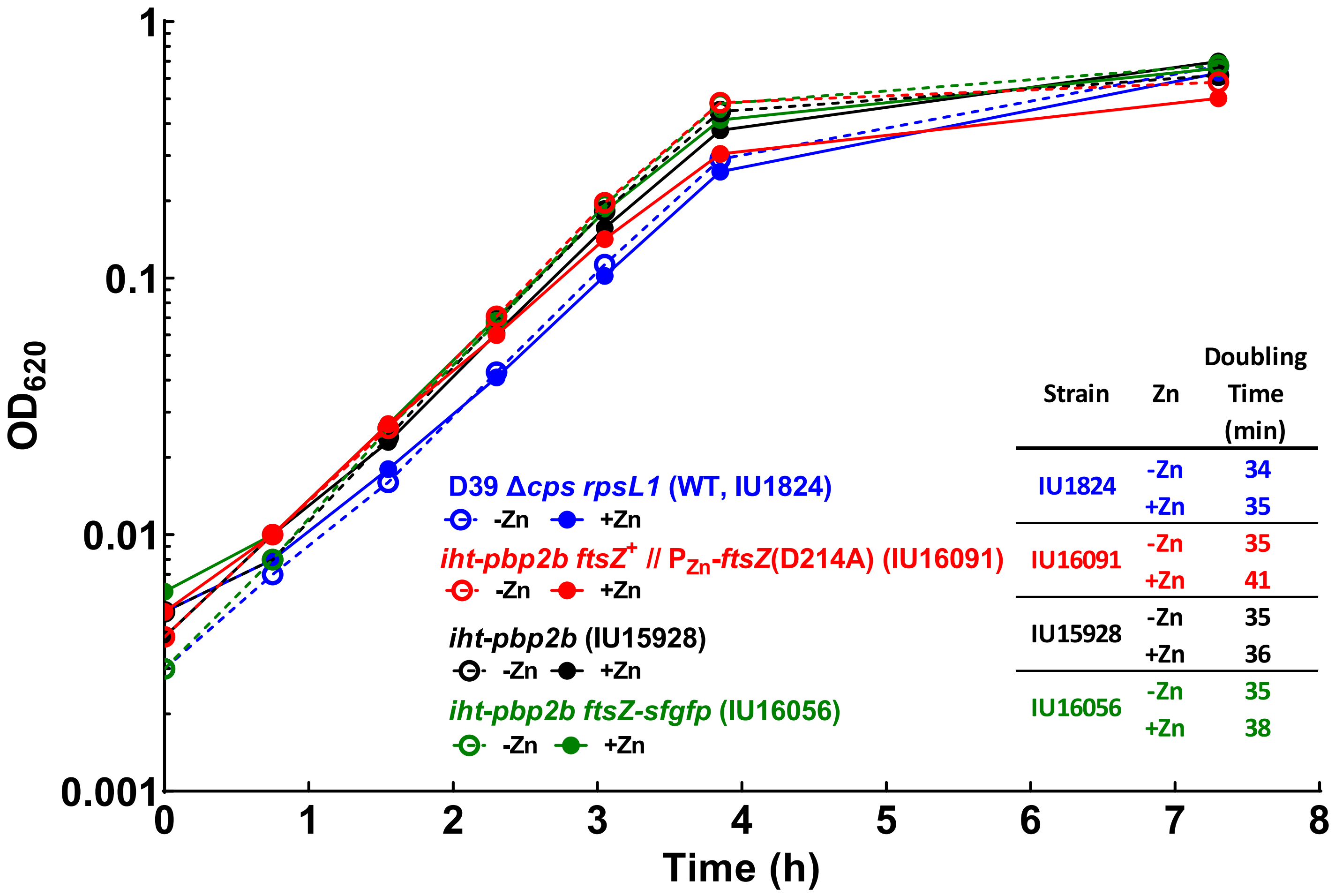


**Fig. S12.** Growth curves of WT (blue, IU1824); *iht-pbp2b ftsZ*$^+$ // $P_{Zn}$-*ftsZ*(D214A) (red, IU16091); *iht-pbp2b* (black, IU15928); and *iht-pbp2b ftsZ-sfgfp* (green, IU16056) strains in C+Y medium lacking (empty circles, dotted lines) or containing 0.25 mM (filled circles, solid lines) Zn inducer. Representative growth curves are shown from two biological replicates that gave similar results.

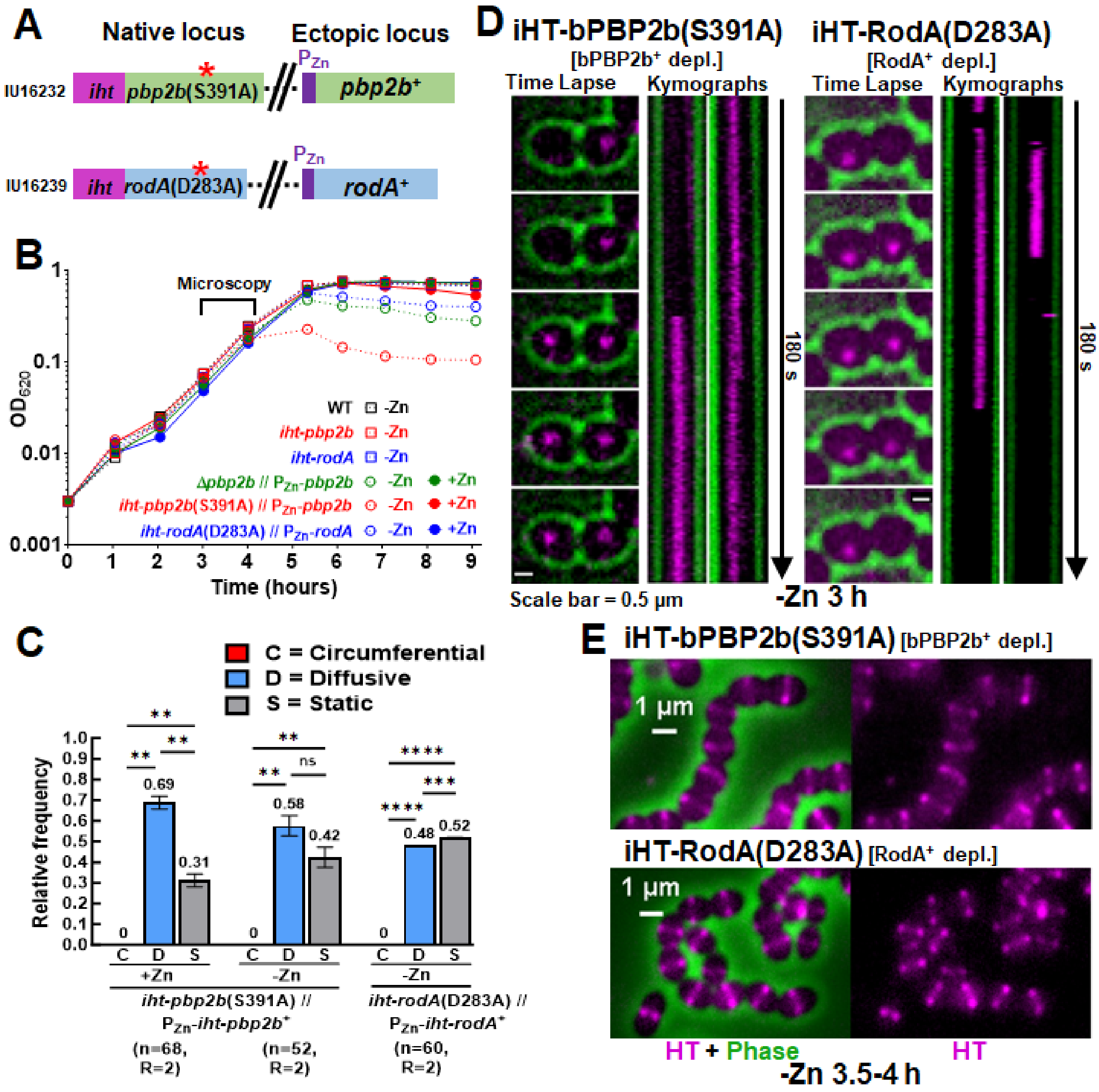
A
Native locus
Ectopic locus
IU16232
iht pbp2b(S391A)
pbp2b+
IU16239
iht rodA(D283A)
rodA+
B
Microscopy
OD620
Time (hours)
WT -Zn
iht-pbp2b -Zn
iht-rodA -Zn
Δpbp2b // PZn-pbp2b -Zn +Zn
iht-pbp2b(S391A) // PZn-pbp2b -Zn +Zn
iht-rodA(D283A) // PZn-rodA -Zn +Zn
C
C = Circumferential
D = Diffusive
S = Static
Relative frequency
+Zn
-Zn
-Zn
iht-pbp2b(S391A) // PZn-iht-pbp2b+
iht-rodA(D283A) // PZn-iht-rodA+
(n=68, R=2)
(n=52, R=2)
(n=60, R=2)
D
iHT-bPBP2b(S391A) [bPBP2b+ depl.]
iHT-RodA(D283A) [RodA+ depl.]
Time Lapse
Kymographs
180 s
Scale bar = 0.5 µm
-Zn 3 h
E
iHT-bPBP2b(S391A) [bPBP2b+ depl.]
iHT-RodA(D283A) [RodA+ depl.]
1 µm
HT + Phase
HT
-Zn 3.5-4 h

Fig. S13

**Fig. S13.** The catalytic activities of bPBP2b and RodA were required for cell viability and PG synthesis, but not localization. (*A*) Diagram of merodiploid strains containing catalytic mutant alleles fused to the *iht* protein at native loci and the Zn-inducible WT allele at the *bgaA* ectopic site. (*B*) Representative growth curves of WT (black square, IU1824), *iht-pbp2b* (red square, IU15928), *iht-rodA* (blue square, IU15970), Δ*pbp2b* // $P_{Zn}$-*pbp2b* (green, IU11258), *iht-pbp2b*(S391A) // $P_{Zn}$-*pbp2b* (red, IU16232), and *iht-rodA*(D283A) // $P_{Zn}$-*rodA* (blue, IU16239) strains grown in C+Y medium with or without 0.2 mM Zn inducer as described in the section on ectopic expression and depletion conditions in *Materials and Methods*. Two biological replicates were performed with similar results. (*C* and *D*) sm-TIRFm was performed at 1 FPS on strains IU16232 and IU16239 as described in *Materials and Methods*. (*C*) Movement patterns of iHT-bPBP2b(S391A) (IU16232) and iHT-RodA(D283A) (IU16239). The layout is the same as Figure 1*B* (see legend for details). Unpaired t-tests were performed to compare relative frequencies of motion types. *ns* (nonsignificant); ***P* < 0.01; ****P* < 0.001; *****P* < 0.0001. (*D*) Representative montages and kymographs of static molecules from strains IU16232 and IU16239 (without Zn inducer for 3 h) are displayed as described for Figure 1*A*, except the scale bar = 0.5 µm. (*E*) 2D-FM of catalytically inactive proteins. Strains IU16232 and IU16239 were grown without Zn to deplete the WT proteins (see (*B*), above) and labeled with a saturating amount (500 nM) of HT-ligand as described in *Materials and Methods*. HT-fusion proteins are shown in magenta. Scale bar = 1 µm. Two biological replicates were performed with similar results.

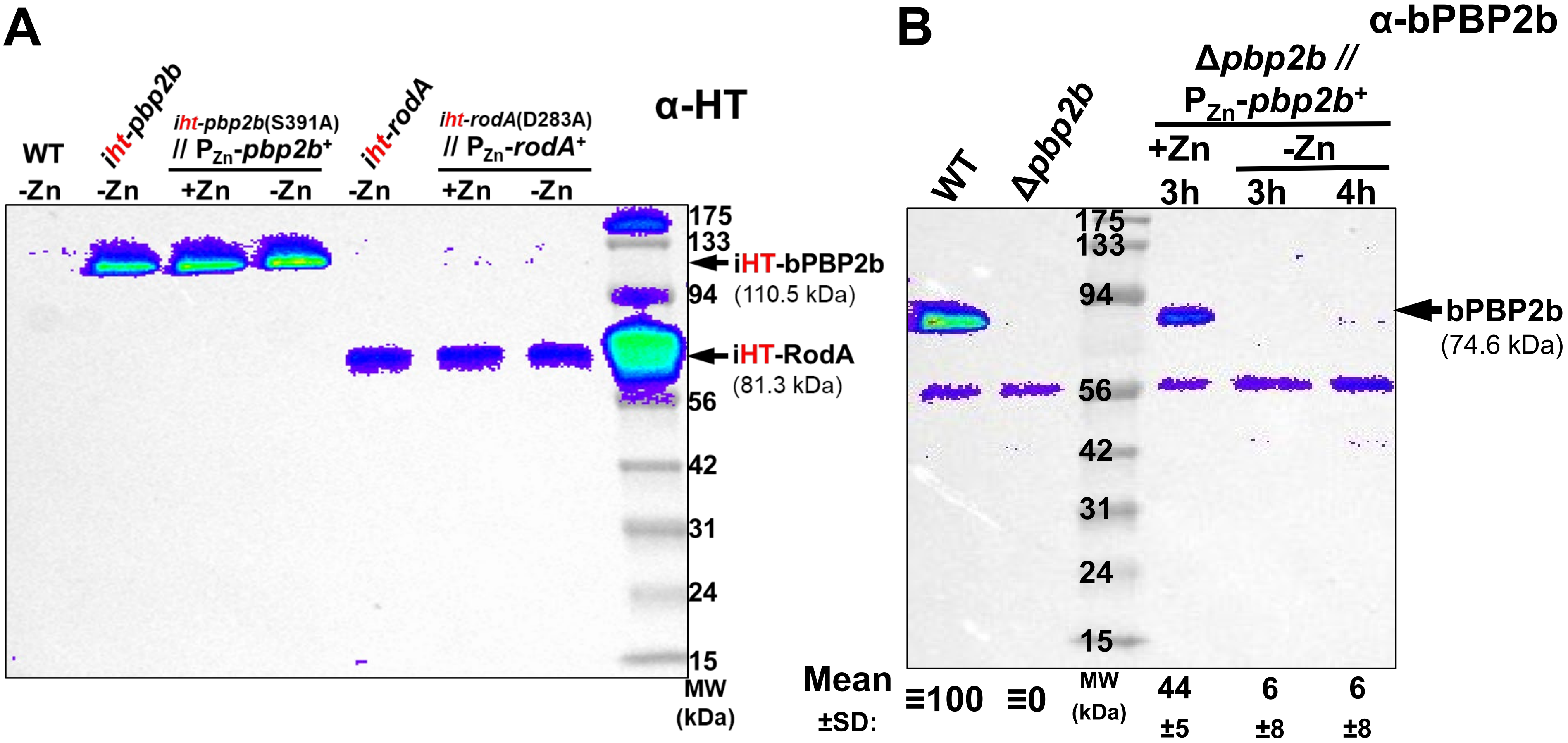


**Fig. S14.** (*A*) iHT-bPBP2b(S391A) and iHT-RodA(D283A) were expressed at comparable amounts compared to iHT-bPBP2b$^+$ and iHT-RodA$^+$. Western blot of D39 Δ*cps rpsL1* (WT, IU1824); *iht-pbp2b* (IU15928); *iht-pbp2b*(S391A) // $P_{Zn}$-*pbp2b*$^+$ (IU16232); *iht-rodA* (IU15970); and *iht-rodA*(D283A) // $P_{Zn}$-*pbp2b*$^+$ (IU16239) cells grown in C+Y medium with or without 0.2 mM Zn inducer for 3.5 h, after which cells were harvested and western blotted with anti-HT antibody as described in *Materials and Methods*. (*B*) bPBP2b is fully depleted after growth in C+Y medium lacking Zn for 3 h. Western blot of D39 Δ*cps rpsL1* (WT, IU1824); Δ*pbp2b mpgA*(Y488D) (Δ*pbp2b*, IU9783); and Δ*pbp2b* // $P_{Zn}$-*pbp2b*$^+$ (IU11258) cells probed with anti-bPBP2b antibody. 0.2 mM Zn inducer was added where indicated. Protein amounts normalized to WT (mean ± SD) are shown at the bottom. Representative blots are shown from two (*A*) or three (*B*) biological replicates that gave similar results. See the section on ectopic expression and depletion conditions in *Materials and Methods* for further details.

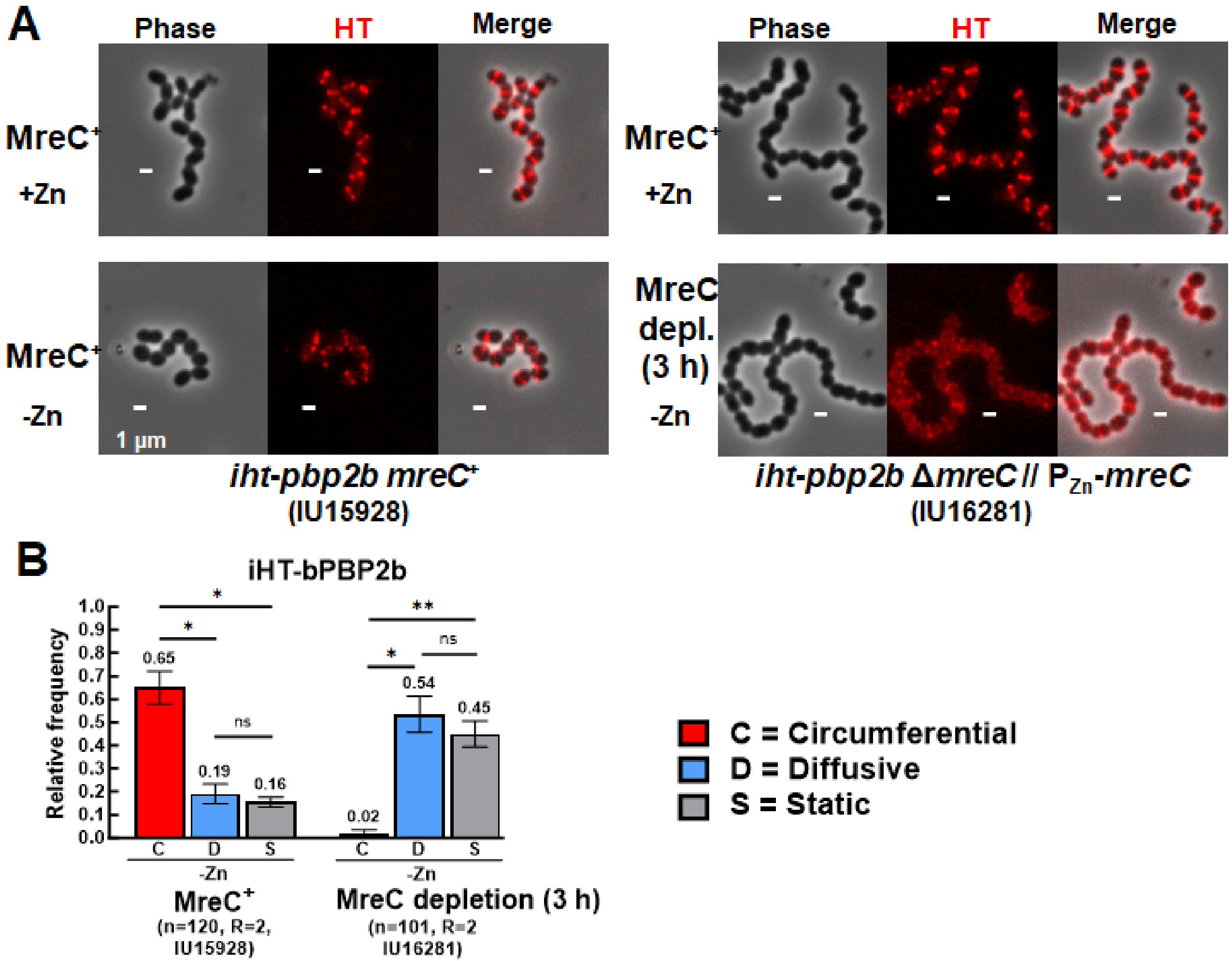


**Fig. S15.** bPBP2b localization became diffuse and circumferential movement was lost upon MreC depletion. (*A*) 2D-FM of strains *iht-pbp2b* (left, IU15928) and *iht-pbp2b ΔmreC* // $P_{Zn}$-*mreC* (right, IU16281). Cells were grown in C+Y media with or without (depletion) 0.2 mM Zn inducer, labeled with a saturating amount (500 nM) of HT-ligand, and imaged after 3 h. See the section on ectopic expression and depletion conditions in *Materials and Methods* for details. Two biological replicates were performed with similar results. (*B*) sm-TIRFm was performed at 1 FPS on strains *iht-pbp2b* (IU15928) and *iht-pbp2b ΔmreC* // $P_{Zn}$-*mreC* (IU16281) after 3 h of growth in C+Y media without Zn inducer (depletion). The layout is the same as Figure 1*B* (see legend for details). Unpaired t-tests were performed to compare relative frequencies of motion types. *ns* (nonsignificant); **P* < 0.05; ***P* < 0.01.

A

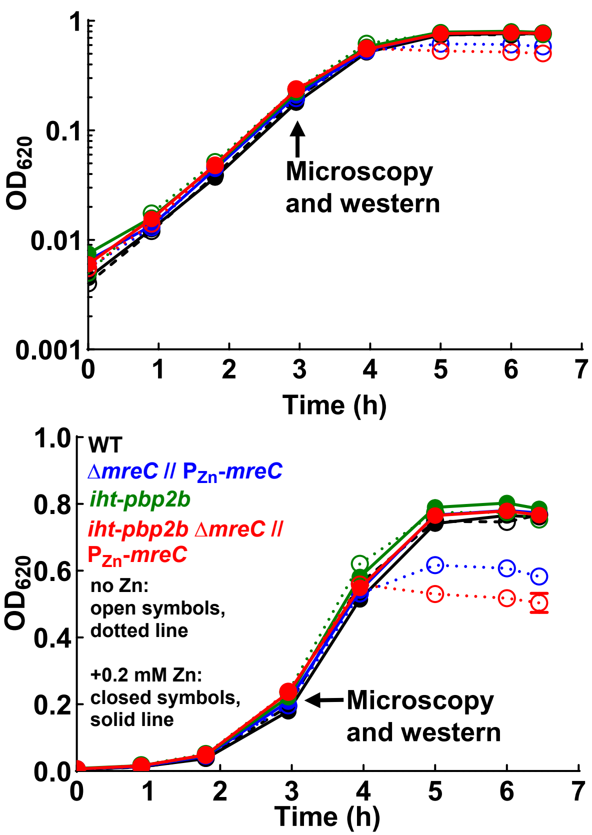


| | mM Zn | WT | *ΔmreC // $P_{Zn}$-mreC* | *iht-pbp2b* | *ΔmreC // $P_{Zn}$-mreC iht-pbp2b* |
|---|---|---|---|---|---|
| Doubling Time (min) | 0 | 30 ±1 | 33 ±1 | 32 ±1 | 29 ±2 |
| | 0.2 | 31 ±0 | 31 ±0 | 33 ±2 | 30 ±0 |
| Growth Yield ($OD_{620}$) | 0 | 0.76 ±0.02 | 0.59 ±0.04 | 0.76 ±0.02 | 0.49 ±0.09 |
| | 0.2 | 0.78 ±0.02 | 0.79 ±0.01 | 0.81 ±0.01 | 0.77 ±0.01 |

mean ± SD, R=2

B

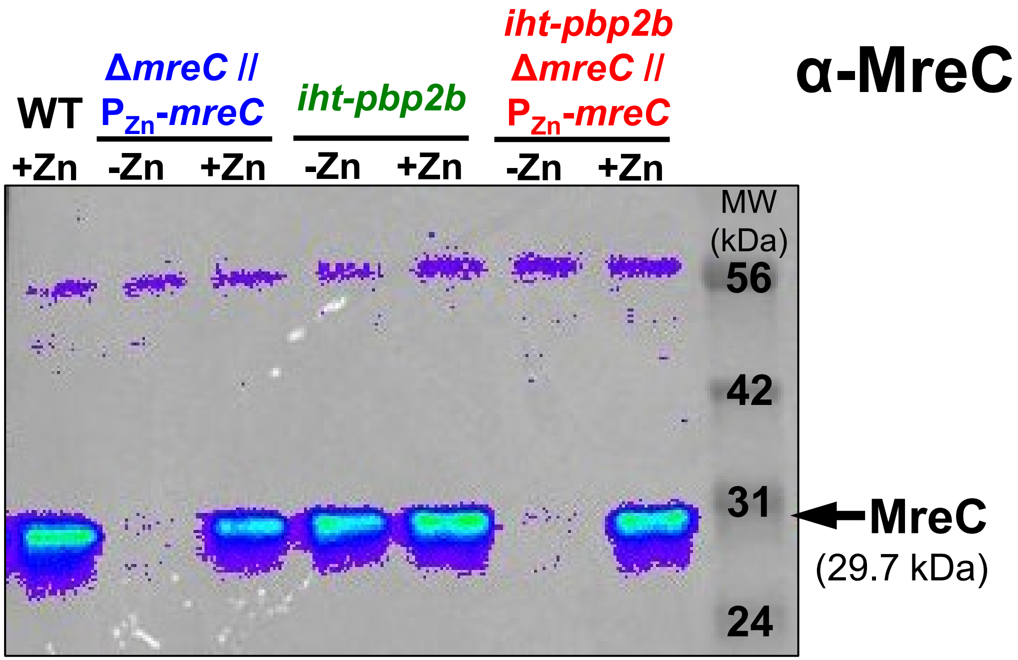


C

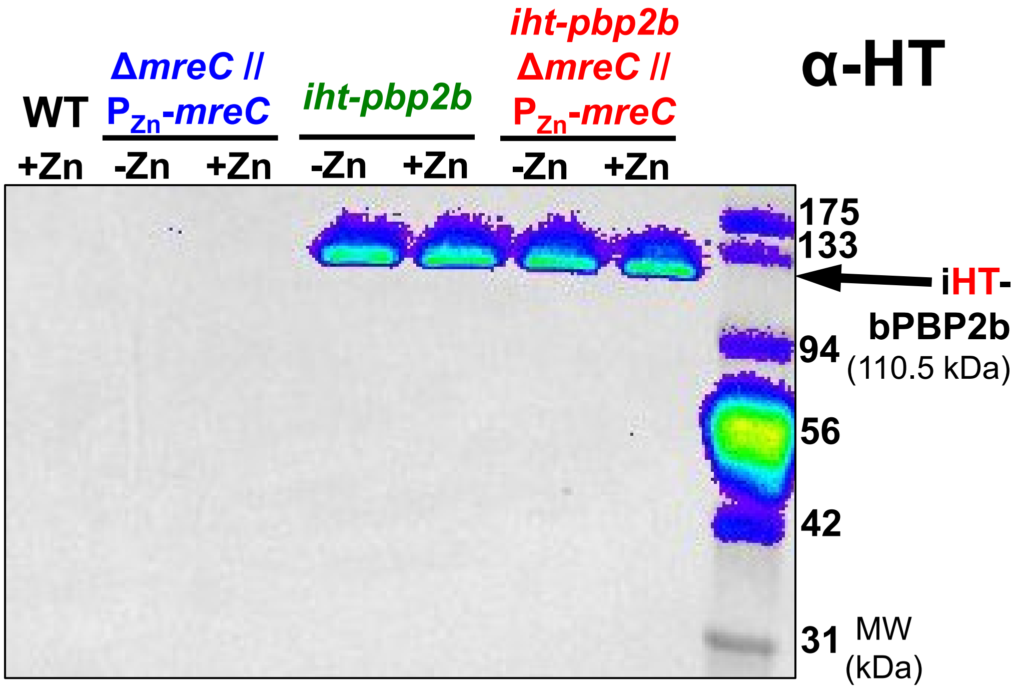


**Fig. S16**

**Fig. S16.** Depletion of MreC reduced growth yield but did not affect cellular bPBP2b amount. (*A* to *C*) Growth curves and western blots of D39 Δ*cps rpsL1* (WT, black, IU1824); Δ*mreC* // $P_{Zn}$-*mreC* (blue, IU12345); *iht-pbp2b* (green, IU15928); and *iht-pbp2b* Δ*mreC* // $P_{Zn}$-*mreC* (red, IU16281) grown in C+Y medium with or without 0.2 mM Zn inducer. See the section on ectopic expression and depletion conditions in *Materials and Methods* for additional details. (*A*) Log and linear plots of representative growth curves. Open symbols and dotted lines represent no Zn, and filled symbols and solid lines represent 0.2 mM Zn inducer. Doubling times and growth yields (mean ± SD) are shown below the graph for two biological replicates. (*B* and *C*) Representative western blots probed with anti-MreC (*B*) or anti-HT (*C*) antibodies are shown for two biological replicates that gave similar results.

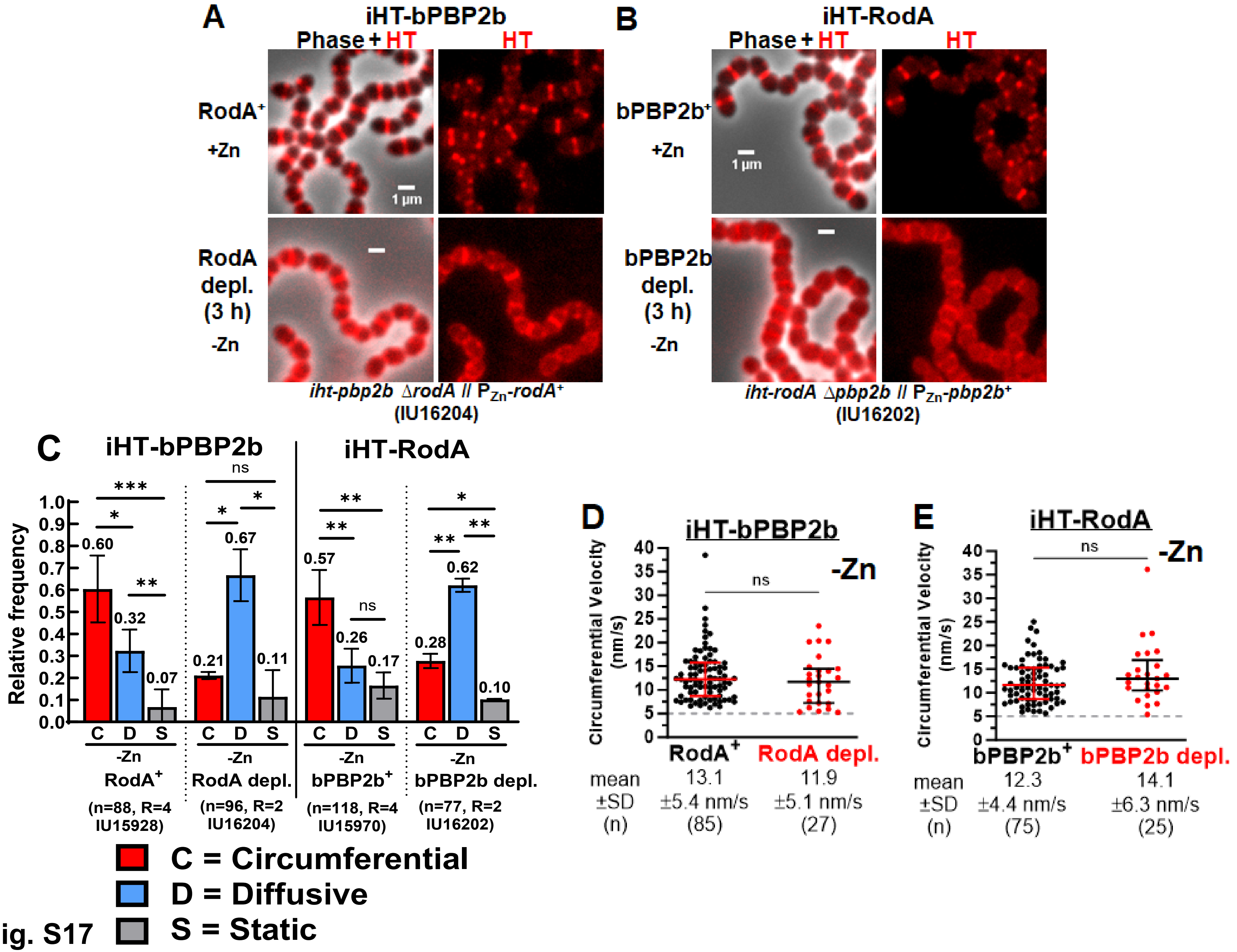
A
iHT-bPBP2b
Phase + HT
HT
RodA+
+Zn
1 µm
RodA
depl.
(3 h)
-Zn
iht-pbp2b ΔrodA // PZn-rodA+
(IU16204)
B
iHT-RodA
Phase + HT
HT
bPBP2b+
+Zn
1 µm
bPBP2b
depl.
(3 h)
-Zn
iht-rodA Δpbp2b // PZn-pbp2b+
(IU16202)
C
iHT-bPBP2b
iHT-RodA
Relative frequency
0.0 0.1 0.2 0.3 0.4 0.5 0.6 0.7 0.8 0.9 1.0
*** * ** ns * * ** ** ns * ** **
0.60 0.32 0.07
0.21 0.67 0.11
0.57 0.26 0.17
0.28 0.62 0.10
C D S
-Zn
RodA+
(n=88, R=4 IU15928)
RodA depl.
(n=96, R=2 IU16204)
bPBP2b+
(n=118, R=4 IU15970)
bPBP2b depl.
(n=77, R=2 IU16202)
C = Circumferential
D = Diffusive
S = Static
D
iHT-bPBP2b
-Zn
Circumferential Velocity (nm/s)
0 5 10 15 20 25 30 35 40
ns
RodA+
RodA depl.
mean ±SD (n)
13.1 ±5.4 nm/s (85)
11.9 ±5.1 nm/s (27)
E
iHT-RodA
-Zn
Circumferential Velocity (nm/s)
0 5 10 15 20 25 30 35 40
ns
bPBP2b+
bPBP2b depl.
mean ±SD (n)
12.3 ±4.4 nm/s (75)
14.1 ±6.3 nm/s (25)

Fig. S17

**Fig. S17.** bPBP2b and RodA required each other for localization and motion. 2D-FM and sm-TIRFm at 1 FPS were performed on strains *iht-pbp2b* Δ*rodA* // $P_{Zn}$-*rodA* (IU16204) and *iht-rodA* Δ*pbp2b* // $P_{Zn}$-*pbp2b* (IU16202). (*A* and *B*) 2D-FM showing localization in cells of strains (*A*) iHT-bPBP2b (IU16204) and (*B*) iHT-RodA (IU16202) grown with or without (depletion; depl.) 0.2 mM Zn inducer for 3 h and labeled with a saturating amount (500 nM) of HT-ligand. See the section on ectopic expression and depletion conditions in *Materials and Methods* for additional details. Two biological replicates were performed with similar results. (*C* to *E*) sm-TIRFm was performed at 1 FPS on IU16204 and IU16202 without Zn inducer (depletion) for 3 hours as described in *Materials and Methods*. (*C*) Movement patterns of HT-labeled molecules. Results for the *iht-pbp2b* Δ*rodA* // $P_{Zn}$-*rodA* (IU16204) and *iht-rodA* Δ*pbp2b* // $P_{Zn}$-*pbp2b* (IU16202) strains were determined and are graphed with data for *iht-pbp2b* (IU15928) and *iht-rodA* (IU15970) replotted from Figure 1*B* for comparison. The layout is the same as Figure 1*B* (see legend for details). Unpaired t-tests were performed to compare relative frequencies of motion types. *ns* (nonsignificant), **P* < 0.05; ***P* < 0.01; ****P* < 0.001. (*D* and *E*) Dot plots of circumferential velocities with mean values ± SD indicated for (*D*) strains IU15928 (black) and IU16204 (red), and (*E*) strains IU15970 (black) and IU16202 (red). Data from stains IU15928 and IU15970 are replotted from Figure 3*A* for comparison. Black and red lines are median ± interquartile, and mean ± SD are indicated. n = total molecules analyzed from two biological replicates. A Mann-Whitney test was used to compare velocities upon protein depletion (depl.). *ns* (nonsignificant).

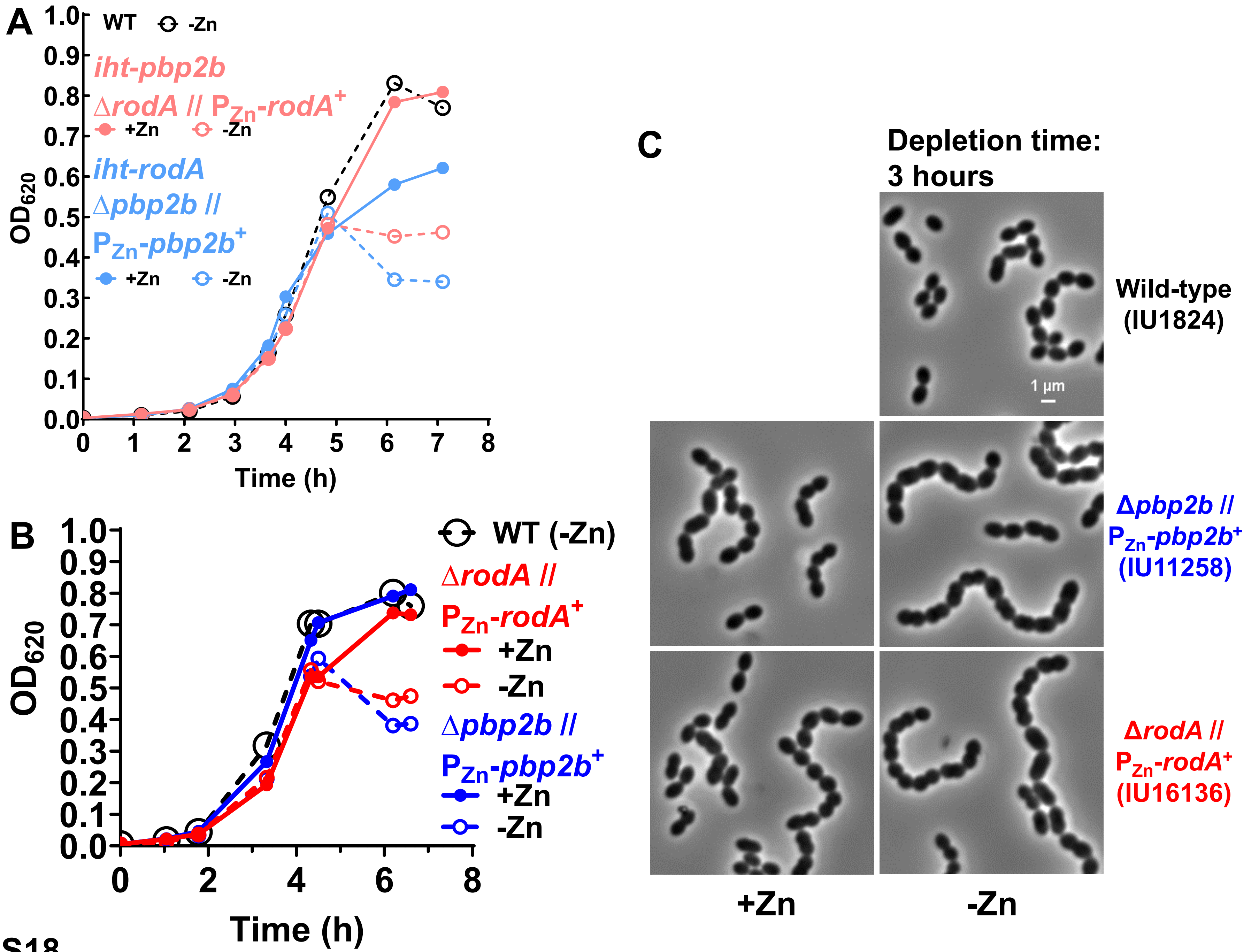
A
WT -Zn
iht-pbp2b
ΔrodA // PZn-rodA+
+Zn -Zn
iht-rodA
Δpbp2b //
PZn-pbp2b+
+Zn -Zn
OD620
Time (h)
B
WT (-Zn)
ΔrodA //
PZn-rodA+
+Zn
-Zn
Δpbp2b //
PZn-pbp2b+
+Zn
-Zn
OD620
Time (h)
C
Depletion time:
3 hours
Wild-type
(IU1824)
1 µm
Δpbp2b //
PZn-pbp2b+
(IU11258)
ΔrodA //
PZn-rodA+
(IU16136)
+Zn
-Zn

Fig. S18

**Fig. S18.** Depletion of RodA or bPBP2b reduced growth yield and led to rounded, short cells. Growth curves of (*A*) D39 Δ*cps rpsL1* (WT, black, IU1824); *iht-pbp2b* Δ*rodA* // $P_{Zn}$-*rodA*$^+$ (pink, IU16204); *iht-rodA* Δ*pbp2b* // $P_{Zn}$-*pbp2b*$^+$ (light blue, IU16202); and (*B*) D39 Δ*cps rpsL1* (WT, black, IU1824); Δ*rodA* // $P_{Zn}$-*rodA*$^+$ (red, IU16136); and Δ*pbp2b* // $P_{Zn}$-*pbp2b*$^+$ (blue, IU11258) grown in C+Y medium lacking or containing 0.2 mM Zn inducer. Open symbols and dotted lines represent no Zn, and filled symbols and solid lines represent 0.2 mM Zn inducer. (*C*) 2D phase contrast microscopy of strains from (*B*) 3 h after removal of Zn inducer to cause depletion. Scale bar = 1 μm. See the section on ectopic expression and depletion conditions in *Materials and Methods* for additional details. Representative growth curves and images are shown from three biological replicates that gave similar results.

A

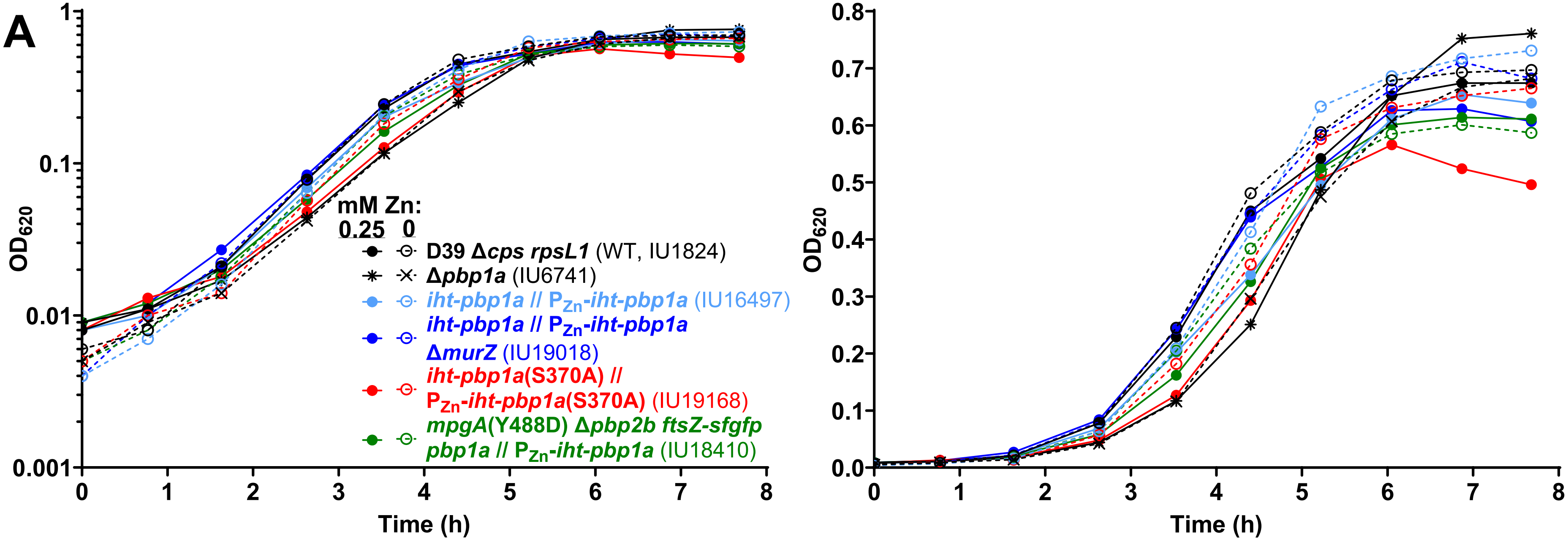


| | mM Zn | WT | *Δpbp1a* | *iht-pbp1a* // $P_{Zn}$-*iht-pbp1a* | *iht-pbp1a* // $P_{Zn}$-*iht-pbp1a* *ΔmurZ* | *iht-pbp1a* (S370A) // $P_{Zn}$-*iht-pbp1a* (S370A) | *mpgA* (Y488D) *Δpbp2b* ftsZ-sfgfp *pbp1a* // $P_{Zn}$-*iht-pbp1a* |
|---|---|---|---|---|---|---|---|
| Doubling Time (m) | 0 | 31 ±2 | 35 ±2 | 30 ±2 | 33 ±0 | 32 ±1 | 32 ±0 |
| | 0.25 | 34 ±0 | 40 ±1 | 35 ±0 | 36 ±0 | 41 ±3 | 37 ±0 |
| Growth Yield | 0 | 0.65 ±0.06 | 0.67 ±0.02 | 0.71 ±0.02 | 0.71 ±0.01 | 0.67 ±0.01 | 0.62 ±0.03 |
| | 0.25 | 0.65 ±0.03 | 0.72 ±0.06 | 0.63 ±0.03 | 0.64 ±0.01 | 0.56 ±0.01 | 0.63 ±0.02 |

mean ± SD, R=2

**Fig. S19**

**(continued on next page)**

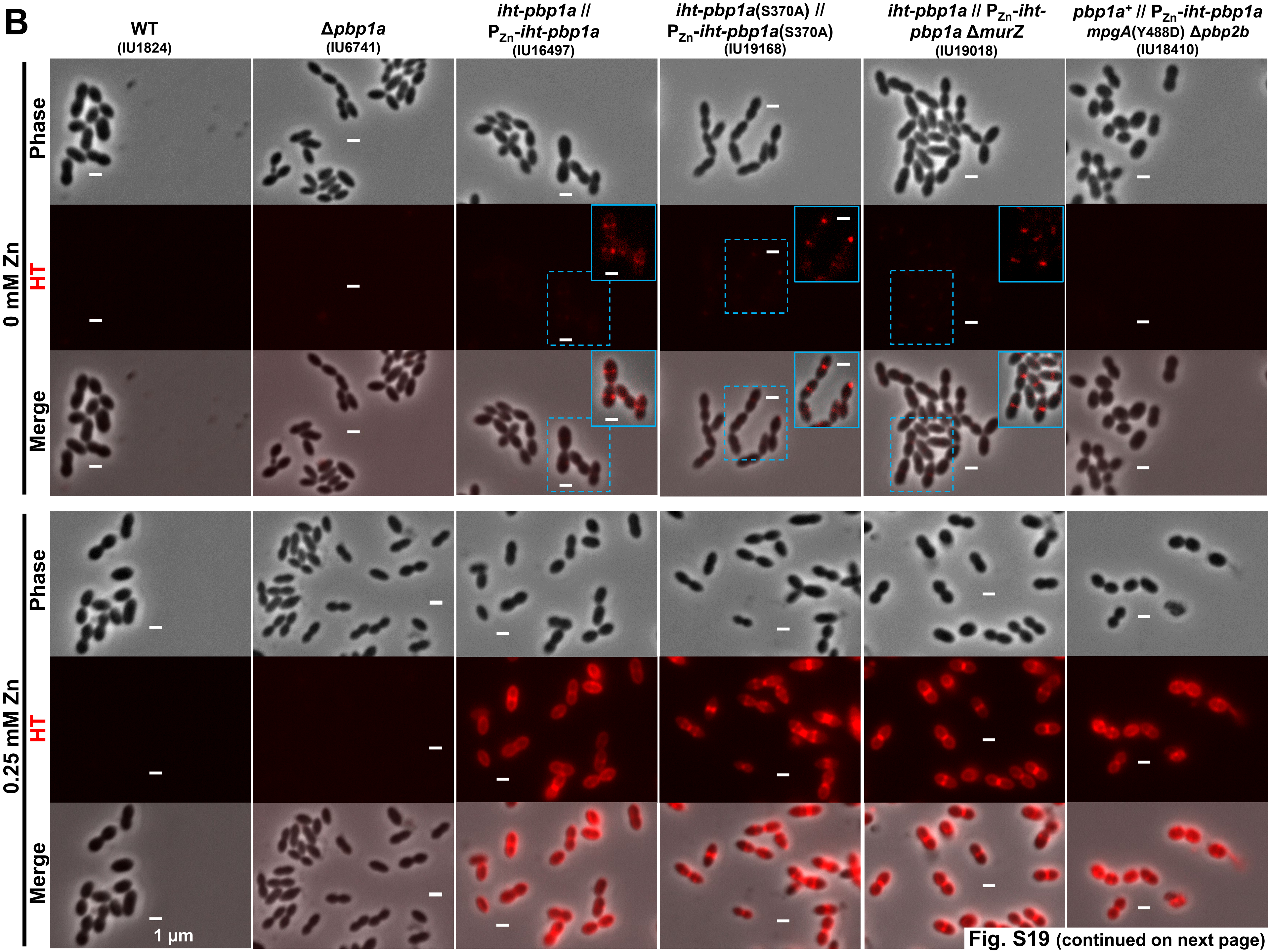


Fig. S19 (continued on next page)

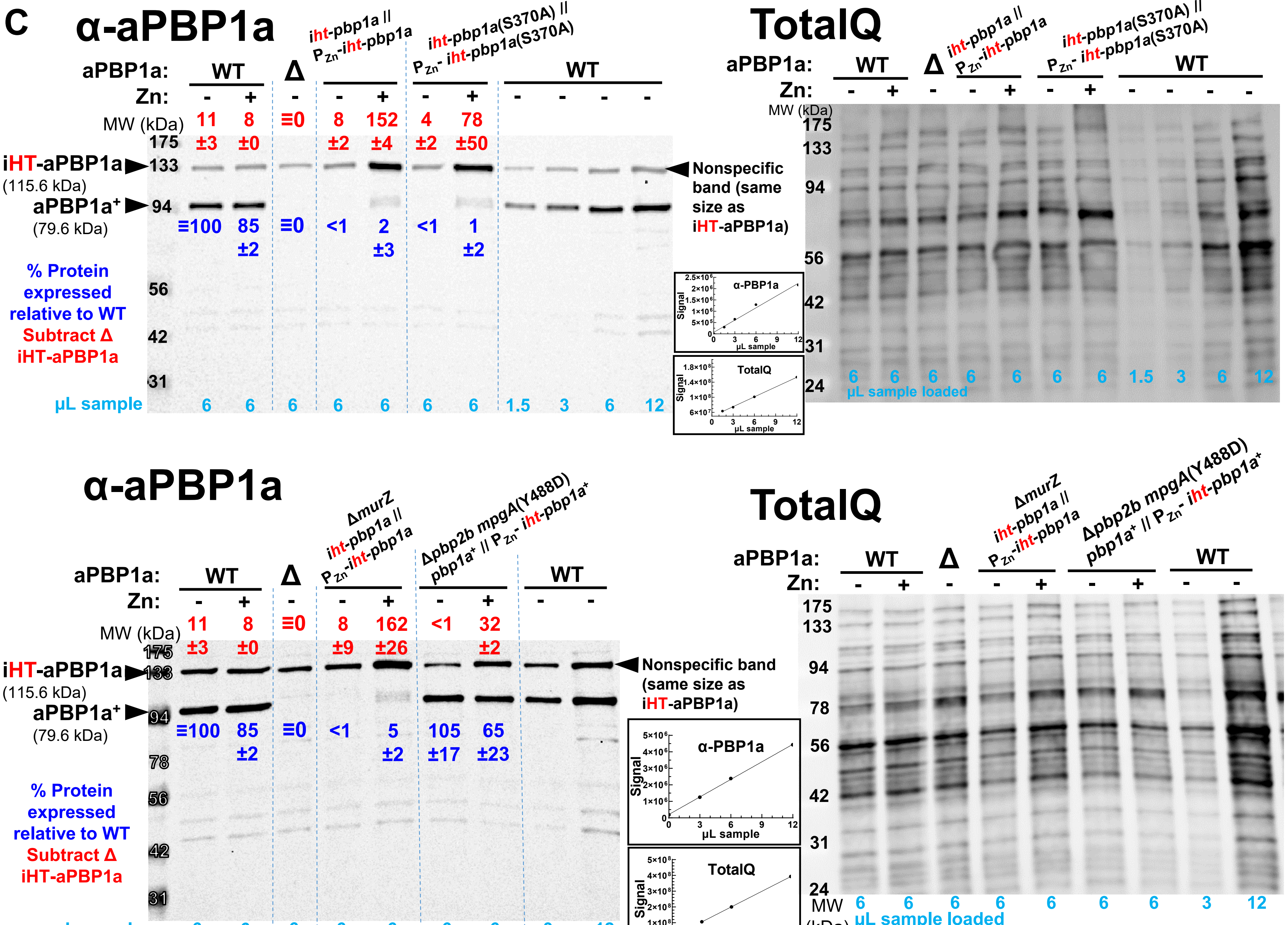


Fig. S19 (continued)

**Fig. S19.** Growth, localization, and expression of iHT-aPBP1a variants in *pbp1a* mutant strains. (*A*) Log and linear plots of representative growth curves in C+Y media ± 0.25 mM Zn inducer of strains D39 Δ*cps rpsL1* (WT, IU1824, black circles); Δ*pbp1a* (IU6741, black ex and asterisk); *iht-pbp1a* // $P_{Zn}$-*iht-pbp1a* (IU16497, light blue); *iht-pbp1a* // $P_{Zn}$-*iht-pbp1a* Δ*murZ* (IU19018, dark blue); *iht-pbp1a*(S370A) // $P_{Zn}$-*iht-pbp1a*(S370A) (IU19168, red); and *mpgA*(Y488D) Δ*pbp2b ftsZ-sfgfp pbp1a* // $P_{Zn}$-*iht-pbp1a* (IU18410, green). Open symbols with dotted lines represent no Zn, and filled symbols with solid lines represent addition of 0.25 mM Zn inducer. Doubling times (m) and growth yields ($OD_{620}$) (mean ± SD) from two biological replicates are compiled below the graphs. (*B*) Phase contrast and fluorescence microscopy images of strains described in (*A*) grown in C+Y ± 0.25 mM Zn inducer for 3.5 h, after which cells were labeled with a saturating amount (500 nM) of HT-TMR ligand. Due to low iHT-aPBP1a amounts in cells without Zn, the HT signal was increased to visualize iHT-aPBP1a localization, shown in light blue insets. The portion of the field shown in the inset is denoted by the dotted blue box. (*C*) Western blot of strains described in (A) grown in C+Y ± 0.25 mM Zn inducer for 3.5 h and blotted against α-PBP1a antibody. Dark blue values show % aPBP1a expressed relative to untagged WT (mean ± SD), with the signal from Δ*pbp1a* set as zero. Red values show % iHT-aPBP1a expressed relative to untagged WT, with the background signal from the non-specific band of Δ*pbp1a* subtracted. Volumes of sample loaded are shown along the bottom of the blot in light blue. The same blots stained with Total Q protein stain are shown at the right. Standard curves used to interpolate values for quantitation are shown between the blots. Best-fit lines were determined by linear regression with GraphPad Prism. Representative growth curves, microscopy images, and western blots are shown from least two biological replicates that gave similar results.

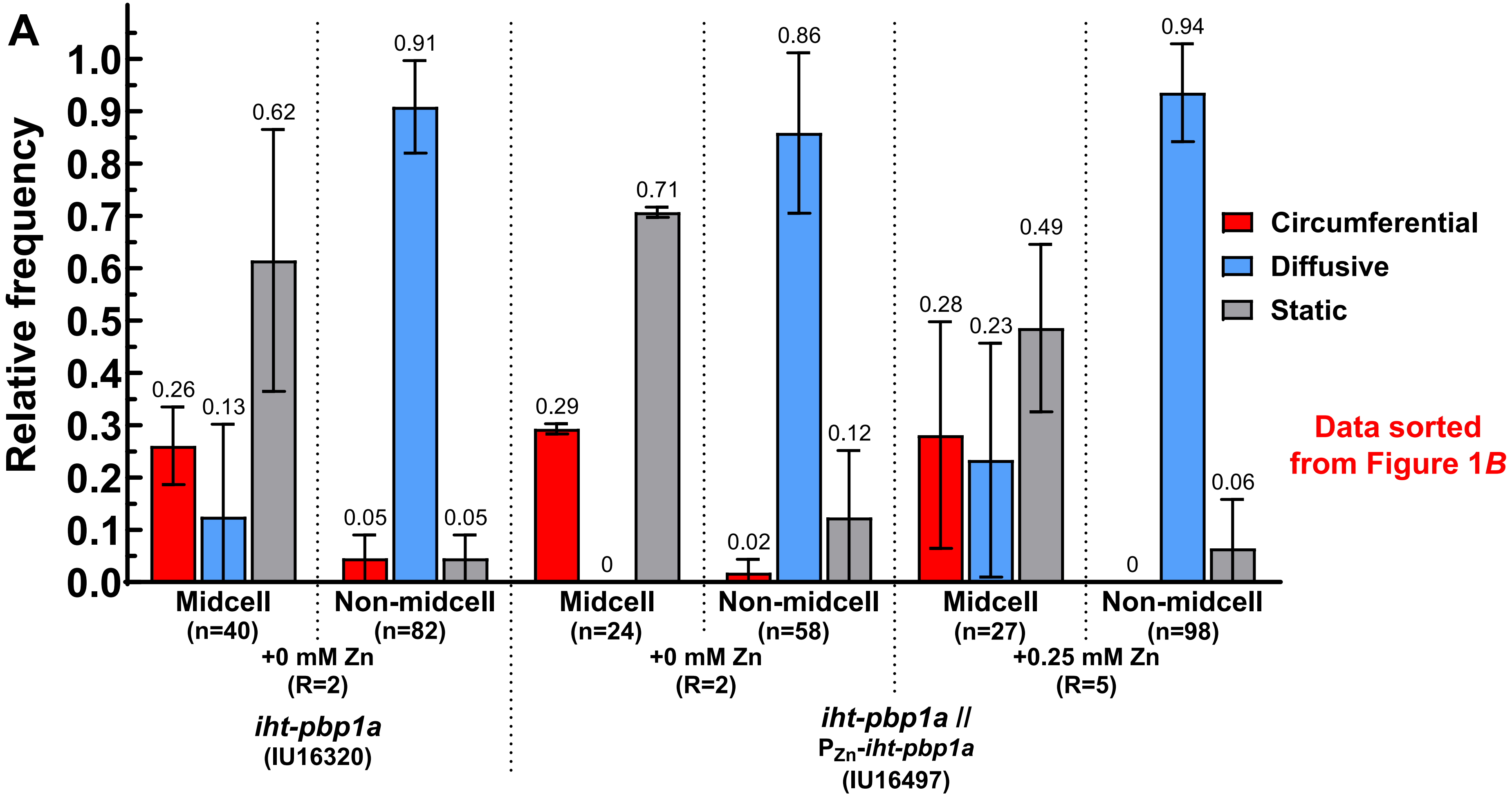


**Fig. S20**
**(continued on next page)**

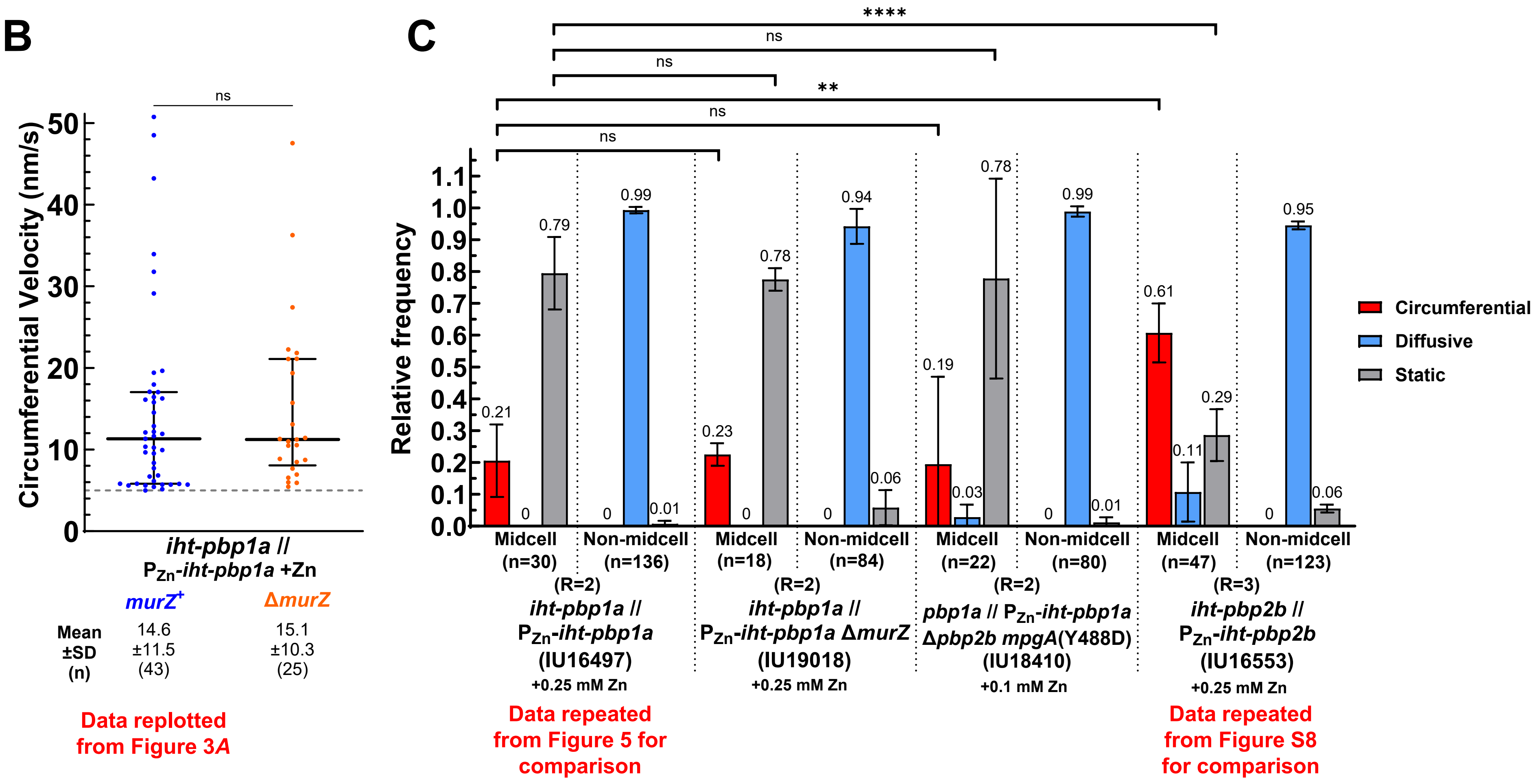


**Fig. S20 (continued)**

**Fig. S20.** aPBP1a dynamics did not change when PG precursors were limited, and aPBP1a did not complement the function of bPBP2b. Sm-TIRFm was performed at 1 FPS as described in *Materials and Methods*. (*A* and *C*) Relative frequency of movement patterns of iHT-aPBP1a or iHT-bPBP2b in midcell regions, where PG synthesis occurred, or elsewhere in cells (non-midcell). The layout is the same as Figure 1*B* (see legend for details). (*A*) iHT-aPBP1a data from Figure 1*B* were re-graphed, and sorted manually into molecules in the midcell region or in non-midcell regions. Strain *iht-pbp1a* (IU16320) was grown in C+Y medium without Zn, and *iht-pbp1a* // $P_{Zn}$-*iht-pbp1a* (IU16497) was grown in C+Y medium without or with 0.25 mM Zn. n is the number of molecules from R biological replicates. (*B*) Dot plots of circumferential velocities of *iht-pbp1a* // $P_{Zn}$-*iht-pbp1a* (IU16497) re-graphed from Figure 3*A* for comparison and *iht-pbp1a* // $P_{Zn}$-*iht-pbp1a* Δ*murZ* (IU19018) + 0.25 mM Zn inducer. Black lines are median ± interquartile, and mean ± SD are indicated. n = total molecules analyzed from 2 (IU19018) or 5 (IU16497) biological replicates. Dotted grey line indicates the minimum threshold velocity (5 nm/s). Mann-Whitney tests were done to compare the velocities between the strains. *ns* (nonsignificant). (*C*) *iht-pbp1a* // $P_{Zn}$-*iht-pbp1a* (IU16497) + 0.25 Zn mM inducer re-graphed from Figure 5 for comparison; *iht-pbp1a* // $P_{Zn}$-*iht-pbp1a* Δ*murZ* (IU19018) + 0.25 mM Zn inducer; *mpgA*(Y488D) Δ*pbp2b ftsZ-sfgfp pbp1a* // $P_{Zn}$-*iht-pbp1a* (IU18410) + 0.1 mM Zn inducer; and *iht-pbp2b* // $P_{Zn}$-*iht-pbp2b* (IU16553) + 0.25 mM Zn inducer re-graphed from Figure 1B for comparison. Mann-Whitney tests were done to compare the relative frequencies of motion type between the strains. *ns* (nonsignificant); ***P* < 0.01; *****P* < 0.0001.

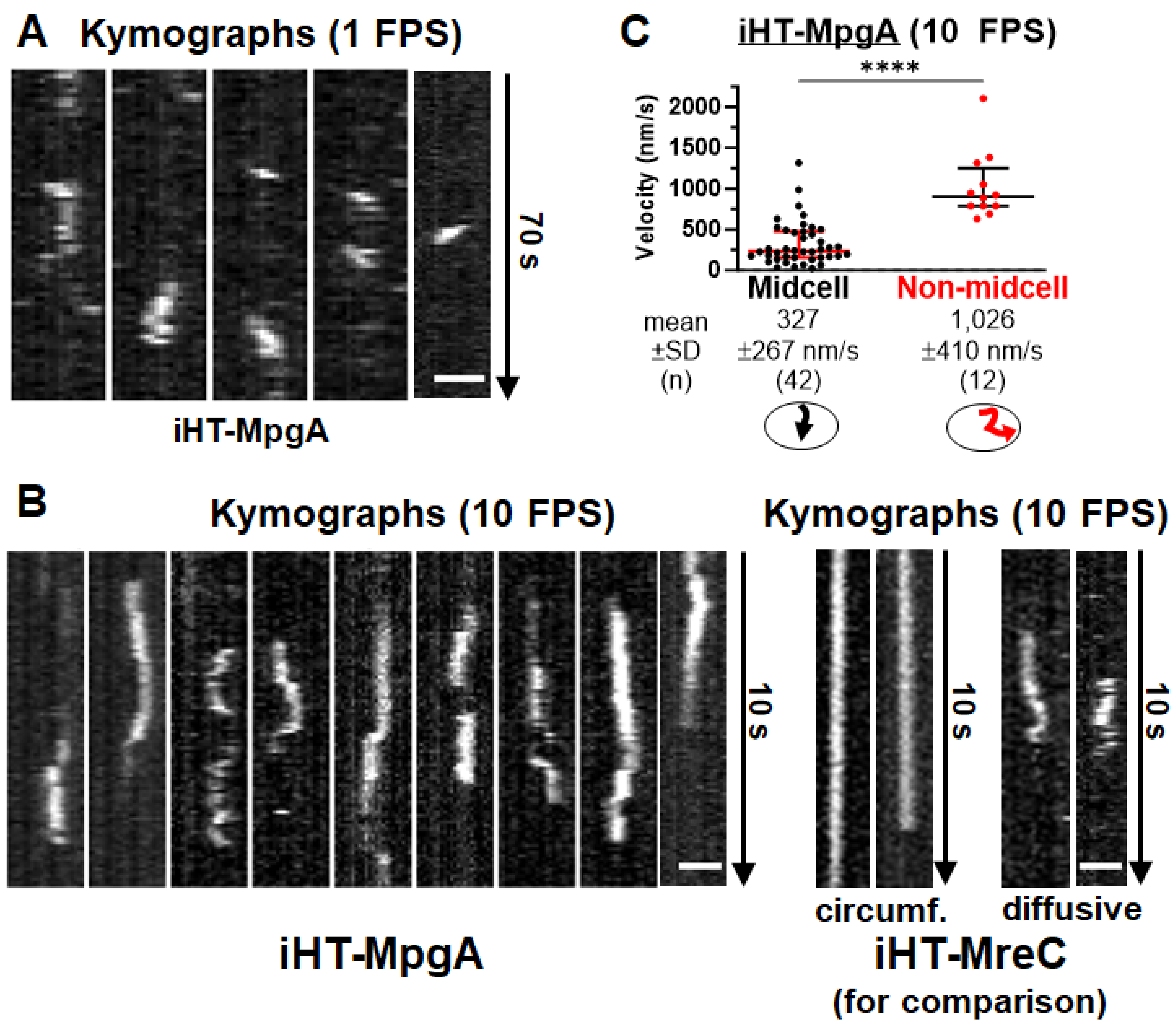
A Kymographs (1 FPS)
70 s
iHT-MpgA
C iHT-MpgA (10 FPS)
****
Velocity (nm/s)
2000
1500
1000
500
0
Midcell
Non-midcell
mean
±SD
(n)
327
±267 nm/s
(42)
1,026
±410 nm/s
(12)
B
Kymographs (10 FPS)
Kymographs (10 FPS)
10 s
10 s
10 s
circumf.
diffusive
iHT-MpgA
iHT-MreC
(for comparison)

Fig. S21

**Fig. S21.** MpgA displayed confined subdiffusive movement at midcell. Sm-TIRFm was performed on strains *iht-mpgA* (IU15997) and *iht-mreC* (IU16344) as described in *Materials and Methods*. Representative kymographs of (*A*) iHT-MpgA molecules imaged at 1 FPS moving in midcell regions. (*B*) iHT-MpgA and iHT-MreC molecules imaged at 10 FPS moving in midcell regions, and iHT-MreC imaged at 10 FPS moving diffusively in non-midcell regions. Kymographs of circumferentially moving iHT-MreC (*middle*) appear vertical due to the increased acquisition rate. (*C*) Dot plots of velocities determined at 10 FPS over short runs of iHT-MpgA molecules in midcell regions (black) and at non-midcell regions (red). Black and red lines are median ± interquartile, and mean ± SD are indicated. n = total molecules analyzed from two biological replicates. Velocities in different regions of cells were compared using a Mann-Whitney test. ****$P < 0.0001$.

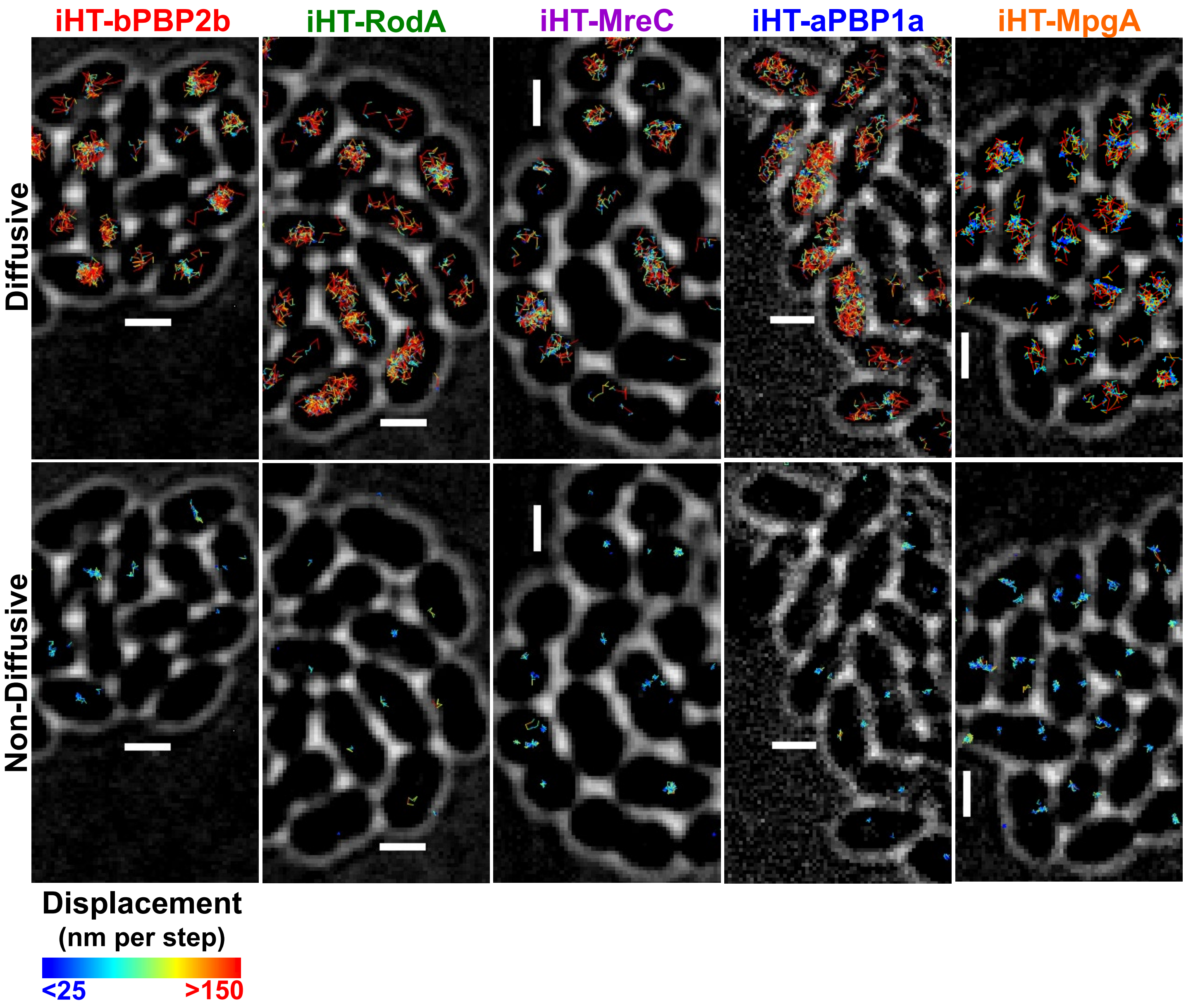
iHT-bPBP2b
iHT-RodA
iHT-MreC
iHT-aPBP1a
iHT-MpgA
Diffusive
Non-Diffusive
Displacement
(nm per step)
<25
>150

Fig. S22

**Fig. S22.** Elongation PG synthesis proteins exhibited different patterns of confined diffusion. Sm-TIRFm was performed at 20 FPS on strains *iht-pbp2b* // $P_{Zn}$-*iht-pbp2b* (IU16553), *iht-rodA* // $P_{Zn}$-*iht-rodA* (IU16496), *iht-mreC* (IU16344), *iht-pbp1a* // $P_{Zn}$-*iht-pbp1a* (IU16497) and *iht-mpgA* (IU15997) as described in *Materials and Methods*. 0.25 mM Zn inducer was added to strains IU16553, IU16496 and IU16497. Representative fields of cells containing multiple single-molecule trajectories (colored lines) were classified as diffusive (top row) and non-diffusive (bottom row). White cell outlines are DIC images. Trajectories with a displacement > 0.13 μm and a velocity standard deviation > 0.63 μm/s were defined as diffusive, and the remaining trajectories (processive or static) were defined as non-diffusive. Full details of trajectory constructions and classification criteria are described in *Materials and Methods*. The color of each trajectory line segment represents the displacement (in nm) of the molecule from one frame to the next. Scale bar = 1 μm.

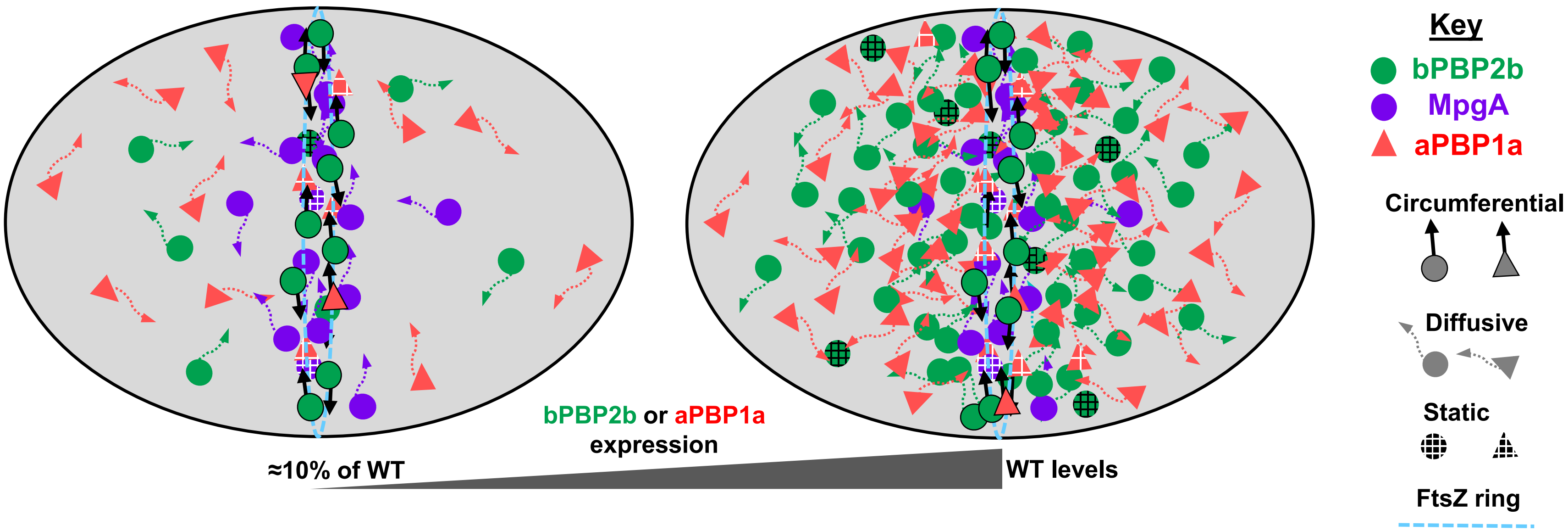
bPBP2b or aPBP1a
expression
≈10% of WT
WT levels
Key
bPBP2b
MpgA
aPBP1a
Circumferential
Diffusive
Static
FtsZ ring

**Fig. S23**

**Fig. S23.** A diagrammatic representation of the motion distribution of core elongasome components (bPBP2b, RodA, and MreC) and proteins linked to PG elongation synthesis (aPBP1a and MpgA) in growing *S. pneumoniae* cells. When underexpressed (left), components of the core PG elongasome are largely confined to PG synthesis at midcell septa and the equators of predivisional daughter cells starting to divide. In contrast, when expressed at WT levels (right), most molecules of components of the PG elongasome (and aPBP1a) diffuse over the cell surface and are not synthesizing PG. Thus, elongasome components and aPBP1a are in excess in growing cells, and only a limited number engage in midcell PG synthesis. In contrast, subdiffusion of MpgA molecules expressed at the WT level is mainly confined to the midcell region by an unknown mechanism. See text for additional details.
